# New interpretable machine learning method for single-cell data reveals correlates of clinical response to cancer immunotherapy

**DOI:** 10.1101/702118

**Authors:** Evan Greene, Greg Finak, Leonard A. D’Amico, Nina Bhardwaj, Candice D. Church, Chihiro Morishima, Nirasha Ramchurren, Janis M. Taube, Paul T. Nghiem, Martin A. Cheever, Steven P. Fling, Raphael Gottardo

## Abstract

High-dimensional single-cell cytometry is routinely used to characterize patient responses to cancer immunotherapy and other treatments. This has produced a wealth of datasets ripe for exploration but whose biological and technical heterogeneity make them difficult to analyze with current tools. We introduce a new interpretable machine learning method for single-cell mass and flow cytometry studies, FAUST, that robustly performs unbiased cell population discovery and annotation. FAUST processes data on a per-sample basis and returns biologically interpretable cell phenotypes that can be compared across studies, making it well-suited for the analysis and integration of complex datasets. We demonstrate how FAUST can be used for candidate biomarker discovery and validation by applying it to a flow cytometry dataset from a Merkel cell carcinoma anti-PD-1 trial and discover new CD4+ and CD8+ effector-memory T cell correlates of outcome co-expressing PD-1, HLA-DR, and CD28. We then use FAUST to validate these correlates in an independent CyTOF dataset from a published metastatic melanoma trial. Importantly, existing state-of-the-art computational discovery approaches as well as prior manual analysis did not detect these or any other statistically significant T cell sub-populations associated with anti-PD-1 treatment in either data set. We further validate our methodology by using FAUST to replicate the discovery of a previously reported myeloid correlate in a different published melanoma trial, and validate the correlate by identifying it *de novo* in two additional independent trials. FAUST’s phenotypic annotations can be used to perform cross-study data integration in the presence of heterogeneous data and diverse immunophenotyping staining panels, enabling hypothesis-driven inference about cell sub-population abundance through a multivariate modeling framework we call <u>P</u>henotypic and <u>F</u>unctional <u>D</u>ifferential <u>A</u>bundance (PFDA). We demonstrate this approach on data from myeloid and T cell panels across multiple trials. Together, these results establish FAUST as a powerful and versatile new approach for unbiased discovery in single-cell cytometry.

## 1 Introduction

Cytometry is used throughout the biological sciences to interrogate the state of an individual’s immune system at the single-cell level. Modern instruments can measure approximately thirty (via fluorescence) or forty (via mass) protein markers per individual cell [1, 2] and increasing throughput can quantify millions of cells per sample. In typical clinical trials, multiple biological samples are measured per subject in longitudinal designs. Consequently, a single clinical trial can produce hundreds of high-dimensional samples that together contain measurements on many millions of cells.

To analyze these data, cell sub-populations of interest must be identified within each sample. The manual process of identifying cell sub-populations is called “gating”. Historically, manual gating has introduced the potential for bias into cytometry data analysis [1, 3]. The choice of gating strategy is one source of bias, since it is fixed in advance and is one of many possible strategies to identify specific cell sub-populations. A different strategy can lead to different gate placements and consequently different cell counts in those sub-populations. More importantly, in modern, high-dimensional panels, bias is now inevitable since manual gating strategies will only identify cell sub-populations deemed important *a-priori* by the investigator. As the number of possible populations grows exponentially with the number of measured protein markers, manual identification cannot be used to perform unbiased discovery and analysis on high-dimensional cytometry data: there are too many combinations of markers to consider.

Researchers have developed numerous computational methods to address these limitations [4, 5]. Such methods [5–8] have helped scientists interrogate the immune system in a variety of clinical settings [9, 10]. Despite successes, computational gating methods continue to face significant challenges when applied to large experimental datasets. Similar to manual gating, methods often require investigators to bound or pre-specify the number of clusters (i.e., cell sub-populations) expected in a sample [6, 11], or to know the relevant clusters in advance [12]. Such information is generally not available. One proposed solution is to partition a dataset into a very large number of clusters in order to capture its main structure [13]. However, as observed in [14], when methods make strong assumptions about the distribution of protein measurements [15, 16], the structure captured by over-partitioning can reflect a method’s assumptions rather than biological signal.

Another challenge for many methods is that biologically equivalent clusters are given arbitrary numeric labels when samples are analyzed independently. In such cases, methods must provide a way to match biologically comparable clusters across samples. One matching approach is to quantify cluster similarities across samples with a user-specified metric [17, 18]. However, as the dimensionality of the data increases, choosing an appropriate metric becomes more difficult due to sparsity [13]. An alternate approach is to concatenate samples together and then cluster the combined data [7, 19, 20]. However, this approach can mask biological signal in the presence of batch effects or large sample-to-sample variation in protein expression. It also introduces the risk that a method will fail to identify small-but-biologically-interesting clusters, since computational limitations can lead authors to recommend sub-sampling cells from each sample before combining the samples for analysis [8].

We have addressed these challenges by creating a new interpretable machine learning method that discovers and annotates cell populations across cytometry experiments named <u>F</u>ull <u>A</u>nnotation <u>U</u>sing <u>S</u>haped-constrained <u>T</u>rees (FAUST, Figure 1). FAUST makes minimal assumptions about the distribution of protein measurements, detects the number of cell sub-populations present in each sample in a data-driven fashion, and provides those sub-populations with phenotypic labels that are standardized across the experiment. FAUST does this by first computing a *depth score* for each marker in each panel that quantifies how consistently a marker separates into sub-populations in each sample’s *annotation forest* – a term we use for the set of all “reasonable” gating strategies in a sample. Next, FAUST selects a subset of high-scoring markers in the panel, and then estimates a standard number of sample-specific annotation thresholds for each selected marker (the number of thresholds estimated for each marker is also data-driven). After this, FAUST grows multiple *discovery forests* for each sample: each tree in the forest is a proposed clustering of the sample, and each leaf of a tree corresponds to a robust, standardized phenotype (relative to the sample’s standardized annotation thresholds). Then, roughly speaking, FAUST conducts a sequence of auctions across these forests (by assigning a “price” to each leaf of each tree in each forest that quantifies the quality of each phenotype), and “purchases” high-quality phenotypes (those with high prices) detected in a sample after each round of the auction. FAUST concludes by constructing a sample-by-phenotype count matrix for the “purchased” phenotypes.

**Figure 1:**
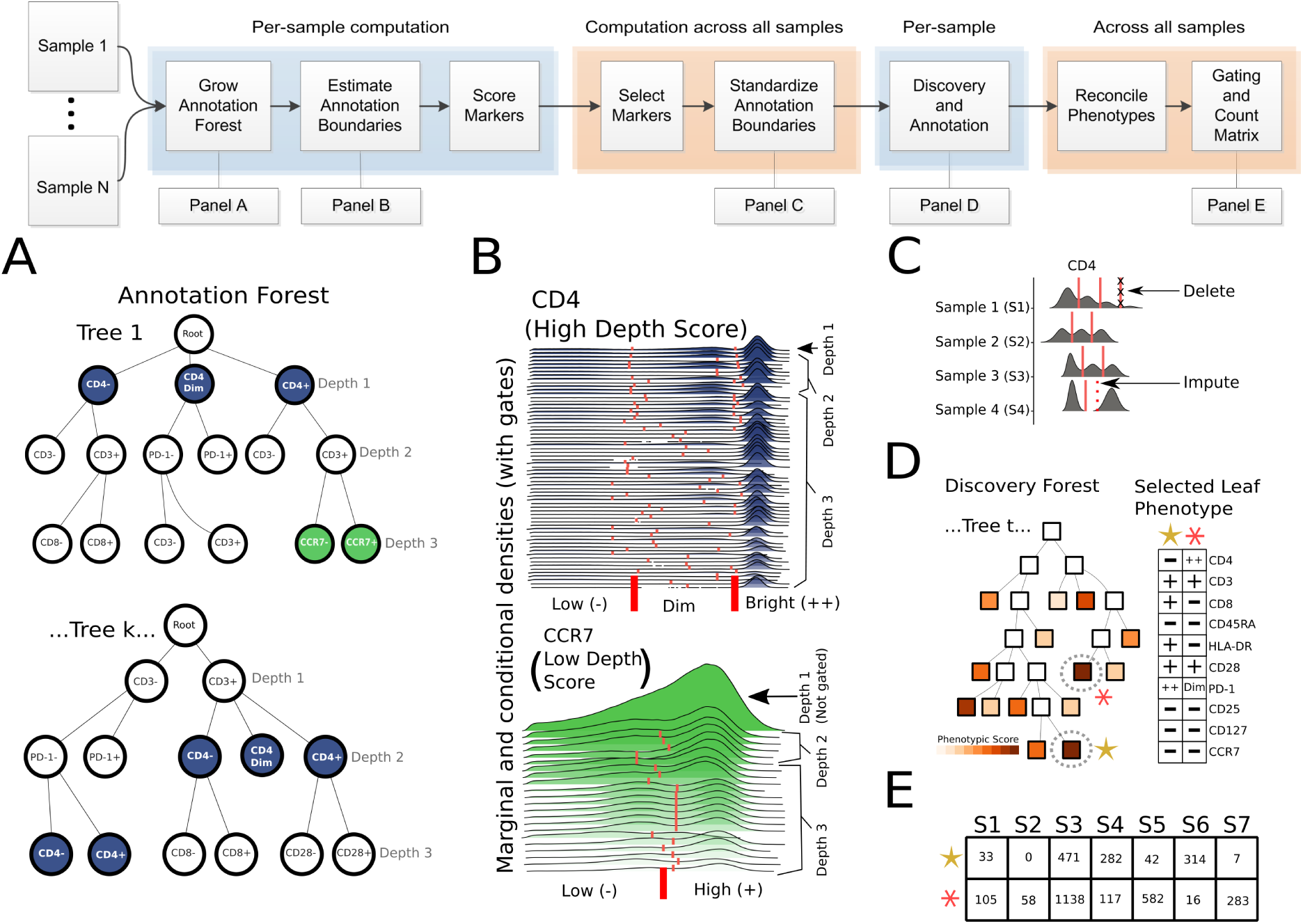
Overview of FAUST. FAUST estimates annotation boundaries for an experimental unit and imputes missing boundaries in a two-stage process. An experimental unit is user–defined and can be a sample, stimulation condition, subject, batch, or site. Imputation is done according to a user-defined imputation-hierarchy and is useful to define classes of samples such as different tissues, batches, time points, or all samples. This schematic overview of FAUST assumes the experimental unit is an individual sample stained with a common panel of markers, while the imputation occurs across all samples in the experiment. A) To estimate annotation boundaries, FAUST grows an exhaustive forest of 1-dimensional, depth-3 gating strategies, constrained by shape: if, prior to depth-3, the cells in a node of the gating strategy have unimodal expression along all markers, the gating strategy along that path terminates. B) Annotation boundaries are estimated for markers within an experimental unit by averaging over gates drawn for that marker over the entire annotation forest. A “depth score” (Methods 4.4) is derived for each marker that quantifies the separation of the marker in each experimental unit. The distribution of scores across experimental units is used to determine whether a marker should be included in the discovery process and to determine the number of annotation boundaries a marker should receive. C) This procedure ensures that FAUST selects a standard set of markers for discovery and annotation as well as a standard number of annotation boundaries per selected marker. D) FAUST then conducts a search to discover and select phenotypes present in each experimental unit. Each discovered phenotype is given a score (described intuitively as a “price” in the main text) that quantifies the homogeneity of cells in an experimental unit with that phenotype; high-scoring phenotypes are then selected for annotation (Methods 4.8). Selected phenotypes are annotated using all selected markers from step C). E) FAUST returns an annotated count matrix with counts of cells in each phenotypic region discovered and selected in step D) that also survives down-selection by frequency of occurrence across experimental units.

By conducting this sequence of auctions, FAUST produces a compact count matrix: it contains counts for far fewer phenotypes than the maximum number of phenotypes implied by the Cartesian product of the annotation thresholds. The count matrix can be made smaller still by adjusting a user parameter that requires a phenotype must be “purchased” in a minimum number of samples before it is included. We also note that since each unique phenotype defines a unique cluster, and since the phenotypes are standardized across samples, phenotypes are comparable across a study by construction. This allows FAUST to process samples independently, which in turn allows FAUST to analyze large datasets without subsampling or concatenating samples. This is an important feature for analyzing studies where there can be substantial differences between biological samples, such as studies with numerous time points, multiple tissue types, or batch-effects.

FAUST facilitates biological discovery and validation since any FAUST-phenotype found to be significantly associated with an outcome of interest in follow-on statistical modeling constitutes a specific hypothesis that can be tested and validated on an independent dataset. We demonstrate this workflow by applying FAUST to data from a Merkel cell carcinoma (MCC) anti-PD-1 trial. In this trial, FAUST identifies several effector memory T cell subsets whose presence in blood at baseline (pre-treatment) is significantly associated with subjects’ positive response to therapy. We then use FAUST to validate these associations by targeting the comparable effector memory T cell phenotypes in baseline anti-PD-1 data from a previously published metastatic melanoma trial, and demonstrate that the same associations hold in the independent dataset.

FAUST also enables targeted hypothesis testing for pre-specified phenotypes in two different ways: a top-down approach and a bottom-up approach. The validation analysis we previously mentioned illustrates the top-down approach: FAUST is used to extract counts for pre-specified phenotypes, without conducting marker selection or discovery and annotation. The bottom-up approach is a multivariate modeling strategy we call “<u>P</u>henotypic and <u>F</u>unctional <u>D</u>ifferential <u>A</u>bundance” (PFDA), inspired by techniques used in gene set enrichment analysis [21]. PFDA fits a multivariate model to all FAUST cell sub-populations whose phenotypes (defined in terms of all markers selected by FAUST) are consistent with a simple, pre-specified phenotype (defined in terms of a subset of the selected markers), and then uses the estimated model coefficients to conduct tests of differential abundance. We contrast these approaches by applying FAUST to myeloid and T cell datasets from three independent cancer immunotherapy trials, and then use both approaches to test hypotheses concerning simple phenotypes. We demonstrate that both approaches offer useful ways to perform cross study analysis, and also show that PFDA can detect signals missed by the top-down approach if the targeted phenotype is mis-specified.

In total, we apply FAUST to six cytometry datasets generated from four independent cancer immunotherapy clinical trials. We demonstrate how FAUST can be used to discover candidate biomarkers associated with treatment outcome, validate these associations on independent data, and perform cross-study analyses in the presence of heterogeneous marker panels.

## 2 Results

### 2.1 FAUST identifies T cell phenotypes in blood, pre-treatment, associated with positive response to anti-PD-1 therapy

We used FAUST to perform cell sub-population discovery in cytometry data generated from fresh, whole blood isolated from patients with Merkel cell carcinoma (MCC) receiving pembrolizumab on the Cancer Immunotherapy Trials Network (CITN) phase 2 clinical trial CITN-09 [22], with the goal of identifying baseline correlates of response to treatment (NCT02267603, see supplementary table S1). We analyzed 78 longitudinal samples stained with immunophenotyping panels to identify T cell subsets within whole blood (Methods 4.10). FAUST selected 10 markers for discovery and subsequently annotated 230 discovered cell sub-populations using these markers, corresponding to 89.6% of cells in the median sample. Of these, 125 had phenotypes that included a “CD3+” annotation. Since the panel was designed to investigate T cells, only these CD3+ sub-populations were used for downstream correlates analysis.

In order to determine if FAUST discovered phenotypes significantly associated with response to therapy, we used binomial generalized linear mixed models (GLMMs, see [23]) to test each sub-population for differential abundance at the baseline time point (prior to receiving anti-PD-1 therapy), between responders and non-responders in 27 subjects (equation (4.5) specifies the model). *Responders* were defined as subjects that exhibited either a complete (CR) or partial (PR) response (per RECIST1.1 [24]), and *non-responders* as subjects exhibiting progressive (PD) or stable (SD) disease. At a Bonferroni-adjusted 10% level (125 tests), four sub-populations were associated with response to therapy (Figure 2A, Figure 3A). Two sub-populations had a CD28+ HLA-DR+ CD8+ effector memory phenotype, with either PD-1 dim (Bonferroni-adjusted p-value: 0.021) or PD-1 bright (Bonferroni-adjusted p-value: 0.051), respectively. The other two were CD4 bright: one had an HLA-DR-CD28+ PD-1 dim phenotype (Bonferroni-adjusted p-value: 0.011); the other an HLA-DR+ CD28+ PD-1 dim (Bonferroni-adjusted p-value: 0.086). The observed CD28+ phenotypes agree with published findings highlighting the importance of CD28 expression in CD8+ T cells in anti-PD1 immunotherapy [25, 26]. Effect sizes with 95% confidence intervals for the correlates are reported in Supplementary Table A.6. All four correlates were annotated CD45RA- and CCR7-, indicating they represented effector-memory T cells. The complete phenotypes are stated in depth-score order (Methods 4.4) in Figure 3.

**Figure 2:**
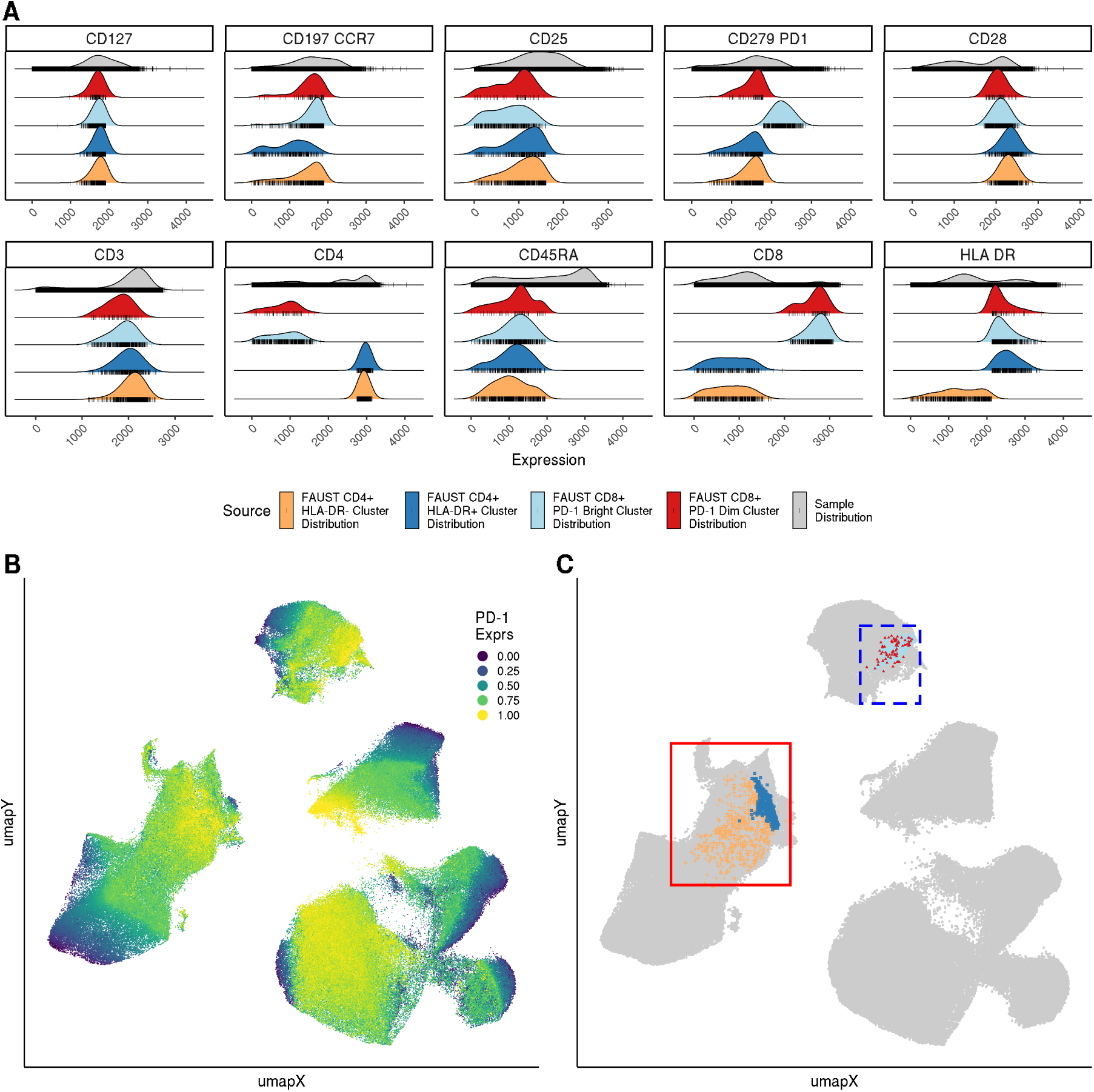
FAUST annotations reflect underlying differences in protein expression not captured by dimensionality reduction. A) In a baseline responder’s sample, the densities of per-marker fluorescence intensity for cells in the four correlates (different colors) as well as the entire non-negative collection of live lymphocytes in the sample (gray) are compared. Cells used in density calculations are marked by tick marks and demonstrate that differences in cluster annotations reflect strict expression differences in the underlying data. B) A UMAP embedding computed from the same sample as panel A using the ten stated protein markers. All cells in the sample were used to compute the embedding. The embedding is colored by the relative intensity of observed PD-1 expression, windsorized at the 1st and 99th percentile, and scaled to the unit interval. The sample contains 271,219 cells, of which 72 were CD8+ PD-1 dim cells, 599 were CD8+ PD-1 bright cells, 673 were CD4 bright HLA-DR-CD28+ PD-1 dim cells, and 376 were CD4 bright HLA-DR+ CD28+ PD-1 dim cells. C) The same UMAP embedding highlighting the location of the cells from the four discovered sub-populations. The HLA-DR+ CD28+ PD-1 dim/bright effector memory CD8 T cells are in the dashed blue box; the HLA-DR+/- CD28+ PD-1 dim CD4 T cells, the solid red box. Exact phenotypes of the four sub-populations can be determined by matching the cluster color to panel A, and are also reported in figure 3.

**Figure 3:**
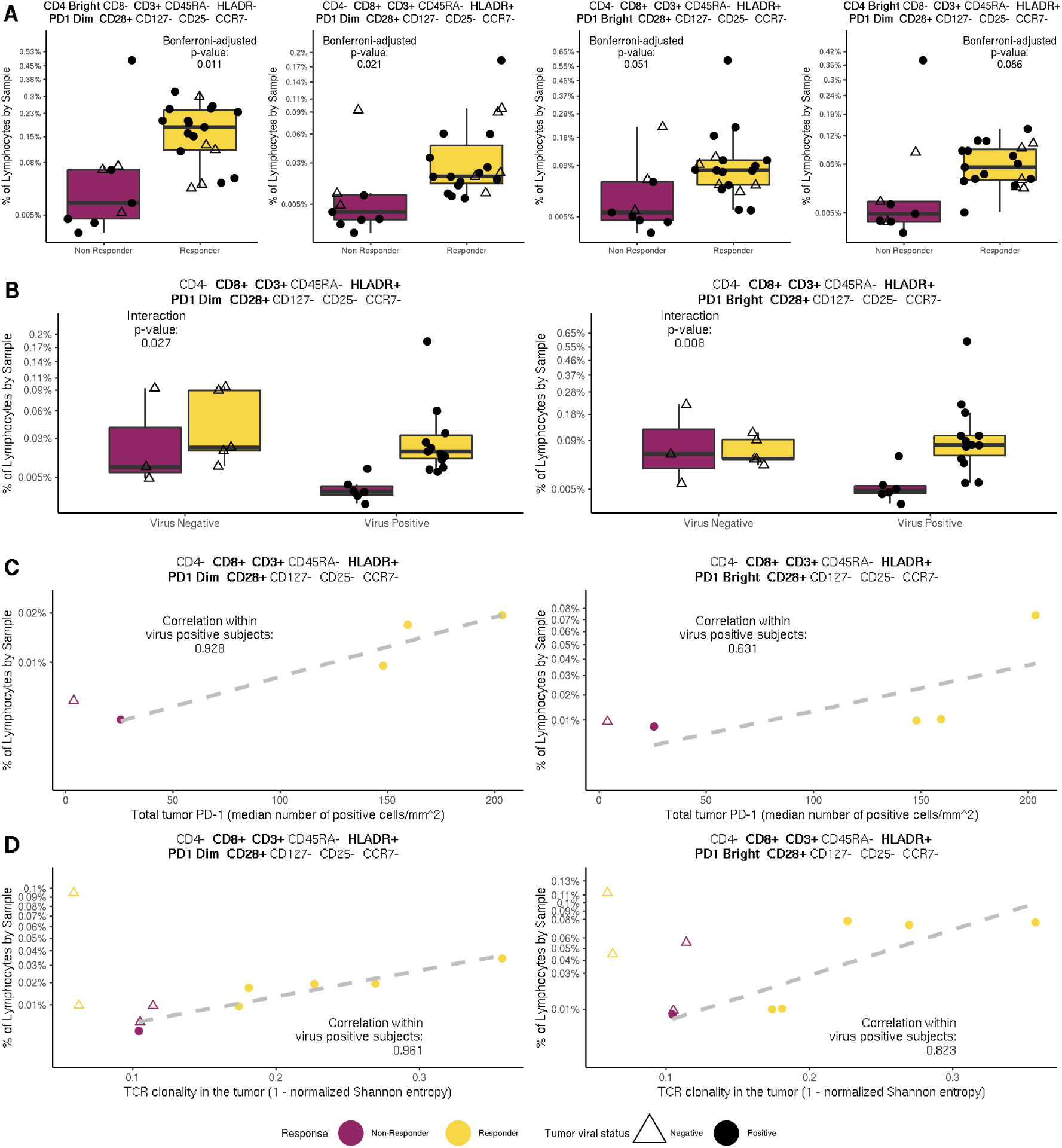
Increased abundance of effector memory CD4/CD8 T cells co-expressing CD28, HLA-DR, and PD-1 is associated with positive response to anti-PD-1 therapy. A) Boxplots of the abundance of the four T cell sub-populations – with FAUST annotations reported at the top of each plot – associated with positive treatement outcome discovered by FAUST, stratified by (subjects’ response to therapy. Bonferroni adjusted p-values reported contrasting all responders (n = 18) against all non-responders (n = 9). B) Boxplots of the abundance of the two CD8 T cell sub-populations stratified by their viral status, with interaction p-value reported. The two CD4+ T cell sub-populations are not presented because we did not observe a significant interaction between a subject’s response to therapy and their viral status. C) The abundance of the two CD8+ T cell correlates among virus positive subjects against total PD-1 expression measured by IHC from tumor biopsies as described in [30], with observed correlation in virus positive subjects (n=4) reported. D) The abundance of the two CD8+ T cell correlates among virus positive subjects plotted against productive clonality (1-normalized entropy) from tumor samples as described in [31], with observed correlation in virus positive subjects (n=6) reported.

We inspected the primary flow cytometry data in order to confirm that the discovered phenotypes were consistent with the underlying protein expression. By plotting cluster densities against samples (Figure 2A), we observed that the FAUST annotations accurately described the observed cellular phenotypes in these sub-populations. We also visualized these data using UMAP embeddings [27] with “qualitative” parameter settings [28] (Figure 2B,C). We observed that FAUST clusters were not typically separated into isolated “islands” in the UMAP embedding (Figure 2C), and that single UMAP “islands” contained significant variation in expression of some of the measured protein markers (Figure 2B). Taken together, these observations demonstrate that visualizations derived via dimensionality reduction (here, UMAP) do not necessarily reflect all variation measured in the underlying protein data, and that any method that solely relies on UMAP for population discovery would likely miss these sub-populations.

MCC is a viral-associated malignancy: Merkel cell polyomavirus occurs in many MCC tumors [22]. Reports that CD8 T cells co-expressing HLA-DR and CD28 can exhibit anti-viral properties [29], as well as reports of CD28 dependent rescue of exhausted CD8 T cells by anti-PD1 therapies in mice [26], led us to investigate the association between the abundance of the therapeutic-response-associated sub-populations discovered by FAUST and tumor viral status of each subject, as we hypothesized that these cells may represent virus specific sub-populations. We adapted the differential abundance GLMM to test for an interaction between response to therapy and tumor viral status in the four significant FAUST phenotypes (Figure 3A). This interaction was statistically significant (PD-1 dim *p* = 0.027; PD-1 bright *p* = 0.008) for both CD8+ correlates (Figure 3B). The interaction was not significant for the two CD4+ correlates (data not shown). This suggested that the CD8+ T cells may be particularly relevant to anti-tumor response in subjects with virus-positive tumors.

In order to further investigate the relevance of these CD8+ T cells, we examined published data on PD-1 immunohistochemistry (IHC) staining in tumor biopsies from the same patients (described in [30]). Importantly, the in-tumor PD-1 measurement is a known outcome correlate in MCC [30]. Limited overlap between the assays resulted in only five subjects where both flow cytometry and tumor biopsy anti-PD-1 IHC staining were available, and only four of these were virus-positive. Nonetheless, the frequencies of the CD8+ T cells were strongly correlated (PD-1 dim *ρ* = 0.928; PD-1 Bright *ρ* = 0.631) with the PD-1 total IHC measurements within the four virus-positive subjects (Figure 3C). We also examined published TCR clonality data generated from patient tumor samples, described in [31]. Ten subjects passing clonality QC were common to the two datasets, six of which were virus positive. Frequencies of the CD8+ FAUST populations within these six subjects were strongly correlated (PD-1 dim *ρ* = 0.961; PD-1 Bright *ρ* = 0.823) with the measurement of productive clonality (Figure 3D).

Together, these results lend support to the hypothesis that the CD8+ T cell correlates discovered by FAUST in blood represents a circulating population of tumor-associated virus-specific T cells that are also detectable in the tumor and whose presence in the tumor is known to correlate with outcome. Due to the small sample size, this hypothesis must be confirmed on an independent, larger set of MCC patient samples. However, since all four reported cell subsets (Figure 3A) were detected without incorporating the viral status of each subject into the statistical model, these results also suggested more general hypotheses: increased abundance of the CD4+ or CD8+ sub-populations discovered by FAUST is associated with positive response to pembrolizumab therapy. We investigate these hypotheses in section 2.2.

### 2.2 FAUST T cell correlates of anti-PD-1 therapy are validated on independent dataset

In order to validate our findings in the MCC study, we tested the hypotheses that increased abundance of the T cell correlates discovered by FAUST in the MCC trial are associated with positive response to pembrolizumab treatment by first downloading a published metastatic melanoma CyTOF dataset from FlowRepository [32]. A full description of the dataset is given by Subrahmanyam et al. in [33]. Here, we restricted our analysis to unstimulated baseline PBMC samples.

After downloading the dataset, we used the computational package openCyto [34] to replicate the pre-gating strategy reported in Figure 1 of [33] (Supplement A.7). The forty marker CyTOF panel used in [33] contained all ten markers used by FAUST to define T cell correlates in the MCC study. To target the MCC correlates in this new study, we down-selected the markers in the CyTOF panel (after pre-gating live intact singlets) to match the annotating markers from the MCC trial. We then used FAUST to estimate data-driven annotation thresholds for the ten matching markers. We note that this type of targeting is an example of the “top-down” method of FAUST analysis where we use FAUST’s annotation thresholds to extract counts for phenotypes of interest (see section 2.5).

Next, we used these thresholds to extract counts of the pre-specified T cell correlates in the CyTOF melanoma dataset. In the CyTOF melanoma study, FAUST defined one standardized threshold for all markers (including PD-1 and CD4). In the MCC study, FAUST defined two standardized thresholds for PD-1 and CD4, and one threshold for all other markers. We therefore mapped “PD-1+” cells in CyTOF melanoma dataset to cells in the MCC dataset with the consolidated phenotype “PD-1 dim/bright”. We used a similar map across datasets for CD4, leading us to extract per-sample counts (relative to the sample thresholds) for three phenotypes in the CyTOF melanoma study (full phenotypes are stated in Figure 4).

**Figure 4:**
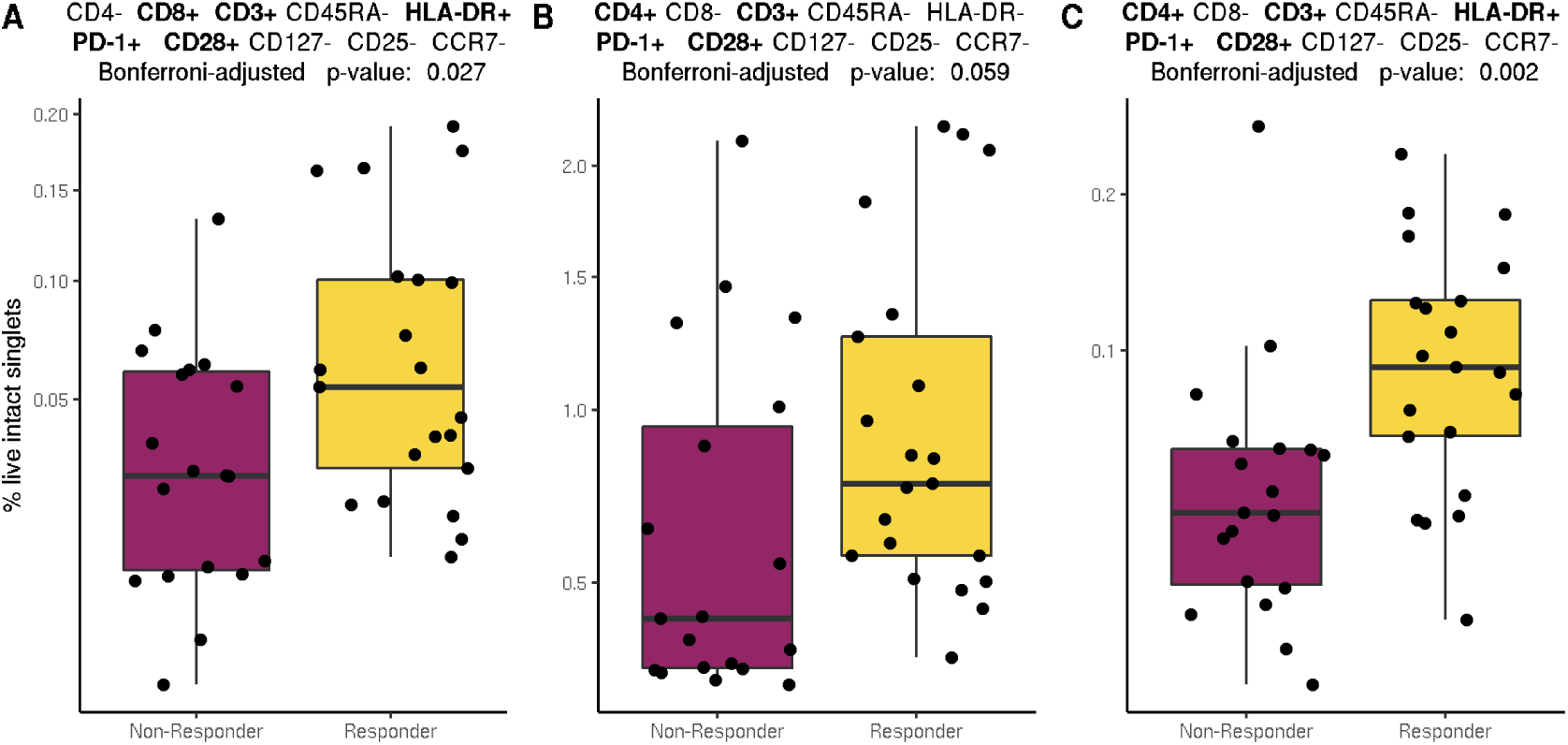
Significant FAUST phenotypes are validated in an independent melanoma trial. A) Boxplots of the abundance of effector memory CD8 T cells co-expressing CD28, HLA-DR, and PD-1 in unstimulated baseline samples for subjects that were treated with pembrolizumab, stratified by responder status, with Bonferroni-adjusted p-value reported (3 tests). B) The same as A, but displaying effector memory CD4 T cells co-expressing PD-1 and CD28. C) The same as A, but displaying effector memory CD4 T cells co-expressing PD-1, HLA-DR, and CD28. All sub-populations were targeted by first using FAUST to compute annotation boundaries for the listed markers within live intact singlets for each sample, and then using those boundaries to determine counts of cells with the stated phenotypes. All phenotypes matched the FAUST phenotypes from the MCC trial exactly, up to the number of annotation boundaries determined for each marker.

To validate the associations, we then tested the three phenotypes for differential abundance between responders and non-responders in samples from subjects that went on to receive pembrolizumab (the same anti-PD-1 therapy used in the MCC trial). We used the same binomial GLMM we used in the MCC trial to conduct the test for differential abundance between responders and non-responders (equation (4.5)), applied the Bonferroni adjustment for three tests, and detected significantly increased abundance of all three tested T cell phenotypes in responding subjects (Figure 4A-C). We note that in the CyTOF melanoma dataset, responders were defined as patients that exhibited progression-free survival for at least 180 days after therapy [33]. This analysis validated the associations detected by FAUST in the MCC trial, and highlights a powerful feature of FAUST: performing targeted validation across studies and technologies.

### 2.3 FAUST sub-populations capture underlying biological and technical signals in longitudinal studies

Consistently identifying and annotating cell populations that are missing across a subset of samples is a significant challenge in computational cytometry analysis [35]. To demonstrate how FAUST’s phenotypic standardization can address this issue we examined the longitudinal profiles of specific cell sub-populations in the MCC anti-PD-1 trial for which we expected longitudinal changes in the abundance due to known technical effects. In the MCC anti-PD-1 trial, we examined all CD8+ T cells with the PD-1-bright FAUST phenotype. The temporal abundance of these cells is shown in (Figure 5A) and reveals that these cells are not detectable in most samples after subjects have received pembrolizumab therapy, presumably from pembrolizumab blocking the detecting antibody. The observed decay post-treatment is consistent with the manual gating of CD3+ CD8+ PD-1+ cells in this study (Supplementary Figure S1).

**Figure 5:**
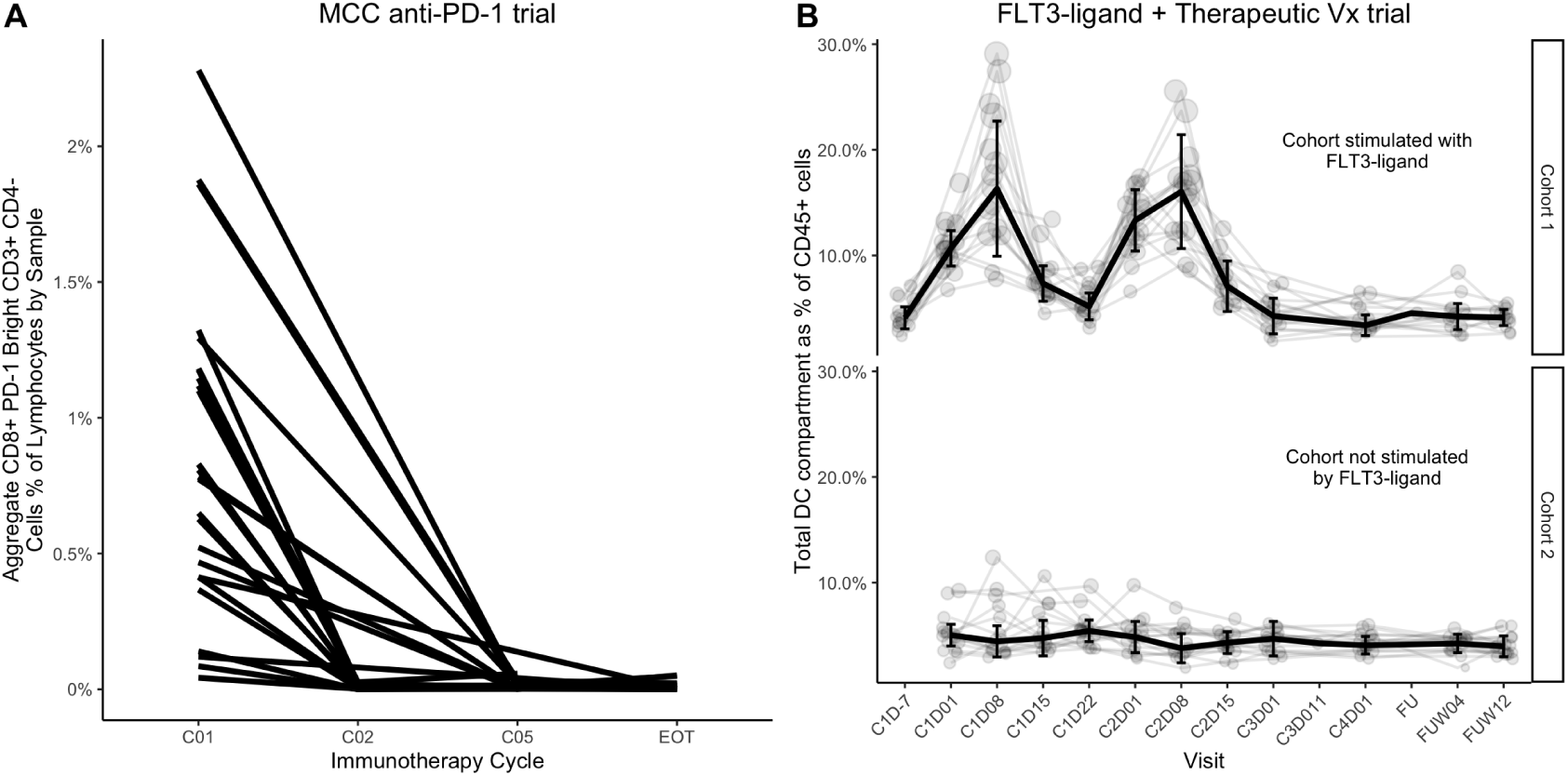
The longitudinal profiles of aggregated FAUST cell populations in a pembrolizumab therapy trial and a FLT3-L + CDX-1401 trial. A) The aggregated frequency of all CD8+ PD-1-bright T-cell populations found by FAUST across all time points. B) The longitudinal profiles of all cell sub-populations with phenotypes consistent with the DC compartment: CD19-, CD3-, CD56-, HLA-DR+, CD14- CD16- and CD11C+/-. Light colored lines show individual subjects. The dark line shows the median across subjects over time. Error bars show the 95% confidence intervals of median estimate at each time point. Cohort 1 (n=16 subjects), cohort 2 (n=16 subjects).

We also analyzed flow cytometry data from a second CITN trial: CITN-07 (NCT02129075, see supplementary table S1for trial data), a randomized phase II trial studying immune responses against a DEC-205/NY-ESO-1 fusion protein (CDX-1401) and a neoantigen-based melanoma vaccine plus poly-ICLC when delivered with or without recombinant FLT3 ligand (CDX-301) in treating patient with stage IIB to stage IV melanoma. The cytometry data consisted of fresh whole blood stained for myeloid cell phenotyping (Methods 4.12). Here, FAUST discovered and annotated 128 cell sub-populations using 10 markers (selected by depth-score), assigning phenotypic labels to 92.9% of cells in the median sample.

In the FLT3-Ligand + therapeutic Vx trial we expected to observe expansion of dendritic cells in response to FLT3-L stimulation [36]. Examination of the longitudinal profile of clusters with phenotypic annotations consistent with dendritic cells (Figure 5B) revealed dynamic expansion and contraction of the total DC compartment in the FLT3-L stimulated cohort but not in the unstimulated-by-FLT3-L-pre-treatment cohort. The expansion peaked at day 8 after FLT3-L simulation in cycles 1 and 2. This dynamic is consistent with observations from manual gating of the DC population [37], the expected biological effect of FLT3-L [36], and the timing of FLT3 administration.

These results demonstrate that FAUST is able to detect, annotate, and correctly assign abundance to cell sub-populations, even those sub-populations that are missing in some samples. Additionally, they demonstrate that FAUST’s phenotypes provide a unique way to validate cluster quality in the absence of human labels: FAUST phenotypes can be checked for biological responses expected from the experimental design. The longitudinal behavior of PD-1 bright T cell populations in the MCC anti-PD-1 trial and the dendritic cells in the FLT3 ligand + CDX-1401 trial are consistent with manual gating of cytometry data and serve as an internal validation of the methodology.

### 2.4 FAUST robustly detects CD14+ signal across multiple immunotherapy trials

Both the MCC anti-PD-1 and FLT3-L + therapeutic Vx trials had cytometry data stained with a myeloid phenotyping panel. We also selected two myeloid phenotyping datasets (one CyTOF discovery and one FACS validation assay) from a previously-published anti-PD-1 trial in metastatic melanoma [9]. We will refer to these as the melanoma anti-PD-1 FACS and melanoma anti-PD-1 CyTOF datasets. We used FAUST to conduct unbiased discovery on these four datasets. A principal finding of the published analysis of the melanoma anti-PD-1 trial was that the frequency of CD14+ CD16-HLA-DR^hi^ cells was associated with positive response to therapy. We wished to determine if FAUST discovered phenotypes significantly associated with outcome that are consistent with the published results across these four independent datasets.

In all datasets, we found that FAUST identified significant baseline phenotypes associated with clinical outcome at the Bonferroni-adjusted 10% level that were consistent with the previously-published CD14+ CD16-HLA-DR^hi^ phenotype (Figure 6). Complete phenotypes, effect sizes and confidence intervals for all myeloid baseline predictors discovered in the MCC anti-PD-1 myeloid phenotyping data are in Supplementary Table S2; those discovered in the FLT3-L + therapeutic Vx trial are in Supplementary Table S3. All statistical models are fully described in section 4.

**Figure 6:**
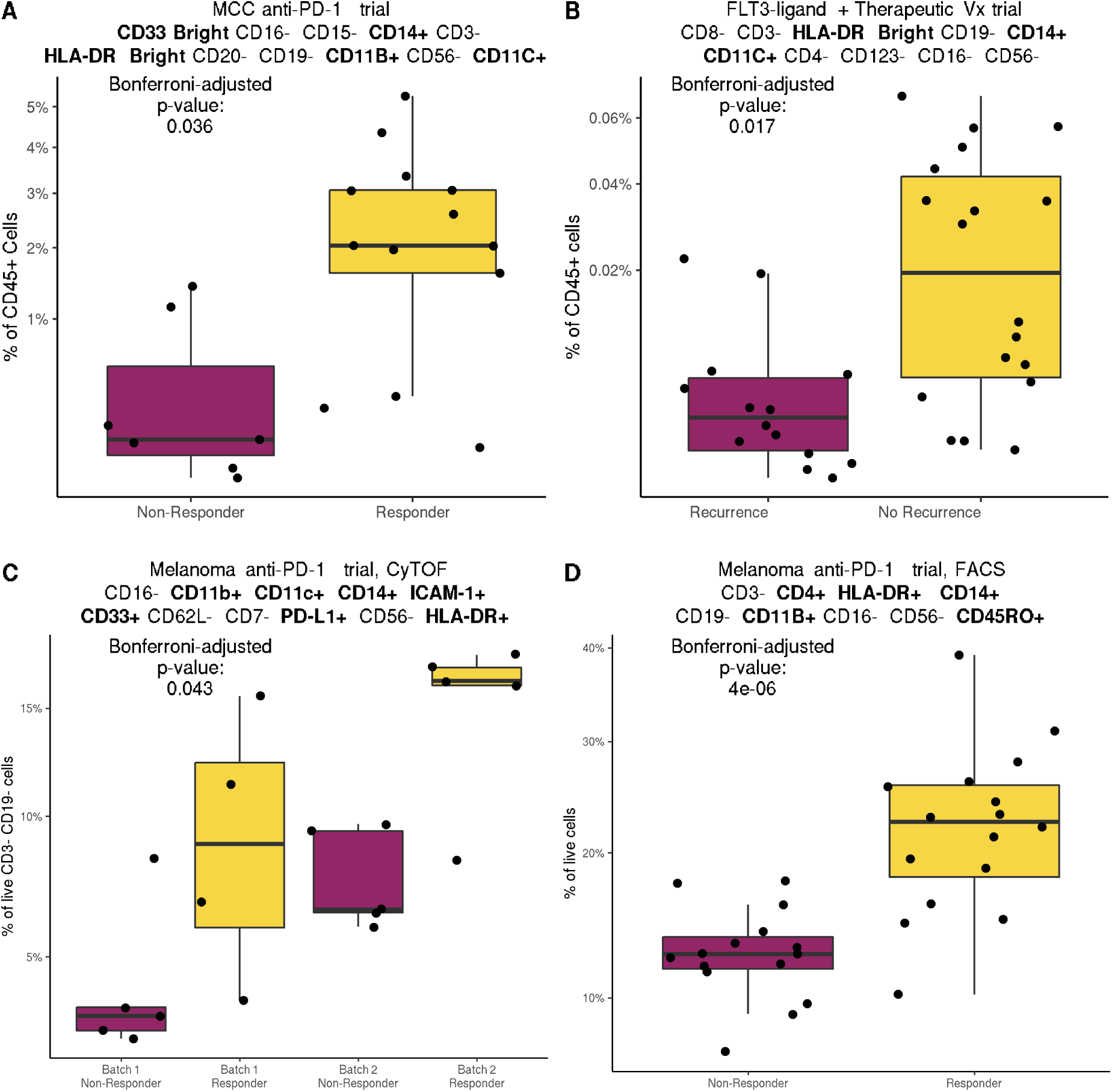
FAUST consistently discovers CD14+ CD16- HLA-DR+ phenotypes associated with outcome at baseline across immunotherapy trials. A) The baseline outcome-associated sub-population discovered by FAUST in the MCC anti-PD-1 trial myeloid data (n=20, 13 Responders, 7 Non-Responders). B) The baseline outcome-associated sub-population discovered by FAUST in the FLT3-L therapeutic Vx trial myeloid data (n=32, 18 No Recurrence, 14 Recurrence). C) The baseline outcome-associated sub-population found by FAUST from the re-analysis of the Krieg CyTOF panel 03 (stratified by batch) (n=19, 9 Responder, 10 Non-Responder). D) The baseline outcome-associated sub-population found by FAUST from the re-analysis of the Krieg FACS validation data (n=31, 16 Responder, 15 Non-Responder). Full FAUST phenotypes are reported above each boxplot.

These results demonstrate the power of our approach to detect candidate biomarkers in a robust manner across different platforms, staining panels, and experimental designs. They also highlight that the complete phenotypes discovered by FAUST may not be directly comparable across datasets due to differences in the marker panels. However, the sub-phenotype CD14+ CD16- HLA-DR+/bright CD3- CD19- CD56- is contained within the marker panels of the three flow cytometry datasets, naturally raising the question: is this sub-phenotype differentially abundant across the datasets? This desire to test specific hypotheses across datasets led us to develop two analysis methods which we describe in section 2.5.

### 2.5 FAUST enables targeted hypothesis testing for pre-specified phenotypes

In this section, we describe two methods that use FAUST to test pre-specified hypotheses. We will informally refer to these methods as the top-down approach and bottom-up approach. The top-down approach is where we use FAUST to extract cell sub-population phenotypes in a targeted manner rather than perform unbiased discovery. This is the method used in section 2.2 (where we validated the T cell correlates discovered in the MCC study of section 2.1), and is the method we recommend using when one wants to test a pre-specified phenotype. This approach consists of the following steps. First, use FAUST to estimate annotation thresholds for pre-specified markers across a cytometry experiment. Then, extract counts for pre-specified phenotypes relative to the estimated thresholds. Finally, test the targeted phenotypes for association with outcome.

The bottom-up approach is an alternative mechanism to interrogate a phenotype-of-interest, that provides some protection against the possibility of phenotype mis-specification. For example, in the MCC trial, we may expect a signal in PD-1+ T cells without knowing which specific sub-phenotypes are associated with outcome. The bottom-up approach was developed to take advantage of the full annotations produced by FAUST – drawing inspiration from the techniques used to perform gene set enrichment analysis in gene expression data [21] – and it works as follows. Suppose we wish to test a phenotype that can be described using only four of ten markers selected by FAUST for differential abundance between responders and non-responders. We fit a multivariate model to all ten-marker FAUST cell sub-populations that contain the four-marker phenotype within their annotations, and then conduct tests of differential abundance using linear combinations of the estimated model coefficients. This approach accounts for the correlation between the different cell sub-populations discovered by FAUST, as well as the different patterns of abundance observed in each of those sub-population. We call this approach “<u>P</u>henotypic and <u>F</u>unctional <u>D</u>ifferential <u>A</u>bundance” (PFDA).

To contrast these two approaches, we apply each approach to two hypotheses. The first hypothesis is based on the published findings of Krieg et al. [9] as well as the unbiased FAUST analysis described in section 2.4: we hypothesize that CD14+ CD16- HLA-DR+ CD3- CD56- CD19- cells exhibit increased abundance in responding subjects at baseline (Figure 7A). The second hypothesis – which we alluded to above – arises from the fact that baseline samples collected in the MCC study (section 2.1) are from subjects that go on to received anti-PD-1 therapy: we hypothesize that total CD4 and total CD8 T cells expressing PD-1 are elevated in responders at baseline.

**Figure 7:**
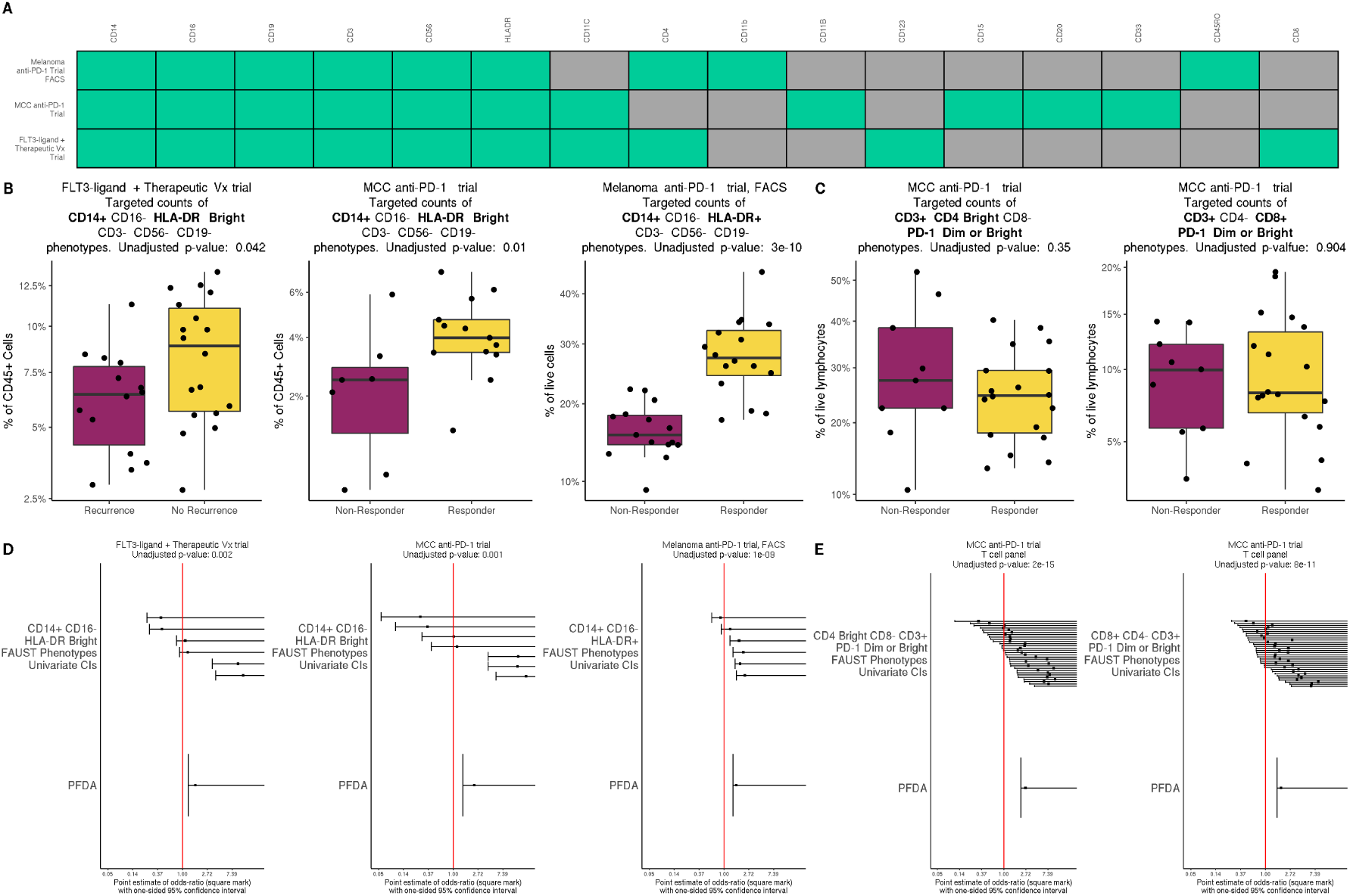
FAUST phenotypes enable cross-study meta-analysis of datasets stained with disparate marker panels. A) Myeloid phenotyping marker panels from three trials. A marker with a green box is present in the stated trial; a marker with a gray box, absent. B) Box plots displaying targeted counts for CD14+ CD16- HLA-DR+/bright CD3- CD56- CD19- phenotypes across three trials. C) Box plots displaying targeted counts for CD3+ CD4 Bright CD8- PD-1 dim or bright and CD3+ CD4- CD8+ PD-1 dim or bright phenotypes in the MCC trial T cell panel. D) Forest plots displaying one-sided 95% confidence intervals (CIs) for increased abundance of CD14+ CD16- HLA-DR+/bright CD3- CD56- CD19- phenotypes in responders vs. non-responders for three trials. In each panel, we first show CIs derived from fitting the univariate model to each CD14+ CD16- HLA-DR FAUST phenotype. We then show the 95% CI derived by fitting a “ PFDA” model jointly to the FAUST populations, and then testing for increased abundance using model coefficients. E) Similar forest plots to those described in D), for CD3+ CD4 Bright CD8- PD-1 dim or bright and CD3+ CD4- CD8+ PD-1 dim or bright phenotypes in the MCC trial T cell panel. Unadjusted p-values are also reported in panels B,C,D,E.

To test the first hypothesis with the top-down approach, we used FAUST to estimate sample-specific annotation thresholds across the three flow-cytometry datasets, extracted counts of CD14+ CD16- HLA-DR+/bright CD3- CD56- CD19- cells in each study (Figure 7B), and then tested for differential abundance using the previously described binomial GLMMs. We excluded the melanoma anti-PD-1 CyTOF dataset from this demonstration, since 10 of 19 baseline samples had fewer than 1500 total cells. We observed significantly increased abundance among responding subjects in each trial. To test the second hypothesis with the top-down approach, we used FAUST to estimate sample-specific annotation thresholds for the samples stained by the T cell panel in the MCC study, extracted counts for cells with CD4 bright CD3+ CD8- PD-1 bright and total CD8+ CD3+ CD4- PD-1 bright phenotypes, and then tested for differential abundance using the previously described binomial GLMMs. In this case, we did not observe a significant increase in abundance among responding subjects for either phenotype (Figure 7C).

Next, we repeat these analyses using PFDA. To test the first hypothesis with PFDA, we fit a multivariate model to the FAUST phenotypes containing the annotation CD14+ CD16- HLA-DR+/bright CD3- CD56- CD19- in each trial, and then tested for increased abundance using the estimated model coefficients (Methods 4.16). In the FLT3-ligand + Therapuetic Vx trial, 6 FAUST phenotypes contained this annotation; in the MCC anti-PD-1 trial, 7 FAUST phenotypes; in the Melanoma anti-PD-1 trial FACS dataset, 6 FAUST phenotypes (Figure 7D). To test the second hypothesis with PFDA, we fit a multivariate models to the 30 CD4 bright and 32 CD8+ respective FAUST phenotypes in the MCC anti-PD-1 T cell dataset, and tested for increased abundance using each models coefficients (Figure 7E). When using PFDA, we observed significantly increased abundance of all phenotypes in responders across datasets (Figure 7D,E).

The total CD4 and total CD8 T cell results demonstrate that the top-down approach does not always detect differential abundance when used to test phenotypes that contain numerous sub-populations with weak effects (Figure 7C,E). On the other hand, PFDA can detect differential abundance in these cases by fitting a joint model to all sub-populations discovered by FAUST within the phenotype-of-interest (so accounting for their correlation structure), and then using the model coefficients to test for an overall increase in abundance (Figure 7E). In the case of the CD14+ CD16- HLA-DR+/bright CD3- CD56- CD19- results, there is less heterogeneity detected within the phenotype-of-interest across the studies, and both the top-down approach and PFDA detect increased abundance in responding subjects (Figure 7B,D). Together, these results demonstrate two ways that FAUST enables targeted hypothesis testing of pre-specified phenotypes, which in turn makes it possible to use FAUST to carry out validation analyses as well as to integrate findings across studies through PFDA.

### 2.6 FAUST outperforms state-of-the-art computational cell population discovery methods for identification of T cell outcome correlates

We re-analyzed the MCC anti-PD-1 T cell dataset described in section 2.1 with the methods *FlowSOM* [6], *Phenograph* [38], and FAUST. We had two goals in re-analyzing this dataset. Our first goal was to test how robust FAUST’s findings were to different data transformations and different selection of samples. Our second goal was to test if other methods detected significant correlates with anti-PD-1 therapy in baseline samples.

In this re-analysis, we ran each method on live lymphocytes from all 78 experimental samples as well as from the 27 baseline samples alone. To test performance under different data transformations, we transformed samples using the biexponential map (used for analysis in section 2.1) as well as inverse hyperbolic sine with co-factor 120 (the transformation reported in [5]). For all methods, we tested clusters for differential abundance between responders and non-responders in baseline samples using the same binomial GLMM employed in section 2.1 (equation (4.5)). For FlowSOM and Phenograph, samples were concatenated before clustering, and we set tuning parameters to the settings reported in [5]. After testing for differential abundance between responders and non-responders, no clusters defined by FlowSOM or Phenograph were associated with response to therapy at the Bonferroni-adjusted 20% level or the FDR-adjusted 20% level (TableS5). On the other hand, across all re-analysis settings FAUST defined an effector memory CD8+ T cell cluster co-expressing CD28, HLA-DR, and PD-1, as well as an effector memory CD4+ T cell cluster co-expressing CD28 and PD-1, and both were associated with response to therapy at baseline at the Bonferroni-adjusted 10% level across all re-analysis settings (TableS5).

We also conducted an *in silico* study that simulated the discovery process in cytometry data analysis by inducing a differentially abundant population associated with a simulated response to therapy, in datasets generated from a variety of mixture models (Supplementary Section A.13). We compared FAUST to FlowSOM in this study since FlowSOM is computationally efficient and is recommended in the review [5]. Across simulation settings, we found that FAUST consistently discovered the phenotype whose increased abundance in a subject was causally linked (by simulation) to an increased probability of response to therapy, when the simulated probability of response to therapy met or exceeded 65%. FlowSOM’s discovery performance was similar to FAUST’s in the multivariate Gaussian setting, but was adversely affected by departures from normality, as well as by simulated batch effects combined with nuisance variables. This study confirms our empirical finding that FAUST robustly detects signals in data that are not found by other discovery methods.

## 3 Discussion

We applied FAUST to six cytometry datasets (CyTOF and flow) from four independent immunotherapy trials. Across three trials, FAUST discovered cell sub-populations and labeled them with annotations that are generally consistent with previous manual gating (when aggregated by appropriate annotation), as well as with the known biological context. In the fourth trial (the melanoma trial of Subrahmanyam et al. [33]) we used FAUST to target specific cellular phenotypes – effector memory CD8+ cell subsets co-expressing PD-1, HLA-DR, and CD28, as well effector memory CD4+ subsets co-expressing PD-1, HLA-DR, and CD28 – to validate that increased abundance of these cells was associated with positive response to anti-PD-1 therapy.

We found that FAUST discovered cell populations significantly associated with clinical outcome that are missed by other methods. Notably, manual gating did not identify statistically significant T cell correlates of outcome in the MCC anti-PD-1 T cell dataset. We note that the significant FAUST phenotypes involve combinations of PD-1, HLA-DR, and CD28 that were not targeted by the manual gating. When we applied FlowSOM and Phenograph to the MCC anti-PD-1 T cell dataset, we did not observe statistically significant correlates under a variety of analysis settings (Section 2.6). Subrahmanyam et al. also reported [33] that CITRUS did not produce a specific predictive model among subjects that went on to receive anti-PD-1 therapy. However, we were able to validate that the effector memory phenotypes discovered by FAUST in the MCC anti-PD-1 T cell dataset were significantly increased in responding subjects in the CyTOF dataset of Subrahmanyam et al. [33] using FAUST’s targeted top-down testing approach (Section 2.2). In part, our validation followed from the specificity of our hypotheses: we only tested 3 phenotypes of interest in the CyTOF dataset, while the authors reported examining 210 cell population fractions using FlowJo in the original analysis ([33]). Similar to the manual MCC analysis, none of the 210 cell populations manually examined in the original CyTOF analysis involved combinations of PD-1, HLA-DR, and CD28 (see Supplementary Figures S2, S4 in [33]). We believe that this is an important scientific feature of our method: when FAUST is used to perform unbiased discovery on datasets (such as in the MCC study), cell sub-populations that are found significant in subsequent modeling constitute testable hypotheses about specific cellular phenotypes (Section 2.1). Furthermore, when independent datasets are publicly available (such as the CyTOF dataset of Subrahmanyam et al. [33]), we demonstrated that FAUST itself can be used to test and validate those hypotheses (Section 2.2).

The cell sub-populations discovered by FAUST are consistent with their immunological context and recent literature. The PD-1 dim CD28+ HLA-DR+ CD8 T cell sub-population identified in the MCC anti-PD-1 trial may represent virus specific T cells as evidenced by their correlation with T cell clonality measurements from the tumor biopsy (Figure 3D). This further accords with literature that highlights the role of CD28 in anti-PD-1 immunotherapy, which reports CD28 signaling disrupted by PD-1 impairs T cell function [25]. It has also been reported that, following PD-1 blockade, CD28 is necessary for CD8 T cell proliferation [26]. The sub-populations are also consistent with reports that certain PD-1^int^ CD8^+^ T cells are responsible for viral control in mice [39] after PD-1 blockade [40]. Taken together with our findings, the PD-1 dim CD28+ HLA- DR+ CD8 T cell sub-population may have prognostic value in MCC subjects with virus-positive tumors, though we emphasize this hypothesis requires further validation in an independent cohort. Supporting this assertion is the surprisingly strong correlation between the CD8 T cell frequency and anti-PD-1 IHC measured from the tumor where the latter is a known prognostic marker. Although this evidence is tempered by small sample size, its strength warrants further investigation. The consistent detection of myeloid sub-populations with a CD14+CD16-HLA-DR+ phenotype across four different datasets from three independent trials spanning different cancer types and therapies strongly supports the assertion that the myeloid correlates represent real biological signal.

Our results demonstrate that FAUST can consistently detect immunologically-plausible candidate biomarkers from measurements made in blood using a simple, well-understood assay. They also demonstrate that FAUST may be of particular use in re-analyzing published datasets that have been distributed in public repositories (as well as giving one demonstration of the value of public repositories themselves) since we applied FAUST to high-dimensional public CyTOF data to validate the T cell correlates detected in the MCC study: without public repositories, we could not have conducted this validation analysis. Many large experimental cytometry datasets have already been published and FAUST could be used to systematically mine these datasets for standardized phenotypes. In turn, this could open the door to the re-analysis and meta-analysis of public datasets using approaches such as PFDA.

## 4 Methods

### 4.1 FAUST method: underlying statistical model

FAUST assumes the following criteria are met in a cytometry experiment consisting of *n* experimental units *E_i_*, 1 *≤ i ≤ n*.

#### Assumption 1.

Each sample in the cytometry experiment has been compensated (as needed) as well as pre-gated to remove debris and dead cells.

If pre-gating has not been performed by an investigator, computational methods [34, 41] can be used before applying FAUST to cytometry data in order to guarantee this assumption is met.

#### Assumption 2.

In each sample, measurements on the live cells are made using a common set of p transformed protein markers.

Let *n_i_* denote the number of events in the *i^th^* experimental unit. FAUST supposes each event *E_i_*,*_j_* in an experimental unit *E_i_*, of dimension *p* (the number of markers), arises as a sample from a finite mixture model

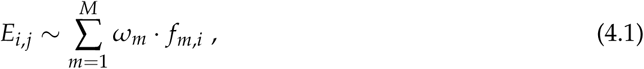

for 1 *≤ j ≤ ni*, with *M ∈* N, 0 *≤ ωm ≤* 1 and 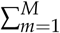 *ωm* = 1 for all 1 *≤ i ≤ n*. FAUST assumes the mixture components *f_m,i_* of an experimental unit in (4.1) are translated and scaled versions of a common collection of densities on the space of protein measurements: for all 1 *≤ i ≤ n* and 1 *≤ m ≤ M*, there exists *λ_m,i_* ∈ R*^p^*, *σ_m,i_* ∈ R such that

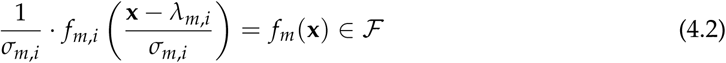

for each experimental unit *i*, with the common class *F* is defined as

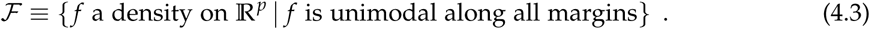

(4.2) expresses the fundamental modeling assumption: each mixture (4.1) that generates an experimental unit consists of a common set of densities (4.3), with unit-specific changes to location (the translations *λ_m,i_*) and scale (the scalar multiples *σ_m,i_*) of the component densities. These unit-specific modifications represent technical and biological effects. We emphasize that we only assume marginal unimodality for the *f* in (4.3), but make no assumptions about the joint-distribution of these densities. We also emphasize that while this model provides intuition for the subsequent FAUST algorithm, the algorithm itself does not fit the model (4.1) but rather assigns individual cells in an experimental unit to standardized modal regions of the joint-distribution using the strategy described in the following sections.

### 4.2 FAUST method: overview

FAUST is designed to perform independent approximate modal clustering of each mixture (4.1) in each experimental unit. Its approximation strategy is to use 1-dimensional densities to grow an exhaustive forest of gating strategies (section 4.3), from which it estimates a standardized set of annotation boundaries for all markers in a mixture, which exhibit 1-dimensional multimodality either marginally or across a large number of conditional 1-dimensional density estimates. Annotation boundaries are estimated (section 4.5) by taking a weighted average of marginal and conditional 1-dimensional antimodes for a marker that FAUST selects, using a score (section 4.4) that quantifies if the marker has persistent multimodality in the experimental unit. FAUST also uses the distribution of the depth score across units to select a subset of markers to use for cluster discovery and annotation (section 4.6).

FAUST defines a cluster as a subset of events in an experimental unit that fall inside either a conical or hyper-rectangular region bounded by the Cartesian product of the standardized set of annotation boundaries. FAUST discovers cluster phenotypes by growing a forest of partition trees for each experimental unit (trees are grown at random, following a strategy related to growing the annotation forest), and locating a sub-collection of homogeneous leaf nodes in the forest relative to the standardized phenotypic boundaries (section 4.8). FAUST collects a list of phenotypes discovered in each experimental unit and counts how often each phenotype appears across the set of lists. If a phenotype exceeds a user-specified filtering threshold, FAUST will annotate that cluster in each experimental unit relative to the standardized annotation boundaries. Intuitively, each annotation is a pointer to a modal region of each experimental unit’s mixture distribution. FAUST concludes by deriving a count matrix, with each row corresponding to a sample in the experiment, each column an annotated cluster, and each entry the cell count corresponding to the annotated cluster in the sample.

### 4.3 FAUST method: growing the annotation forest

For all markers in a sample, all cells for each marker are tested for unimodality using the dip test [42]. The hypothesis of unimodality is rejected for any marker that has dip test p-values below 0.25. All markers which are deemed multimodal according to this dip criterion are then used to start gating strategies. Gate locations for each strategy are determined using the taut string density estimator [43]. The location of each gate is the mid-point of any anti-modal component of the taut string. Since the taut string makes no assumptions about the number of modes present in a density, in principle this approach can lead to estimating an arbitrary number of gates in a given strategy. In practice, we only pursue strategies containing 4 or fewer gates under the assumption that marker expression of 5 or more expression categories does not reflect biological signal.

Once the initial set of gates are computed for a given marker, events are divided into sub-collections relative to the gates for that marker and the procedure recurses and repeats along each sub-collection. Algorithm1gives an overview of the procedure. A gating strategy terminates when it meets any of the following stopping conditions. First, once a strategy involves any three combinations of markers, it terminates. This is because the space of gating strategies grows factorially with the number of markers. Due to this growth rate, nodes in the forest are penalized factorially relative to their depth in the gating strategy when we subsequently compute the depth score. Second, if at any point in a strategy FAUST fails to reject the null hypothesis of unimodality for all tested markers, the strategy terminates regardless of depth. Finally, a gating strategy terminates along a branch if all nodes along the branch contain too few cells. The algorithm displayed here assumes event measurements are distinct in the cytometry dataset, and all nodes in the forest contain in excess of 500 events. For details of how FAUST breaks ties and deals with nodes containing between 25 and 500 events, we refer the reader to [44].

### 4.4 FAUST method: depth score computation

Suppose there are *p >* 1 active markers in a sample. To compute the depth score for any of the *p* markers, the annotation forest is first examined to determine the following quantities: *d*_1_, the number of times different markers are gated in the root population; *d*_2_, the number of times children of the root are gated; and *d*_3_ the number of times grandchildren of the root are gated. For *i ∈ {*1, 2, 3*}* define

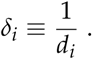

For 1 *≤ m ≤ p*, let

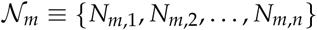

be the set of all *n* parent nodes in the annotation forest for which the null hypothesis of unimodality is rejected for marker *m*. For a parent node 1 *≤ j ≤ n*, let 1*_R_* denote the indicator function that is 1 when *N_m,j_* is the root population. Similarly, let 1*_C_* denote an indicator of a child of the root, and 1*_G_* a grandchild of the root. Define the scoring function

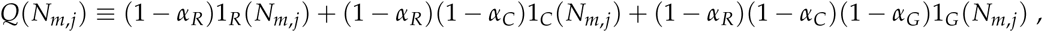

where, abusing notation, we let

*α_R_ ≡ α_R_* (*N_m_*,*_j_*) *≡* the dip test p-value in the root population of the gating strategy that led to *N_m_*,*_j_*.

#### Algorithm 1 Grow Annotation Forest

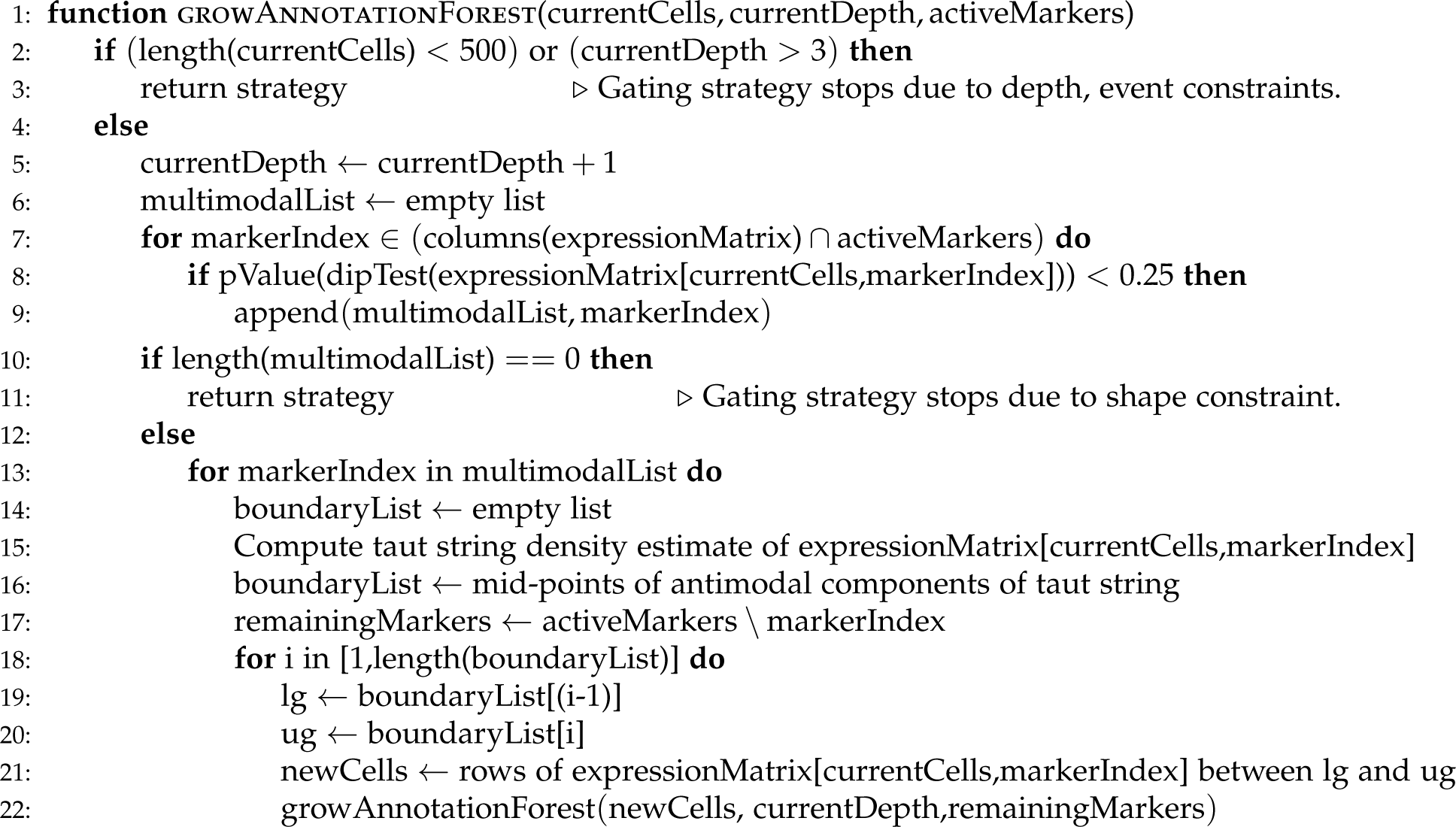

We allow *α_C_* and *α_G_* to be defined similarly. The function *Q* can be interpreted as a measure of the quality of the gating strategy that led to node *N_m,j_*. In the case of a grandchild node that had clear modal separation along all markers in the strategy, *Q*(*N_m,j_*) *≈* 1, while a grandchild node that had p-values of 0.25 at each ancestral node, *Q*(*N_m,j_*) *≈* 27/64 = 0.75^3^.

Let *𝒫_m_* be the population size for marker *m* in the root population. Next define

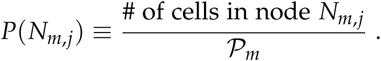

Finally, define

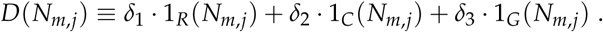

The depth score is defined to be the normalized sum

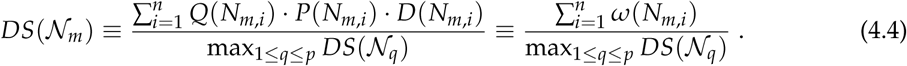

The depth score maps *𝒩_m_* into [0, 1], with at least one marker in a gated sample achieving the maximal score of 1. This is taken as a measure of separation quality: the best scoring marker according to the depth score is taken to be the best separated marker in that sample at the root population, and conditionally along all other gating strategies. Normalizing to the unit interval allows depth scores to be compared across experimental units for given markers. By using the factorial weights *δ_i_*, the depth score also explains why FAUST only explores gating strategies involving, at most, combinations of three markers in its scoring and marker selection phase. Adding more combinations of markers induces a factorial increase in computational cost. But any marker that enters a gating strategy at depth 4 (or beyond) will be dominated in depth score by those markers initially gated in the annotation forest at or near the root population. Consequently, after normalization in experiments with a large number of markers, such markers have depth score an *€* above zero, and are effectively never selected by FAUST for discovery and annotation. Hence the restriction to 3-marker gating strategies.

### 4.5 FAUST method: annotation boundary estimation

The depth score is also used to estimate annotation boundaries. Recalling FAUST only explores gating strategies with 4 or fewer annotation boundaries, FAUST partitions the set

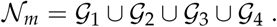

Define

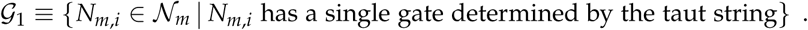

*𝒢*_2_, *𝒢*_3_, and *𝒢*_4_ are defined similarly. In other words each *𝒢_i_* is the subset of nodes in the annotation forest for marker *m* with *i* gates. Recalling (4.4) (which defines *ω*), we can partition the score sum

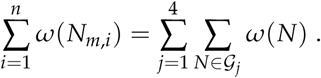

FAUST selects the number of annotation boundaries for the marker *m* by choosing the set *𝒢_j_* with the maximal sum ∑*_N∈Gj_ ω*(*N*). Letting *g*1 (*Nm*,*j*) denote the smallest gate location estimated by the taut string in node *N_m,j_* (which is the only gate location if FAUST selects *𝒢*_1_), FAUST estimates the phenotypic boundary locations for the marker by taking the weighted average

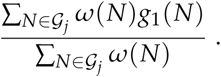

In the event FAUST selects *𝒢_j_*, *j >* 1 (i.e., multiple annotation boundaries), similar weighted averages are taken for *g*_2_ (*N_m,j_*), etc.

### 4.6 FAUST method: marker selection

Markers are selected by comparing the user-selected, empirical depth score quantile 4.9.6 across experimental units to a user-selected threshold value 4.9.7. All markers whose empirical quantile exceeds the threshold are used for discovery and annotation.

### 4.7 FAUST method: boundary standardization

FAUST standardizes the number of annotation boundaries for each marker by majority vote. The most frequently occurring number of annotation boundaries across experimental units is chosen as the *standard* number. This behavior can be modified via the preference list tuning parameter (see 4.9.8) in order to incorporate prior biological information into FAUST.

Next, for a given marker, FAUST selects the set of experimental units where the number of annotation boundaries for that marker matches the standard. Then, by rank, FAUST computes the median location of each phenotypic boundary across experimental units. We refer to these median boundary locations as the *standard boundaries*.

FAUST enforces standardization of annotation boundaries for non-conforming experimental units by imputation or deletion. Imputation in an experimental unit occurs when FAUST estimates fewer boundaries than the standard. In this case, each boundary in the non-conforming unit is matched to one of the standards by distance. Unmatched standards are used to impute the missing boundaries. Similar distance computations are done in the case of deletion, but FAUST deletes boundaries that are farthest from the standards. For both imputation and deletion, if multiple boundaries match the same standard, then the boundary minimizing the distance is kept, and the other boundaries are deleted. Should this result in standards that don’t map to any boundaries, then those unmatched standards are used to impute the missing boundaries.

If the user modifies the *imputation hierarchy* parameter (Methods 4.9), the previously described imputation process is modified in the following way. First, FAUST iterates over distinct values of the *imputation hierarchy* setting, and will attempt to impute missing boundaries for an experimental unit by only using data from experimental units with the same setting. Similarly, deletions occur only using data from experimental units with the same imputation hierarchy setting. Once this is complete, FAUST will then impute missing boundaries and delete excess boundaries using data across all experimental units.

### 4.8 FAUST method: phenotype discovery and cluster annotation

For each experimental unit, FAUST constructs a forest of partition trees (randomly sampled) and annotates selected leaves from this forest relative to the standardized annotation boundaries. Partition tree construction is similar to tree construction for the annotation forest (4.3), but they are not depth-constrained: a tree continues to grow following the previously described strategy until each leaf is unimodal according to the dip test [42] or contains fewer than 25 cells. Consequently, a single partition tree defines a clustering of an experimental unit. Clusterings from the forest of partition trees are combined into a single clustering in the following manner. To ensure cells are not assigned to multiple clusters, a subset of leaves of the partition forest are selected by scoring leaves according to shape criteria, and then selecting a subset of leaves across partition trees that share no cells to maximize their total shape score. The shape score is defined in terms of trimmed sample L moments [45, 46] as well dip test p values [42] of leaf margins. Only the selected leaves are given phenotypic annotations. Phenotypes are determined for a selected leaf by comparing the median of each margin to the standardized annotation boundary for the sample, and then assigning the leaf a label that locates the leaf relative to the sample boundaries. In the case of a single boundary for a phenotype, the location then maps to “-” or “+”; for two boundaries, “-”, “dim”, and “bright”. FAUST keeps a list of discovered phenotypes for each experimental unit, and concludes by returning exact counts of cells in each sample whose phenotypes exceed a user-specified occurrence frequency threshold. For more details of the scoring and selection procedure, we refer the reader to [44].

### 4.9 FAUST method: tuning parameters

We describe the key tuning parameters of FAUST.

#### 4.9.1 Experimental unit

This parameter is used to link individual experimental samples together into a single experimental unit. All samples with the same “experimental unit” value are concatenated prior to FAUST conducting discovery and annotation.

#### 4.9.2 Imputation hierarchy

This parameter is used to link experimental units together during the imputation process. All experimental units with the same “imputation hierarchy” value will define a block of units across which the boundary standardization process (4.7) first occurs. For example, suppose a study contains tissue samples and blood samples stained by the same marker panel. The imputation hierarchy can be used so that if a boundary is missing for a marker from a blood sample, FAUST will impute the missing boundary using boundaries from other blood samples if possible. Similarly, if a boundary is missing for a marker from a tissue sample, this parameter will have FAUST estimate the missing boundary location using other tissue samples alone if possible. If it is not possible to impute a missing boundary within a class (since the boundary is never estimated with the class), imputation then occurs across all experimental samples.

#### 4.9.3 Starting cell population

The name of the population in the manual gating strategy where FAUST conducts discovery and annotation.

#### 4.9.4 Active markers

A list of all markers in the experiment that can possibly be used for discovery and annotation in the starting cell population. FAUST will only compute the depth score for markers in this initial set.

#### 4.9.5 Marker boundary matrix

A 2 *× n* matrix of lower and upper protein expression bounds. By default, it is set for *−* inf and inf for all markers in a flow experiment. When the manual gating strategy does not remove all debris or doublets from the starting cell population, samples can appear to have clusters of events along at very low or very high expression values for some markers. By setting boundaries for those markers to exclude these doublet or debris clusters, FAUST treats all events below the lower and above the upper bounds as default low or high, respectively. These events are not dropped from the experiment. However, they are ignored when testing for multimodality and subsequent density estimation. In the case of mass cytometry experiments, the default lower boundary is set to 0 for all markers in an experiment in order to accommodate the zero-inflation common to mass cytometry data. The number of events in a marker that fall between the lower and upper marker boundaries in the starting cell population define the *effective sample size* for that marker.

#### 4.9.6 Depth-score selection quantile

The empirical quantile of a marker’s depth-score across all experimental units that is used to compare against a user-selected depth-score threshold. By default, this parameter is set to the median.

#### 4.9.7 Depth-score selection threshold

A value in [0, 1] used to select a subset of markers to be used in discovery and annotation based on their empirical depth score selection quantile. By default, this parameter is set to 0.01.

#### 4.9.8 Supervised Boundary Estimation List

Allows the user to modify FAUST’s default gate standardization methodology for each marker. This parameter is one way to incorporate prior (biological) knowledge in the FAUST procedure: if a marker is known to have a certain range of expression, such as low-dim-bright, this can be used to encourage or force FAUST to estimate the corresponding number of annotation boundaries from the data. Similarly, if FMO controls have been collected for a marker, this parameter can be used to set the phenotypic boundary according to the controls.

#### 4.9.9 Phenotype Occurrence Threshold

An integer value (set to 1 by default) used to include or exclude discovered phenotypes in the final count matrix returned by FAUST. If a phenotype appears at least Phenotype Occurrence Threshold times across experimental units, it is included in the final counts matrix. By default, all discovered phenotypes are included. Phenotypes exceeding the threshold are assumed to be biological signal while those that fall below it are assumed to be sample-or batch-specific effects. A consequence of this assumption is that all cells in a sample associated with any phenotype falling below the threshold are re-annotated with a common non-informative label indicating those phenotypes ought not be analyzed due to their rarity.

### 4.10 CITN-09 T cell Panel Analysis

The CITN-09 T cell staining panel is described in the supplementary information A.11.1. FAUST tuning parameter settings (above) for this dataset are described in supplementary section A.10.2. Between one and four samples were collected from 27 patients with stage IV and unresectable stage IIIB Merkel Cell Carcinoma and [22, 47] spanning the course of treatment. All 27 patients had samples collected at baseline (cycle C01, before initiation of anti-PD-1 therapy); 16 at cycle C02 (3 weeks post-treatment of the second cycle of therapy); 22 at cycle C05 (12 weeks post-treatment of the fifth cycle of therapy); and 13 at end of trial (EOT, patient specific). 18 of 27 subjects responded to therapy (CR/PR) for an observed response rate of 67%. Each sample was pre-gated to remove debris and identify live lymphocytes. Let *c_i,k_* denote the number of events in FAUST cluster *k* for sample *i*. Let *n_i_* denote the number of events in the *i^th^* subject’s baseline sample. Similar to [23], we assume *c_i,k_ ∼* Binomial(*n_i_*, *µ_i,k_*). Our model is

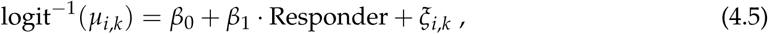

where Responder is an indicator variable equal to 1 when the subject exhibits complete or partial response to therapy, and 0 otherwise, and each 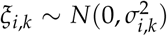 is a subject-level random effect. The R package **lme4** was used to fit all GLMMs [48].

### 4.11 CITN-09 Myeloid Panel

The CITN-09 Myeloid staining panel is described in supplementary information A.11.2. FAUST tuning parameter settings are described in supplementary information A.10.3. This dataset consisted of 69 samples stained to investigate myeloid cells. An initial screen comparing the ratio of the number of events in the singlet gate to the number of events in the root population led us to remove 3 samples from analysis due to low quality. We ran FAUST on the remaining 66 samples which consisted of 21 samples collected at cycle C01, before initiation of anti-PD-1 therapy; 16 at cycle C02; 18 at cycle C05; 1 at C08; and 10 at EOT. Of the 21 baseline samples, 1 was coded as inevaluable “NE”. This sample was removed from downstream statistical analysis. 13 of the 20 subjects with baseline samples available responded to therapy (PR/CR), for an observed response rate of 65%. Discovery and annotation was run at the individual sample level using cells in the “45+” node of the manual gating strategy. FAUST selected 11 markers: CD33, CD16, CD15, CD14, CD3, HLA-DR, CD20, CD19, CD11B, CD56, CD11C. FAUST annotated 147 cell sub-populations in terms of these markers, labeling 92.9% of the cells in the median sample. The statistical model used here is identical to (4.5), with counts are now derived from the 15 baseline samples.

### 4.12 CITN-07 Phenotyping Panel Analysis

We ran FAUST on this dataset comprising of a total of 358 longitudinal samples from 35 subjects in two cohorts (Cohort 1: with FLT-3 pre-treatment and Cohort 2: without pre-treatment), with between 4 and 12 samples per subject over four cycles of therapy and at end of trial. Subjects were given FLT-3 ligand seven days prior to the start of the first two of four treatment cycles. FLT-3 ligand was given to promote the expansion of myeloid and dendritic cell compartments in order to investigate whether expansion improved response to therapy. FAUST was configured to perform cell population discovery and annotation per sample in order to account for biological and technical heterogeneity. Debris, dead cells and non-lymphocytes were excluded by pre-gating. The CITN-07 Phenotyping staining panel is described in supplementary information A.11.3. FAUST tuning parameter settings are described in supplementary information A.10.1. FAUST discovered 128 cell populations. We tested each discovered cell population at the cohort-specific baseline (32 samples) for association with recurrence of disease (14 subjects had disease recur, 18 did not have disease recur). We analyzed the baseline counts using a model similar to (4.5). Here, the model was adjusted for subject-to-subject variability using a random effect, while cohort status, recurrence, and NYESO-1 staining of the tumor by immunohistochemistry (measured as positive, negative, or undetermined) were modeled as population effects.

### 4.13 Krieg et al. CyTOF Analysis

The markers used for the Krieg et al. [9] CyTOF panel are described in supplementary information A.11.4. FAUST tuning parameter settings are described in supplementary information A.10.4. We used FAUST to discover and annotate cell populations in the mass cytometry datasets stained to investigate myeloid cells. Following [9], we removed samples with fewer than 50 cells from our analysis, leaving 19 samples (from 19 subjects) at baseline for downstream statistical analysis. 10 of the 19 samples at baseline were from subjects that went on to exhibit response to therapy. To account for batch effects and small sample sizes, all samples within a batch were concatenated and processed by FAUST. FAUST selected 11 markers for discovery and annotation: CD16, CD11b, CD11c, CD14, ICAM-1, CD33, CD62L, CD7, PD-L1, CD56, and HLA-DR. FAUST annotated 40 cell sub-populations in terms of these markers, labeling 61.1% of cells in the median sample.

Our baseline model was similar to (4.5), but was modified by

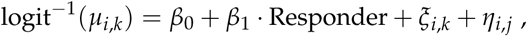

where *j ∈ {*1, 2*}*, and 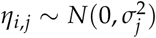 is a random effect included to model the batch effects.

### 4.14 Krieg et al. FACS Analysis

The Krieg et al. [9] FACS staining panel is described in supplementary information A.11.5. FAUST tuning parameter settings are described in supplementary information A.10.5. We used FAUST to process 31 baseline flow cytometry samples from responders and non-responders to therapy (16 responders, 15 non-responders). FAUST was run at the individual sample level on live cells from the manual gating strategy used by [9]. QC and review of the manual gating strategy let us to make manual adjustments to the “Lymphocytes” gate of 7 samples in this dataset. An example of this gate adjustment is shown in the supplementary information (S3) FAUST selected 9 markers for discovery and annotation: CD3, CD4, HLA-DR, CD14, CD19, CD11b, CD16, CD56, and CD45RO. FAUST annotated 48 cell sub-populations in terms of these markers, labeling 97.1% of cells in the median sample. The statistical model used here is identical to (4.5), with *c_i,k_* now denoting the 40 clusters in the FACS data, and *n_i_* refers to the baseline FACS sample counts.

### 4.15 Subrahmanyam et al. CyTOF Analysis

The markers used from the Subrahmanyam et al. [33] CyTOF panel is described in supplementary information A.11.6. FAUST tuning parameter settings are described in supplementary information A.10.6. We used openCyto [34] to reproduce the manual gating strategy reported in [33] to identify live intact singlets in each analyzed sample. We then used FAUST to process 64 pre-treatment, unstimulated CyTOF samples from responders and non-responders to Ipilimumab (anti-CTLA-4, 10 responders, 14 non-responders) and Pembrolizumab (anti-PD-1, 21 responders, 19 non-responders). FAUST was run through the boundary standardization phase at the individual sample level on live intact singlets (as identified by openCyto). FAUST was used to determined standardized annotation boundaries for 10 markers: CD4, CD3, CD8, CD45RA, HLA-DR, CD28, PD-1, CD25, CD127, CCR7. Once these boundaries were computed, cells with CD4+ or CD8+ phenotypes corresponding to significant phenotypes discovered in the MCC anti-PD-1 trial (Section 2.1) were targeted in each sample using the standardized boundaries. This produced counts for two cell populations. Counts from these two cell populations were taken from the 40 samples from subjects that went on to receive anti-PD-1 therapy, and tested for association with response to therapy. The model used here is identical to (4.5), *mutatis mutandis*.

### 4.16 PFDA multivariate model

We describe the multivariate PFDA model for the CD14+ CD16- HLA-DR+/bright cells; the T cell model is the same, modified only by the inclusion criteria. All FAUST phenotypes annotated as CD14+ CD16- HLA-DR+/bright CD3- CD56- CD19- and included in the univariate analysis were selected in CITN-07 (the FLT3-ligand + therapeuitc vx trial), CITN-09 (the MCC anti-PD-1 trial), and the Krieg et al. Melanoma anti-PD-1 trial FACS dataset [9]. Let *k^∗^* denote the number of FAUST phenotypes within a given study. Let *n* denote the number of subjects at baseline, and *N* = *n · k^∗^*. For 1 *≤ i ≤ N*, 1 *≤ j ≤ k^∗^* our statistical model is

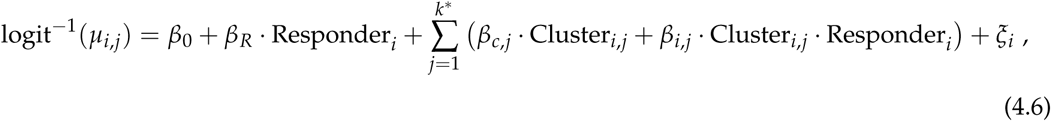

where Cluster*_i,j_* is an indicator variable that is 1 when observation *i* is from cluster *j* and 0 otherwise, Responder*_i_* is an indicator variable when observation *i* is taken from a responding subject, and 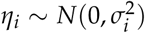 is an observation-level random effect. After estimating model coefficients *β_i,j_* in (4.6), we test for differential abundance by testing t for positivity of linear combination of the coefficients:

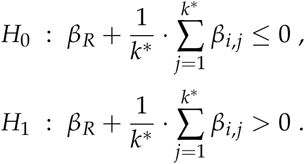

### 4.17 Code availability

FAUST is available as an R package athttps://github.com/RGLab/FAUST.

### 4.18 Author contributions

E.G., G.F., R.G. designed the FAUST method as well as statistical methods for analyzing FAUST cell populations. E.G., G.F. implemented FAUST and conducted data analyses using FAUST. E.G., G.F., R.G. contributed to design of data analysis plans, data analysis interpretation, and wrote the manuscript. L.A.D., C.D.C., C.M., N.R., J.M.T., P.T.N., M.A.C., S.P.F. contributed to the design of CITN-09 data analysis plans and data analysis interpretation. L.A.D., N.B., N.R., M.A.C., S.P.F. contributed to to the design of CITN-07 data analysis plans and data analysis interpretation. All authors discussed results and commented on the manuscript. All authors approve of this manuscript.

## Acknowledgments

The authors gratefully acknowledge the clinical trials patients and their families. The authors thank Dr. Suzanne Topalian for helpful discussions and critical review of the manuscript. This work was supported by [1P01CA22551701] to P.T.N., R.G., and E.G.; [UM1CA15496708] to M.A.C.; [R01GM118417] to G.F.; [K24-CA139052] to C.C., P.T.N.; [1U01CA154967] to M.A.C., S.P.F. from the National Cancer Institute; [P30-CA015704] to S.P.F., M.A.C., and P.T.N. from the NIH/NCI Cancer Center Support Grant in Seattle.

## Notes

https://github.com/RGLab/FAUST

## References

[1] Yvan Saeys, Sofie Van Gassen, and Bart N Lambrecht. Computational flow cytometry: helping to make sense of high-dimensional immunology data. Nature Reviews Immunology, 16(7):449, 2016.

[2] Gérald Grégori, Valery Patsekin, Bartek Rajwa, James Jones, Kathy Ragheb, Cheryl Holdman, and J Paul Robinson. Hyperspectral cytometry at the single-cell level using a 32-channel photodetector. Cytometry Part A, 81(1):35–44, 2012.

[3] Greg Finak, Marc Langweiler, Maria Jaimes, Mehrnoush Malek, Jafar Taghiyar, Yael Korin, Khadir Raddassi, Lesley Devine, Gerlinde Obermoser, Marcin L Pekalski, Nikolas Pontikos, Alain Diaz, Susanne Heck, Federica Villanova, Nadia Terrazzini, Florian Kern, Yu Qian, Rick Stanton, Kui Wang, Aaron Brandes, John Ramey, Nima Aghaeepour, Tim Mosmann, Richard H Scheuermann, Elaine Reed, Karolina Palucka, Virginia Pascual, Bonnie B Blomberg, Frank Nestle, Robert B Nussenblatt, Ryan Remy Brinkman, Raphael Gottardo, Holden Maecker, and J Philip McCoy. Standardizing flow cytometry immunophenotyping analysis from the human ImmunoPhenotyping consortium. Sci. Rep., 6:20686, February 2016.

[4] Nima Aghaeepour, Greg Finak, The FlowCAP Consortium, The DREAM Consortium, Holger Hoos, Tim R Mosmann, Ryan Brinkman, Raphael Gottardo, and Richard H Scheuermann. Critical assessment of automated flow cytometry data analysis techniques. Nature methods, 10(3):228, 2013.

[5] Lukas M Weber and Mark D Robinson. Comparison of clustering methods for high-dimensional single-cell flow and mass cytometry data. Cytometry Part A, 89(12):1084–1096, 2016.

[6] Sofie Van Gassen, Britt Callebaut, Mary J Van Helden, Bart N Lambrecht, Piet Demeester, Tom Dhaene, and Yvan Saeys. Flowsom: Using self-organizing maps for visualization and interpretation of cytometry data. Cytometry Part A, 87(7):636–645, 2015.

[7] Eirini Arvaniti and Manfred Claassen. Sensitive detection of rare disease-associated cell subsets via representation learning. Nature communications, 8:14825, 2017.

[8] Robert V Bruggner, Bernd Bodenmiller, David L Dill, Robert J Tibshirani, and Garry P Nolan. Automated identification of stratifying signatures in cellular subpopulations. Proceedings of the National Academy of Sciences, 111(26):E2770–E2777, 2014.

[9] Carsten Krieg, Malgorzata Nowicka, Silvia Guglietta, Sabrina Schindler, Felix J Hartmann, Lukas M Weber, Reinhard Dummer, Mark D Robinson, Mitchell P Levesque, and Burkhard Becher. High-dimensional single-cell analysis predicts response to anti-pd-1 immunotherapy. Nature medicine, 24(2):144, 2018.

[10] Joseph A. Fraietta, Simon F. Lacey, Elena J. Orlando, Iulian Pruteanu-Malinici, Mercy Gohil, Stefan Lundh, Alina C. Boesteanu, Yan Wang, Roddy S. OâĂ Ź Connor, Wei-Ting Hwang, Edward Pequignot, David E. Ambrose, Changfeng Zhang, Nicholas Wilcox, Felipe Bedoya, Corin Dorfmeier, Fang Chen, Lifeng Tian, Harit Parakandi, Minnal Gupta, Regina M. Young, F. Brad Johnson, Irina Kulikovskaya, Li Liu, Jun Xu, Sadik H. Kassim, Megan M. Davis, Bruce L. Levine, Noelle V. Frey, Donald L. Siegel, Alexander C. Huang, E. John Wherry, Hans Bitter, Jennifer L. Brogdon, David L. Porter, Carl H. June, and J. Joseph Melenhorst. Determinants of response and resistance to cd19 chimeric antigen receptor (car) t cell therapy of chronic lymphocytic leukemia. Nature medicine, 24(5):563, 2018.

[11] Nima Aghaeepour, Radina Nikolic, Holger H Hoos, and Ryan R Brinkman. Rapid cell population identification in flow cytometry data. Cytometry Part A, 79(1):6–13, 2011.

[12] Markus Lux, Ryan Remy Brinkman, Cedric Chauve, Adam Laing, Anna Lorenc, Lucie Abeler-Dörner, Barbara Hammer, and Jonathan Wren. flowlearn: Fast and precise identification and quality checking of cell populations in flow cytometry. Bioinformatics, 1:9, 2018.

[13] Yvan Saeys, Sofie Van Gassen, and Bart Lambrecht. Response to Orlova et al.âĂ IJscience not art: statistically sound methods for identifying subsets in multi-dimensional flow and mass cytometry data sets”. Nature Reviews Immunology, 18(1):78, 2018.

[14] Guenther Walther, Noah Zimmerman, Wayne Moore, David Parks, Stephen Meehan, Ilana Belitskaya, Jinhui Pan, and Leonore Herzenberg. Automatic clustering of flow cytometry data with density-based merging. Advances in bioinformatics, 2009, 2009.

[15] Kenneth Lo, Ryan Remy Brinkman, and Raphael Gottardo. Automated gating of flow cytometry data via robust model-based clustering. Cytometry Part A: the journal of the International Society for Analytical Cytology, 73(4):321–332, 2008.

[16] Daniel Commenges, Chariff Alkhassim, Raphael Gottardo, Boris Hejblum, and Rodolphe Thiebaut. cytometree: A binary tree algorithm for automatic gating in cytometry analysis. bioRxiv, page 335554, 2018.

[17] Darya Y Orlova, Noah Zimmerman, Stephen Meehan, Connor Meehan, Jeffrey Waters, Eliver EB Ghosn, Alexander Filatenkov, Gleb A Kolyagin, Yael Gernez, Shanel Tsuda, Wayne Moore, Richard B. Moss, Leonore A. Herzenberg, and Guenther Walther. Earth moverâĂ Ź s distance (emd): a true metric for comparing biomarker expression levels in cell populations. PloS one, 11(3):e0151859, 2016.

[18] Darya Y Orlova, Stephen Meehan, David Parks, Wayne A Moore, Connor Meehan, Qian Zhao, Eliver EB Ghosn, Leonore A Herzenberg, and Guenther Walther. Qfmatch: multidimensional flow and mass cytometry samples alignment. Scientific reports, 8(1):3291, 2018.

[19] Lukas M. Weber, Malgorzata Nowicka, Charlotte Soneson, and Mark D Robinson. diffcyt: Differential discovery in high-dimensional cytometry via high-resolution clustering. Nature Communications Biology, 2019.

[20] Zicheng Hu, Chethan Jujjavarapu, Jacob J Hughey, Sandra Andorf, Hao-Chih Lee, Pier Federico Gherardini, Matthew H Spitzer, Cristel G Thomas, John Campbell, Patrick Dunn, Jeff Wiser, Brian A. Kidd, Joel T. Dudley, Garry P. Nolan, Sanchita Bhattacharya, and Atul J. Butte. Metacyto: A tool for automated meta-analysis of mass and flow cytometry data. Cell Reports, 24(5):1377–1388, 2018.

[21] Di Wu and Gordon K Smyth. Camera: a competitive gene set test accounting for inter-gene correlation. Nucleic Acids Res., 40(17):e133, September 2012.

[22] Paul Nghiem, Shailender Bhatia, Evan J. Lipson, William H. Sharfman, Ragini R. Kudchadkar, Andrew S. Brohl, Phillip A. Friedlander, Adil Daud, Harriet M. Kluger, Sunil A. Reddy, Brian C. Boulmay, Adam I. Riker, Melissa A. Burgess, Brent A. Hanks, Thomas Olencki, Kim Margolin, Lisa M. Lundgren, Abha Soni, Nirasha Ramchurren, Candice Church, Song Y. Park, Michi M. Shinohara, Bob Salim, Janis M. Taube, Steven R. Bird, Nageatte Ibrahim, Steven P. Fling, Blanca Homet Moreno, Elad Sharon, Martin A. Cheever, and Suzanne L. Topalian. Durable tumor regression and overall survival in patients with advanced merkel cell carcinoma receiving pembrolizumab as first-line therapy. Journal of Clinical Oncology, 37(9):693–702, 2019. PMID: 30726175.

[23] Malgorzata Nowicka, Carsten Krieg, Lukas M Weber, Felix J Hartmann, Silvia Guglietta, Burkhard Becher, Mitchell P Levesque, and Mark D Robinson. Cytof workflow: differential discovery in high-throughput high-dimensional cytometry datasets. F1000Research, 6, 2017.

[24] E.A. Eisenhauer, P. Therasse, J. Bogaerts, L.H. Schwartz, D. Sargent, R. Ford, J. Dancey, S. Arbuck, S. Gwyther, M. Mooney, L. Rubinstein, L. Shankar, L. Dodd, R. Kaplan, D. Lacombe, and J. Verweij. New response evaluation criteria in solid tumours: Revised recist guideline (version 1.1). European Journal of Cancer, 45(2):228 – 247, 2009. Response assessment in solid tumours (RECIST): Version 1.1 and supporting papers.

[25] Enfu Hui, Jeanne Cheung, Jing Zhu, Xiaolei Su, Marcus J. Taylor, Heidi A. Wallweber, Dibyendu K. Sasmal, Jun Huang, Jeong M. Kim, Ira Mellman, and Ronald D. Vale. T cell costimulatory receptor cd28 is a primary target for pd-1–mediated inhibition. Science, 355(6332):1428–1433, 2017.

[26] Alice O. Kamphorst, Andreas Wieland, Tahseen Nasti, Shu Yang, Ruan Zhang, Daniel L. Barber, Bogumila T. Konieczny, Candace Z. Daugherty, Lydia Koenig, Ke Yu, Gabriel L. Sica, Arlene H. Sharpe, Gordon J. Freeman, Bruce R. Blazar, Laurence A. Turka, Taofeek K. Owonikoko, Rathi N. Pillai, Suresh S. Ramalingam, Koichi Araki, and Rafi Ahmed. Rescue of exhausted cd8 t cells by pd-1–targeted therapies is cd28-dependent. Science, 355(6332):1423– 1427, 2017.

[27] L. McInnes, J. Healy, and J. Melville. UMAP: Uniform Manifold Approximation and Projection for Dimension Reduction. ArXiv e-prints, February 2018.

[28] Etienne Becht, Leland McInnes, John Healy, Charles-Antoine Dutertre, Immanuel WH Kwok, Lai Guan Ng, Florent Ginhoux, and Evan W Newell. Dimensionality reduction for visualizing single-cell data using umap. Nature biotechnology, 37(1):38, 2019.

[29] Alan L Landay, Carl E Mackewicz, and Jay A Levy. An activated cd8+ t cell phenotype correlates with anti-hiv activity and asymptomatic clinical status. Clinical immunology and immunopathology, 69(1):106–116, 1993.

[30] Nicolas A. Giraldo, Peter Nguyen, Elizabeth L. Engle, Genevieve J. Kaunitz, Tricia R. Cottrell, Sneha Berry, Benjamin Green, Abha Soni, Jonathan D. Cuda, Julie E. Stein, Joel C. Sunshine, Farah Succaria, Haiying Xu, Aleksandra Ogurtsova, Ludmila Danilova, Candice D. Church, Natalie J. Miller, Steve Fling, Lisa Lundgren, Nirasha Ramchurren, Jennifer H. Yearley, Evan J. Lipson, Mac Cheever, Robert A. Anders, Paul T. Nghiem, Suzanne L. Topalian, and Janis M. Taube. Multidimensional, quantitative assessment of pd-1/pd-l1 expression in patients with merkel cell carcinoma and association with response to pembrolizumab. Journal for immunotherapy of cancer, 6(1):99, 2018.

[31] Natalie J. Miller, Candice D. Church, Steven P. Fling, Rima Kulikauskas, Nirasha Ramchurren, Michi M. Shinohara, Harriet M. Kluger, Shailender Bhatia, Lisa Lundgren, Martin A. Cheever, Suzanne L. Topalian, and Paul Nghiem. Merkel cell polyomavirus-specific immune responses in patients with merkel cell carcinoma receiving anti-pd-1 therapy. Journal for immunotherapy of cancer, 6(1):131, 2018.

[32] Josef Spidlen, Karin Breuer, Chad Rosenberg, Nikesh Kotecha, and Ryan R Brinkman. Flowrepository: A resource of annotated flow cytometry datasets associated with peer-reviewed publications. Cytometry Part A, 81(9):727–731, 2012.

[33] Priyanka B Subrahmanyam, Zhiwan Dong, Daniel Gusenleitner, Anita Giobbie-Hurder, Mariano Severgnini, Jun Zhou, Michael Manos, Lauren M Eastman, Holden T Maecker, and F Stephen Hodi. Distinct predictive biomarker candidates for response to anti-ctla-4 and anti-pd-1 immunotherapy in melanoma patients. Journal for immunotherapy of cancer, 6(1):18, 2018.

[34] Greg Finak, Jacob Frelinger, Wenxin Jiang, Evan W Newell, John Ramey, Mark M Davis, Spyros A Kalams, Stephen C De Rosa, and Raphael Gottardo. Opencyto: an open source infrastructure for scalable, robust, reproducible, and automated, end-to-end flow cytometry data analysis. PLoS computational biology, 10(8):e1003806, 2014.

[35] Rossella Melchiotti, Filipe Gracio, Shahram Kordasti, Alan K Todd, and Emanuele de Rinaldis. Cluster stability in the analysis of mass cytometry data. Cytometry A, 91(1):73–84, January 2017.

[36] L Fong, Y Hou, A Rivas, C Benike, A Yuen, G A Fisher, M M Davis, and E G Engleman. Altered peptide ligand vaccination with flt3 ligand expanded dendritic cells for tumor immunotherapy. Proc. Natl. Acad. Sci. U. S. A., 98(15):8809–8814, July 2001.

[37] Nina Bhardwaj, Anna C. Pavlick, Marc S. Ernstoff, Brent Allen Hanks, Mark R. Albertini, Jason John Luke, Michael Jay Yellin, Tibor Keler, Thomas A. Davis, Andrea Crocker, Laura Vitale, Chihiro Morishima, Philip Adam Friedlander, Martin A. Cheever, and Steven Fling. A phase ii randomized study of cdx-1401, a dendritic cell targeting ny-eso-1 vaccine, in patients with malignant melanoma pre-treated with recombinant cdx-301, a recombinant human flt3 ligand. Journal of Clinical Oncology, 34(15):9589, 2016.

[38] Jacob H. Levine, Erin F. Simonds, Sean C. Bendall, Kara L. Davis, El ad D. Amir, Michelle D. Tadmor, Oren Litvin, Harris G. Fienberg, Astraea Jager, Eli R. Zunder, Rachel Finck, Amanda L. Gedman, Ina Radtke, James R. Downing, Dana PeâĂ Ź er, and Garry P. Nolan. Data-driven phenotypic dissection of aml reveals progenitor-like cells that correlate with prognosis. Cell, 162(1):184 – 197, 2015.

[39] Ran He, Shiyue Hou, Cheng Liu, Anli Zhang, Qiang Bai, Miao Han, Yu Yang, Gang Wei, Ting Shen, Xinxin Yang, Lifan Xu, Xiangyu Chen, Yaxing Hao, Pengcheng Wang, Chuhong Zhu, Juanjuan Ou, Houjie Liang, Ting Ni, Xiaoyan Zhang, Xinyuan Zhou, Kai Deng, Yaokai Chen, Yadong Luo, Jianqing Xu, Hai Qi, Yuzhang Wu, and Lilin Ye. Follicular cxcr5-expressing cd8+ t cells curtail chronic viral infection. Nature, 537(7620):412, 2016.

[40] Jolanda Brummelman, Emilia M.C. Mazza, Giorgia Alvisi, Federico S. Colombo, Andrea Grilli, Joanna Mikulak, Domenico Mavilio, Marco Alloisio, Francesco Ferrari, Egesta Lopci, Pierluigi Novellis, Giulia Veronesi, and Enrico Lugli. High-dimensional single cell analysis identifies stem-like cytotoxic cd8+ t cells infiltrating human tumors. Journal of Experimental Medicine, 215(10):2520–2535, 2018.

[41] Greg Finak and Mike Jiang. Flowworkspace: Infrastructure for representing and interacting with the gated cytometry. R package version, 3(3), 2011.

[42] John A Hartigan and PM Hartigan. The Dip Test of Unimodality. The Annals of Statistics, pages 70–84, 1985.

[43] P Laurie Davies and Arne Kovac. Densities, spectral densities and modality. The Annals of Statistics, 32(3):1093–1136, 2004.

[44] Evan Greene, Greg Finak, and Raphael Gottardo. Selective clustering annotated using modes of projections. arXiv preprint arXiv:1807.10328, 2018.

[45] J.R.M. Hosking. L-moments: Analysis and estimation of distributions using linear combinations of order statistics. Journal of the Royal Statistical Society: Series B (Methodological), 52(1):105–124, 1990.

[46] J.R.M. Hosking. Some theory and practical uses of trimmed l-moments. Journal of Statistical Planning and Inference, 137(9):3024–3039, 2007.

[47] Paul T. Nghiem, Shailender Bhatia, Evan J. Lipson, Ragini R. Kudchadkar, Natalie J. Miller, Lakshmanan Annamalai, Sneha Berry, Elliot K. Chartash, Adil Daud, Steven P. Fling, Philip A. Friedlander, Harriet M. Kluger, Holbrook E. Kohrt, Lisa Lundgren, Kim Margolin, Alan Mitchell, Thomas Olencki, Drew M. Pardoll, Sunil A. Reddy, Erica M. Shantha, William H. Sharfman, Elad Sharon, Lynn R. Shemanski, Michi M. Shinohara, Joel C. Sunshine, Janis M. Taube, John A. Thompson, Steven M. Townson, Jennifer H. Yearley, Suzanne L. Topalian, and Martin A. Cheever. Pd-1 blockade with pembrolizumab in advanced merkel-cell carcinoma. New England Journal of Medicine, 374(26):2542–2552, 2016. PMID: 27093365.

[48] Douglas Bates, Martin Mächler, Ben Bolker, and Steve Walker. Fitting linear mixed-effects models using lme4. arXiv preprint arXiv:1406.5823, 2014.

[49] Christian Hennig, Marina Meila, Fionn Murtagh, and Roberto Rocci. Handbook of cluster analysis. CRC Press, 2015.

[50] Yoav Benjamini and Yosef Hochberg. Controlling the false discovery rate: a practical and powerful approach to multiple testing. Journal of the royal statistical society. Series B (Methodological), pages 289–300, 1995.

