## supplemental_data for "New interpretable machine learning method for single-cell data reveals correlates of clinical response to cancer immunotherapy"

### A Supplementary materials

#### A.1 Summary of CITN clinical trials

| | CITN-07 | CITN-09 |
|---|---|---|
| ClinicalTrials.gov Identifier* | NCT02129075 | NCT02267603 |
| Study Type* | Interventional (Clinical Trial) | Interventional (Clinical Trial) |
| Intervention Model* | Parallel Assignment | Single Group Assignment |
| Masking* | None (Open Label) | None (Open Label) |
| Enrollment* | 100 participants (Estimated, recorded 6/11/2019) | 50 participants (Actual, recorded 6/11/2019) |
| Primary Purpose* | Treatment | Treatment |
| Actual Study Start Date* | April 9, 2014 | November 25, 2014 |
| Primary Completion Date* | February 2, 2020 (Estimated, recorded 6/11/2019) | February 6, 2018 (Actual, recorded 6/11/2019 |
| Total number of longitudinal samples analyzed by FAUST | 358 | 78 (T cell pane), 66 (Myeloid panel) |
| Total number of baseline samples (pre-treatment) | 32 | 27 (T cell panel), 20 (Myeloid panel) |
| Phase* | Phase 2 | Phase 2 |
| Intervention/Treatment* | Biological: DEC-205/NY-ESO-1 Fusion Protein CDX-1401 Other: Laboratory Biomarker Analysis Biological: Neoantigen-based Melanoma-Poly-ICLC Vaccine Other: Pharmacological Study Biological: Recombinant Flt3 Ligand | Biological: Pembrolizumab Other: Laboratory Biomarker Analysis |
| Condition/Disease* | Cutaneous Melanoma Mucosal Melanoma NY-ESO-1 Positive Tumor Cells Present Ocular Melanoma Stage IIB Cutaneous Melanoma AJCC v6 and v7 Stage IIC Cutaneous Melanoma AJCC v6 and v7 Stage III Cutaneous Melanoma AJCC v7 Stage IIIA Cutaneous Melanoma AJCC v7 Stage IIIB Cutaneous Melanoma AJCC v7 Stage IIIC Cutaneous Melanoma AJCC v7 Stage IV Cutaneous Melanoma AJCC v6 and v7 | Recurrent Merkel Cell Carcinoma Stage III Merkel Cell Carcinoma AJCC v7 Stage IIIA Merkel Cell Carcinoma AJCC v7 Stage IIIB Merkel Cell Carcinoma AJCC v7 Stage IV Merkel Cell Carcinoma AJCC v7 |

Table S1: Data listed in all rows with * taken from https://clinicaltrials.gov on June 11, 2019.

#### A.2 Baseline predictors MCC anti-PD-1 trial myeloid phenotyping panel

| Population | Effect Size | Lower 2.5% | Upper 97.5% | BonferroniP |
|---|---|---|---|---|
| CD33 Bright CD16- CD15- CD14- CD3- HLA-DR Bright CD20- CD19- CD11B+ CD56- CD11C+ | 1.923 | 0.902 | 2.952 | 0.017 |
| CD33 Bright CD16- CD15- CD14+ CD3- HLA-DR Bright CD20- CD19- CD11B+ CD56- CD11C- | 2.600 | 1.168 | 4.095 | 0.032 |
| CD33 Bright CD16- CD15- CD14+ CD3- HLA-DR Bright CD20- CD19- CD11B+ CD56- CD11C+ | 2.618 | 1.151 | 4.089 | 0.036 |
| CD33 Bright CD16- CD15+ CD14+ CD3- HLA-DR Bright CD20- CD19- CD11B+ CD56- CD11C+ | 2.944 | 1.474 | 4.557 | 0.009 |

Table S2: All statistically significant (bonferroni adjusted significance threshold of 5%) from the MCC anti-PD-1 trial.

#### A.3 Baseline predictors FLT3-L + therapeutic Vx trial.

The complete set of baseline predictors from the FLT3-L + therapeutic Vx trial are listed in Table S3. The top populations, by magnitude, were CD14+CD16- monocyte populations.

| Population | estimate | lower | upper | std.error | statistic | p.value | adjusted.p.value |
|---|---|---|---|---|---|---|---|
| CD8- CD3- HLA-DR Bright CD19- CD14+ CD11C+ CD4- CD123- CD16- CD56- | 2.42 | 1.12 | 3.72 | 0.66 | 3.65 | 1.29e-04 | 0.02 |
| CD8 Dim CD3- HLA-DR Bright CD19- CD14+ CD11C+ CD4- CD123- CD16- CD56- | 2.21 | 0.98 | 3.44 | 0.63 | 3.53 | 2.09e-04 | 0.03 |
| CD8 Dim CD3- HLA-DR Dim CD19- CD14- CD11C- CD4- CD123- CD16+ CD56+ | 1.90 | 0.92 | 2.87 | 0.50 | 3.81 | 6.84e-05 | 0.01 |
| CD8- CD3- HLA-DR Dim CD19- CD14- CD11C- CD4- CD123- CD16+ CD56+ | 1.60 | 0.70 | 2.49 | 0.46 | 3.49 | 2.40e-04 | 0.03 |
| CD8 Dim CD3- HLA-DR- CD19- CD14- CD11C- CD4- CD123- CD16+ CD56+ | 1.60 | 0.72 | 2.47 | 0.44 | 3.59 | 1.67e-04 | 0.02 |
| CD8 Bright CD3+ HLA-DR Bright CD19- CD14- CD11C- CD4- CD123- CD16- CD56- | 1.48 | 0.69 | 2.28 | 0.41 | 3.66 | 1.27e-04 | 0.02 |
| CD8- CD3- HLA-DR- CD19- CD14- CD11C- CD4- CD123- CD16+ CD56+ | 1.46 | 0.60 | 2.32 | 0.44 | 3.33 | 4.42e-04 | 0.06 |
| CD8 Dim CD3- HLA-DR Dim CD19- CD14- CD11C- CD4- CD123- CD16+ CD56- | 1.24 | 0.78 | 1.70 | 0.24 | 5.26 | 7.03e-08 | 0.00 |
| CD8- CD3- HLA-DR Bright CD19- CD14- CD11C- CD4- CD123- CD16- CD56- | 1.22 | 0.60 | 1.83 | 0.31 | 3.87 | 5.39e-05 | 0.01 |
| CD8- CD3- HLA-DR Dim CD19- CD14- CD11C- CD4- CD123- CD16+ CD56- | 1.12 | 0.70 | 1.54 | 0.21 | 5.25 | 7.63e-08 | 0.00 |
| CD8 Dim CD3- HLA-DR- CD19- CD14- CD11C- CD4- CD123- CD16+ CD56- | 0.98 | 0.59 | 1.36 | 0.19 | 5.01 | 2.70e-07 | 0.00 |
| CD8- CD3- HLA-DR- CD19- CD14- CD11C- CD4- CD123- CD16+ CD56- | 0.93 | 0.55 | 1.32 | 0.19 | 4.79 | 8.48e-07 | 0.00 |
| CD8 Dim CD3- HLA-DR Dim CD19- CD14- CD11C- CD4- CD123- CD16- CD56- | 0.82 | 0.33 | 1.31 | 0.25 | 3.30 | 4.81e-04 | 0.06 |
| CD8- CD3- HLA-DR Dim CD19- CD14- CD11C- CD4- CD123- CD16- CD56- | 0.77 | 0.34 | 1.19 | 0.22 | 3.54 | 2.02e-04 | 0.03 |

Table S3: All statistically significant (Bonferroni adjusted significance threshold of 5%) from the FLT3-L + therapeutic Vx trial.

#### A.4 The temporal abundance of of manually gated PD-1+ CD8 T cells in the in MCC anti-PD-1 trial

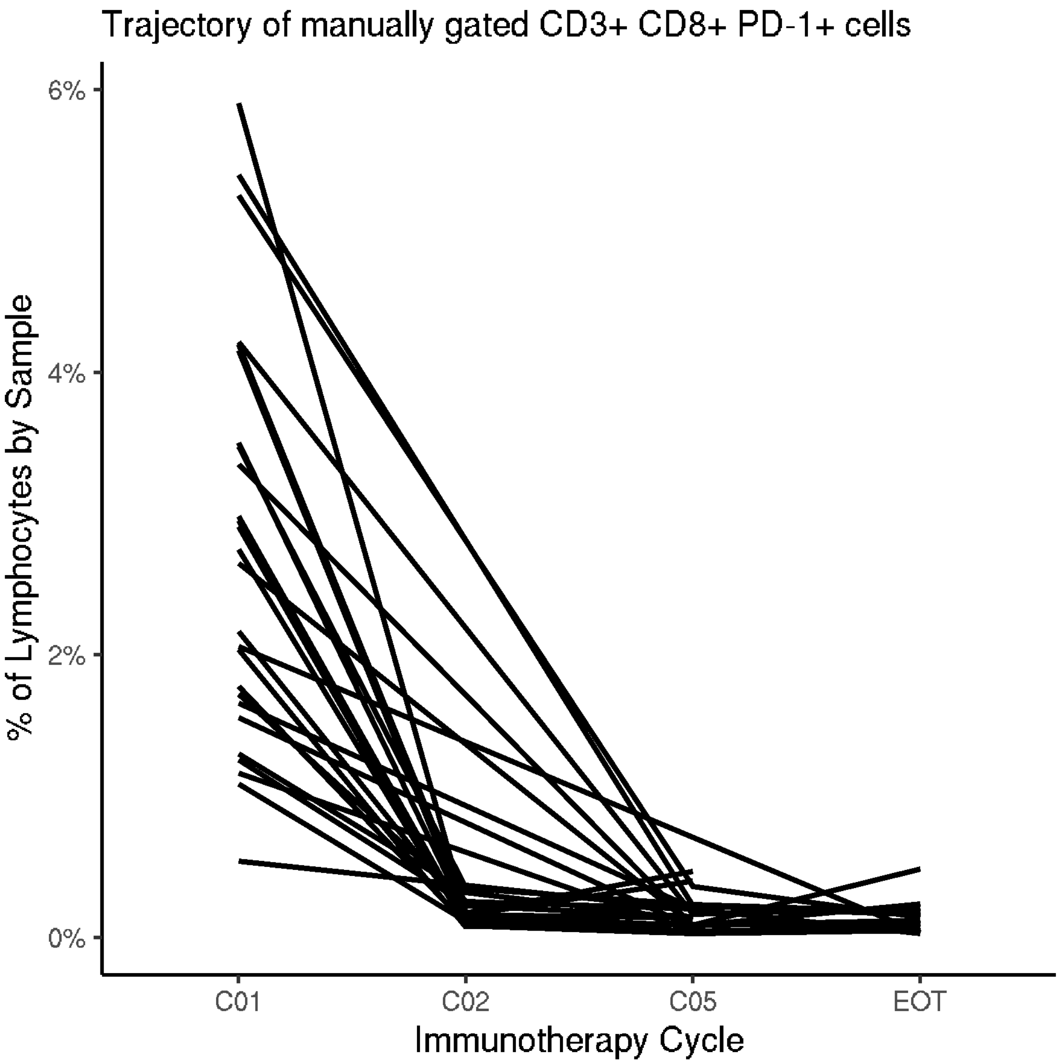


Figure S1: This figure displays the temporal abundance of manually gated CD3+ CD8+ PD-1+ cells across all collected time-points in the MCC anti-PD-1 trial discussed in section 2.1. Each line represents an individual subject.

#### A.5 Additional sub-populations associated with with clinical outcome at baseline in CITN-09

| Phenotype | fdr.adjusted.p.value |
|---|---|
| CD4- CD8+ CD3+ CD45RA- HLADR+ PD1 Dim CD28+ CD127- CD25- CCR7- | 0.010 |
| CD4 Bright CD8- CD3+ CD45RA- HLADR- PD1 Dim CD28+ CD127- CD25- CCR7- | 0.010 |
| CD4- CD8+ CD3+ CD45RA- HLADR+ PD1 Bright CD28+ CD127- CD25- CCR7- | 0.017 |
| CD4 Bright CD8- CD3+ CD45RA- HLADR+ PD1 Dim CD28+ CD127- CD25- CCR7- | 0.021 |
| CD4- CD8+ CD3+ CD45RA- HLADR- PD1 Dim CD28+ CD127- CD25- CCR7- | 0.088 |
| CD4- CD8+ CD3+ CD45RA+ HLADR- PD1 Dim CD28- CD127- CD25- CCR7- | 0.100 |
| CD4- CD8+ CD3+ CD45RA+ HLADR- PD1 Dim CD28- CD127- CD25+ CCR7- | 0.100 |
| CD4 Bright CD8- CD3+ CD45RA- HLADR- PD1 Dim CD28+ CD127- CD25- CCR7+ | 0.100 |
| CD4 Bright CD8- CD3+ CD45RA+ HLADR- PD1 Dim CD28- CD127- CD25- CCR7- | 0.100 |
| CD4 Bright CD8- CD3+ CD45RA+ HLADR- PD1 Dim CD28+ CD127+ CD25- CCR7- | 0.100 |
| CD4- CD8+ CD3+ CD45RA- HLADR- PD1 Bright CD28+ CD127- CD25- CCR7- | 0.103 |
| CD4 Dim CD8+ CD3+ CD45RA- HLADR- PD1 Dim CD28+ CD127+ CD25- CCR7- | 0.103 |
| CD4 Bright CD8- CD3+ CD45RA+ HLADR- PD1 Dim CD28+ CD127- CD25- CCR7+ | 0.113 |
| CD4 Bright CD8- CD3+ CD45RA- HLADR- PD1 Dim CD28+ CD127+ CD25- CCR7- | 0.139 |
| CD4 Bright CD8- CD3+ CD45RA- HLADR- PD1 Dim CD28+ CD127+ CD25- CCR7+ | 0.139 |
| CD4 Bright CD8- CD3+ CD45RA- HLADR- PD1 Dim CD28+ CD127- CD25+ CCR7- | 0.153 |
| CD4 Bright CD8- CD3+ CD45RA- HLADR- PD1 Bright CD28+ CD127- CD25- CCR7- | 0.153 |
| CD4 Bright CD8- CD3+ CD45RA+ HLADR- PD1- CD28+ CD127- CD25- CCR7+ | 0.153 |
| CD4- CD8+ CD3+ CD45RA- HLADR- PD1 Dim CD28- CD127- CD25- CCR7- | 0.157 |
| CD4- CD8+ CD3+ CD45RA- HLADR- PD1 Dim CD28+ CD127+ CD25- CCR7- | 0.157 |
| CD4- CD8+ CD3+ CD45RA+ HLADR- PD1 Dim CD28+ CD127+ CD25+ CCR7+ | 0.157 |
| CD4 Bright CD8- CD3+ CD45RA- HLADR- PD1- CD28+ CD127- CD25- CCR7+ | 0.171 |
| CD4 Bright CD8- CD3+ CD45RA+ HLADR- PD1- CD28+ CD127+ CD25- CCR7+ | 0.177 |
| CD4- CD8+ CD3+ CD45RA- HLADR- PD1- CD28+ CD127- CD25- CCR7- | 0.187 |
| CD4 Bright CD8- CD3+ CD45RA- HLADR- PD1- CD28+ CD127- CD25- CCR7- | 0.187 |
| CD4 Bright CD8- CD3+ CD45RA- HLADR- PD1 Dim CD28- CD127- CD25- CCR7- | 0.187 |
| CD4- CD8+ CD3+ CD45RA+ HLADR- PD1 Dim CD28+ CD127- CD25- CCR7- | 0.195 |

Table S4: FAUST Phenotypes associated with outcome in the CITN-09 T cell data at the FDR-adjusted 20% level.

#### A.6 Effect Sizes and Confidence Intervals in CITN-09 T cell panel

| Population | Effect Size (Log odds) | Lower 2.5% | Upper 97.5% |
|---|---|---|---|
| CD4 Bright CD8- CD3+ CD45RA- HLA-DR- PD-1 Dim CD28+ CD127- CD25- CCR7- | 2.030 | 1.003 | 3.126 |
| CD4- CD8+ CD3+ CD45RA- HLA-DR+ PD-1 Dim CD28+ CD127- CD25- CCR7- | 1.845 | 0.882 | 2.893 |
| CD4- CD8+ CD3+ CD45RA- HLA-DR+ PD-1 Bright CD28+ CD127- CD25- CCR7- | 1.871 | 0.814 | 2.981 |
| CD4 Bright CD8- CD3+ CD45RA- HLA-DR+ PD-1 Dim CD28+ CD127- CD25- CCR7- | 1.826 | 0.758 | 2.962 |

#### A.7 openCyto Gating Strategy replication

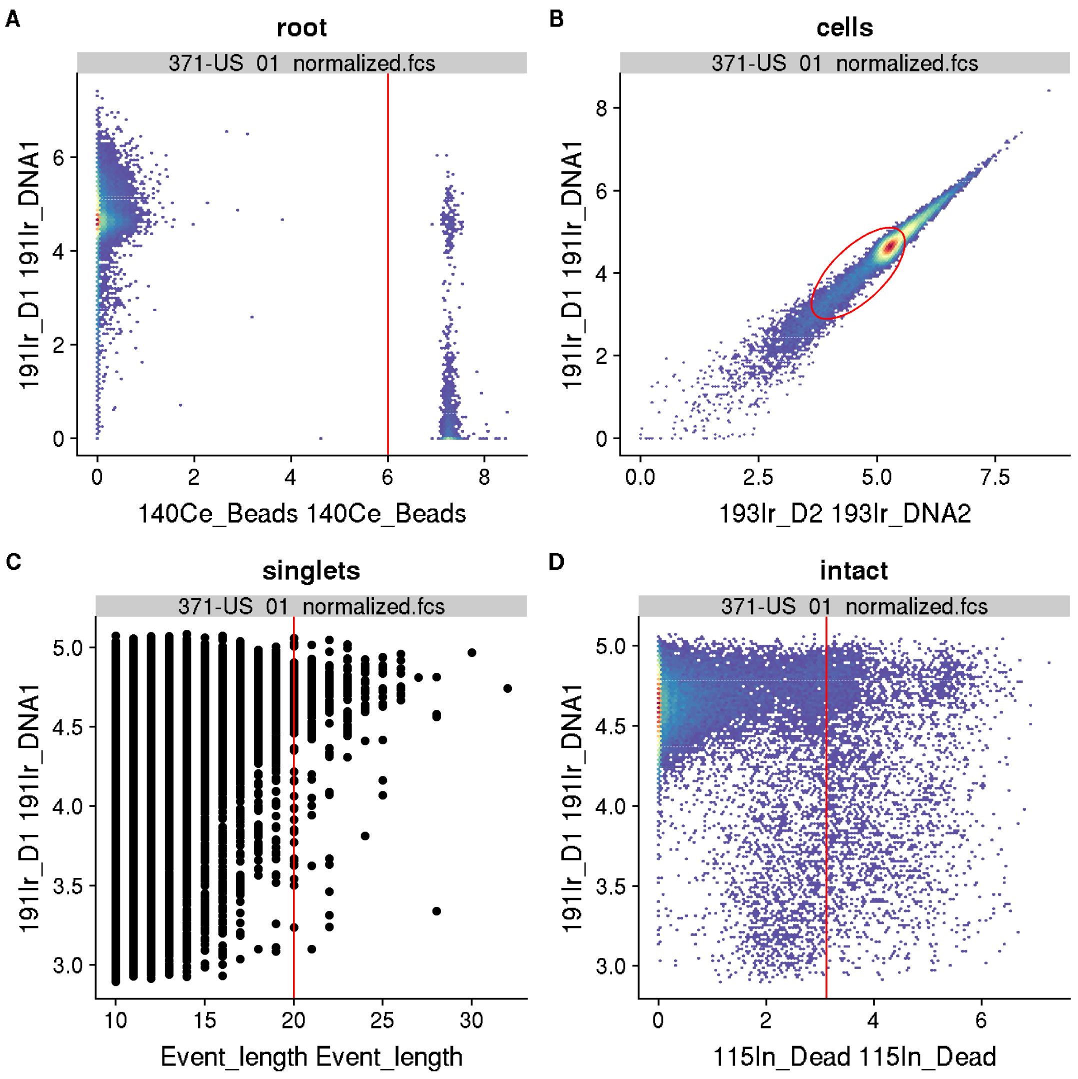


Figure S2: This supplementary figure shows an example of the openCyto [34] replication of the gating strategy described in Figure 1 of [33]. In the displayed sample, live intact singlets are identified.

#### A.8 Gating strategy modification examples

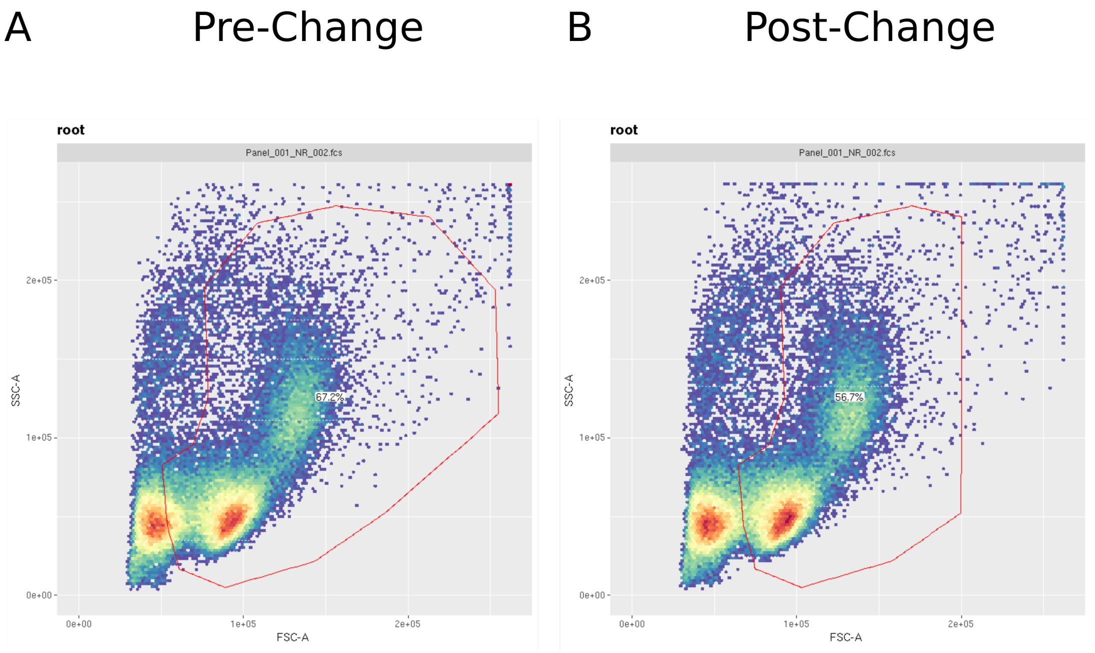


Figure S3: An example of modification to the manual gating strategy of the Krieg et al. FACS data. Panel A shows the initial manual gating strategy for the Lymphocytes of a sample. Panel B shows the same sample with the modified gate.

#### A.9 Alternative analysis of CITN-09 T cell panel

| Method | Transformation | Input Data | Num Clusters | Best Bonferroni | Second Bonferroni | Best FDR | Second FDR |
|---|---|---|---|---|---|---|---|
| FAUST | Asinh | All | 133 | 0.00 | 0.07 | 0.00 | 0.03 |
| FlowSOM | Asinh | All | 100 | 1.00 | 1.00 | 0.77 | 0.77 |
| FlowSOM | Asinh | All | 400 | 0.38 | 1.00 | 0.38 | 0.45 |
| Phenograph, k=30 | Asinh | All | 47 | 0.40 | 0.70 | 0.31 | 0.31 |
| FAUST | Biexp | All | 125 | 0.01 | 0.02 | 0.01 | 0.01 |
| FlowSOM | Biexp | All | 100 | 1.00 | 1.00 | 0.73 | 0.73 |
| FlowSOM | Biexp | All | 400 | 0.51 | 0.56 | 0.28 | 0.28 |
| Phenograph, k=30 | Biexp | All | 49 | 1.00 | 1.00 | 0.34 | 0.34 |
| FAUST | Asinh | Baseline | 154 | 0.00 | 0.04 | 0.00 | 0.02 |
| FlowSOM | Asinh | Baseline | 100 | 1.00 | 1.00 | 0.68 | 0.68 |
| FlowSOM | Asinh | Baseline | 400 | 0.98 | 1.00 | 0.54 | 0.54 |
| Phenograph, k=30 | Asinh | Baseline | 46 | 0.76 | 0.80 | 0.35 | 0.35 |
| FAUST | Biexp | Baseline | 142 | 0.01 | 0.06 | 0.01 | 0.03 |
| FlowSOM | Biexp | Baseline | 100 | 0.48 | 1.00 | 0.48 | 0.70 |
| FlowSOM | Biexp | Baseline | 400 | 1.00 | 1.00 | 0.67 | 0.67 |
| Phenograph, k=30 | Biexp | Baseline | 45 | 0.59 | 1.00 | 0.36 | 0.36 |

Table S5: **Results of applying the clustering methods FlowSOM [6], and Phenograph [38], and FAUST to MCC anti-PD-1 trial T cell panel.** In all runs, tuning parameters for FlowSOM and Phenograph (including the number of clusters for FlowSOM) were set to the parameter settings reported in [5] (supporting table "cytoa23030-sup-0001-suppinfo.xlsx" entries for Phenograph and "FlowSOM pre"). Flow cytometry data are reported in [5] as being transformed by the hyperbolic arcsine transformation with cofactor 120. Here, we transform data using both the biexponential transformation (used by CITN) as well as the hyperbolic arcsine with cofactor 120 (The "Asinh" rows). Samples were concatenated before analysis by each method except FAUST, which was run at the sample level for all analyses. The reported FAUST number of clusters is the number of clusters with "CD3+" annotations. The tuning parameters for FAUST in the Asinh runs and the baseline Biexponential run were taken from the FAUST "Biexponential All" run (which is reported in the paper). FAUST parameters were transformed to the different scenarios from the settings underpinning Section 2.1 using empirical quantiles.

#### A.10 FAUST tuning parameter settings for data analysis

##### A.10.1 CITN-07 Phenotyping panel

The marker boundary matrix for CITN-07. Specific values were set by inspecting histograms of the individual markers across individual samples.

| | Low | High |
|---|---|---|
| CD123 | 1500.00 | 3000.00 |
| CD4 | 100.00 | 2500.00 |
| CD14 | 1.00 | 3000.00 |
| CD11C | 1.00 | 3000.00 |
| CD56 | 1.00 | 3000.00 |
| CD8 | 1.00 | 3000.00 |
| CD16 | 1000.00 | 3000.00 |
| CD3 | 1.00 | 3000.00 |
| CD122 | 1000.00 | 3000.00 |
| CD19 | 1501.00 | 3000.00 |
| HLA DR | 1.00 | 3500.00 |

The selection quantile was set to 0.5. The depth score threshold was set to 0.1. The supervised list was used to encourage CD4 to have two annotation boundaries estimated. The phenotype occurrence number was set to 70. The experimental unit was set to the individual sample. The imputation hierarchy was set to sample time point (with all follow-up appointments treated as a single time-point). The starting cell population was "45+" cells in the manual gating strategy.

##### A.10.2 CITN-09 T cell panel

The marker boundary matrix. Non-zero values were set by inspecting histograms of the individual samples.

| | Low | High |
|---|---|---|
| CD278 ICOS | -500.00 | 3000.00 |
| CD3 | -500.00 | 3000.00 |
| CD127 | -500.00 | 3000.00 |
| CD197 CCR7 | 500.00 | 2250.00 |
| CD279 PD1 | -250.00 | 3500.00 |
| CD8 | 500.00 | 3000.00 |
| CD4 | 500.00 | 3000.00 |
| CD28 | -500.00 | 3000.00 |
| CD25 | 500.00 | 2250.00 |
| HLA DR | -500.00 | 3000.00 |
| CD45RA | 1000.00 | 3000.00 |

The selection quantile was set to 0.05. The depth score threshold was set to The supervised list was used to encourage PD-1 to have two annotation boundaries estimated. The phenotype occurance number was set to 10. The experimental unit was set to the individual sample. The imputation hierarchy was to visit. The starting cell population was "L" cells (live lymphocytes) in the manual gating strategy.

##### A.10.3 CITN-09 Myeloid panel

The marker boundary matrix. Non-zero values were set by inspecting histograms of the individual samples.

| | 1_Low | 1_High | 2_Low | 2_High | 3_Low | 3_High | 4_Low | 4_High |
|---|---|---|---|---|---|---|---|---|
| CD11B | 1000 | Inf | -1500 | Inf | -1500 | Inf | 0 | Inf |
| CD20 | 1000 | Inf | -1000 | Inf | 0 | Inf | -2500 | Inf |
| CD14 | 1000 | Inf | 1000 | Inf | 500 | Inf | 1000 | Inf |
| CD11C | 1000 | Inf | -5000 | Inf | -5000 | Inf | 1000 | Inf |
| CD56 | 2000 | Inf | -2000 | Inf | -2000 | Inf | 2300 | Inf |
| CD33 | -1000 | 2100 | 2200 | Inf | 0 | Inf | -20000 | Inf |
| CD16 | -Inf | Inf | 1000 | Inf | 1000 | Inf | 1000 | Inf |
| CD3 | 1000 | Inf | -3000 | Inf | -5000 | Inf | 2000 | Inf |
| CD15 | 1000 | Inf | 1000 | Inf | 0 | Inf | 0 | Inf |
| CD19 | 1750 | Inf | -1000 | Inf | -1000 | Inf | -1000 | Inf |
| HLA DR | -1000 | 3750 | 1000 | 3750 | -500 | 3750 | -2500 | 3750 |

The selection quantile was set to 0.50. The depth score threshold was set to 0.01. The supervised list was used to encourage both CD33 and HLA-DR to have two annotation boundaries estimated, and CD15 one. The phenotype occurance number was set to 10. The experimental unit was set to the individual sample. The imputation hierarchy was set to four distinct values, with the marker boundary matrices of each value reported above. The starting cell population was "45+" cells in the manual gating strategy.

##### A.10.4 Krieg et al. CyTOF

The marker boundary matrix. Upper bounds were set to the $99^{th}$ quantiles across the dataset. Lower bounds were set to zero by default. Non-zero values were set by concatenating experimental samples together by batch, and inspecting histograms of the concatenated batches.

| | Low | High |
|---|---|---|
| 209Bi_CD11b | 0.00 | 5.55 |
| 162Dy_CD11c | 0.00 | 5.28 |
| 163Dy_CD7 | 0.00 | 4.85 |
| 166Er_CD209 | 1.00 | 3.00 |
| 167Er_CD38 | 0.00 | 4.10 |
| 151Eu_CD123 | 0.00 | 3.89 |
| 153Eu_CD62L | 0.00 | 4.57 |
| 152Gd_CD66b | 2.00 | 4.00 |
| 154Gd_ICAM-1 | 0.00 | 4.95 |
| 155Gd_CD1c | 0.00 | 2.30 |
| 156Gd_CD86 | 0.00 | 3.88 |
| 160Gd_CD14 | 0.00 | 4.33 |
| 165Ho_CD16 | 0.00 | 4.53 |
| 175Lu_PD-L1 | 0.00 | 3.73 |
| 146Nd_CD64 | 0.00 | 2.95 |
| 147Sm_CD303 | 2.00 | 4.00 |
| 148Sm_CD34 | 1.00 | 3.00 |
| 149Sm_CD141 | 0.00 | 3.96 |
| 150Sm_CD61 | 0.00 | 4.29 |
| 169Tm_CD33 | 0.00 | 3.68 |
| 89Y_CD45 | 0.00 | 5.25 |
| 173Yb_CD56 | 0.00 | 3.43 |
| 174Yb_HLA-DR | 0.00 | 5.89 |

The selection quantile was set to 1. The experimental unit was set to "batch": after concatenating by batch, there were two experimental units in this dataset. The imputation hierarchy was flat. The selection threshold was set to 0.05. The starting cell population was "root" cells, that consist of pre-gated CD3- CD19- cells.

##### A.10.5 Krieg et al. FACS

The marker boundary matrix. Non-zero values were set by inspecting histograms of the individual samples.

| | CD3 | CD4 | CD11b | CD33 | HLA-DR | CD56 | CD45RO | CD11c | CD16 | CD14 | CD19 |
|---|---|---|---|---|---|---|---|---|---|---|---|
| Low | -Inf | -20.00 | -Inf | -Inf | -Inf | -Inf | -Inf | -Inf | 85.00 | -Inf | -Inf |
| High | Inf | Inf | Inf | Inf | Inf | Inf | Inf | Inf | Inf | Inf | Inf |

The selection quantile was set to 0. The depth score threshold was set to 0.01. The supervised list was not used. The phenotype occurance number was set to 10. The experimental unit was set to the individual sample. The imputation hierarchy was flat. The starting cell population was "life" cells in the manual gating strategy.

##### A.10.6 Subrahmanyam et al. CyTOF

The marker boundary matrix. Non-zero values were set by inspecting histograms of the individual samples.

| | Low | High |
|---:|---:|---:|
| 145Nd_CD4 | 0.75 | 3.75 |
| 146Nd_CD8 | 1.00 | 4.00 |
| 141Pr_CD25 | 0.25 | 1.50 |
| 153Eu_CD45RA | 1.00 | 5.00 |
| 165Ho_CD127 | 0.00 | 5.32 |
| 169Tm_CCR7 | 0.00 | 2.75 |
| 154Sm_CD3 | 0.00 | 6.23 |
| 157Gd_HLA-DR | 2.00 | 6.25 |
| 170Er_PD-1 | 0.25 | 2.00 |
| 155Gd_CD28 | 0.25 | 4.54 |

The selection quantile was set to 1.0. The depth score threshold was set to 0.01. The supervised list was not used. The phenotype occurance number was not used, since FAUST was terminated after the annotation boundaries were standardized across samples. The experimental unit was set to the individual sample. The imputation hierarchy was flat. The starting cell population was "live" cells in the manual gating strategy recreated using openCyto.

#### A.11 Staining panels used in FAUST analyses

Staining panels from the experiments used in FAUST analyses are provided here.

##### A.11.1 CITN-09 T cell Staining Panel

| | name | desc |
|---|---|---|
| $P1 | FSC-A | |
| $P2 | FSC-H | |
| $P3 | SSC-A | |
| $P4 | SSC-H | |
| $P5 | <PE-A> | CD278 ICOS |
| $P6 | <FITC-A> | CD3 |
| $P7 | <BV 421-A> | CD127 |
| $P8 | <Alexa Fluor 700-A> | CD197 CCR7 |
| $P9 | <PE-Cy7-A> | CD279 PD-1 |
| $P10 | <PerCP-Cy5-5-A> | CD8 |
| $P11 | <APC-Cy7-A> | CD4 |
| $P12 | <ECD-A> | CD28 |
| $P13 | <APC-A> | CD25 |
| $P14 | PE-Cy5-A | |
| $P15 | <AmCyan-A> | CD45 |
| $P16 | <BV 605-A> | HLA DR |
| $P17 | <BV 650-A> | CD45RA |
| $P18 | Time | |

##### A.11.2 CITN-09 Myeloid Staining Panel

| | name | desc |
|---|---|---|
| $P1 | FSC-A | |
| $P2 | FSC-H | |
| $P3 | SSC-A | |
| $P4 | SSC-H | |
| $P5 | <PE-A> | CD11B |
| $P6 | <FITC-A> | CD20 |
| $P7 | <BV 421-A> | CD14 |
| $P8 | <Alexa Fluor 700-A> | CD11C |
| $P9 | <PE-Cy7-A> | CD56 |
| $P10 | <PerCP-Cy5-5-A> | CD33 |
| $P11 | <APC-Cy7-A> | CD16 |
| $P12 | <ECD-A> | CD3 |
| $P13 | <APC-A> | CD15 |
| $P14 | <PE-Cy5-A> | CD19 |
| $P15 | <AmCyan-A> | CD45 |
| $P16 | <BV 605-A> | HLA DR |
| $P17 | BV 650-A | |
| $P18 | Time | |

##### A.11.3 CITN-07 Phenotyping Staining Panel

| | name | desc |
|---|---|---|
| $P1 | FSC-A | |
| $P2 | FSC-H | |
| $P3 | SSC-A | |
| $P4 | SSC-H | |
| $P5 | <PE-A> | CD123 |
| $P6 | <FITC-A> | CD4 |
| $P7 | <BV 421-A> | CD14 |
| $P8 | <Alexa Fluor 700-A> | CD11C |
| $P9 | <PE-Cy7-A> | CD56 |
| $P10 | <PerCP-Cy5-5-A> | CD8 |
| $P11 | <APC-Cy7-A> | CD16 |
| $P12 | <ECD-A> | CD3 |
| $P13 | <APC-A> | CD122 |
| $P14 | <PE-Cy5-A> | CD19 |
| $P15 | <AmCyan-A> | CD45 |
| $P16 | <BV 605-A> | HLA DR |
| $P17 | BV 650-A | |
| $P18 | Time | |

##### A.11.4 Krieg et al. Myeloid CyTOF Panel

| | name | desc |
|---|---|---|
| $P1 | Bi209Di | 209Bi_CD11b |
| $P2 | Dy162Di | 162Dy_CD11c |
| $P3 | Dy163Di | 163Dy_CD7 |
| $P4 | Er166Di | 166Er_CD209 |
| $P5 | Er167Di | 167Er_CD38 |
| $P6 | Eu151Di | 151Eu_CD123 |
| $P7 | Eu153Di | 153Eu_CD62L |
| $P8 | Gd152Di | 152Gd_CD66b |
| $P9 | Gd154Di | 154Gd_ICAM-1 |
| $P10 | Gd155Di | 155Gd_CD1c |
| $P11 | Gd156Di | 156Gd_CD86 |
| $P12 | Gd160Di | 160Gd_CD14 |
| $P13 | Ho165Di | 165Ho_CD16 |
| $P16 | Lu175Di | 175Lu_PD-L1 |
| $P18 | Nd146Di | 146Nd_CD64 |
| $P22 | Sm147Di | 147Sm_CD303 |
| $P23 | Sm148Di | 148Sm_CD34 |
| $P24 | Sm149Di | 149Sm_CD141 |
| $P25 | Sm150Di | 150Sm_CD61 |
| $P26 | Tm169Di | 169Tm_CD33 |
| $P27 | Y89Di | 89Y_CD45 |
| $P29 | Yb173Di | 173Yb_CD56 |
| $P30 | Yb174Di | 174Yb_HLA-DR |

##### A.11.5 Krieg et al. FACS Panel

| | name | desc |
|---|---|---|
| $P1 | FSC-A | |
| $P2 | FSC-H | |
| $P3 | FSC-W | |
| $P4 | SSC-A | |
| $P5 | SSC-H | |
| $P6 | SSC-W | |
| $P7 | Comp-Brilliant Violet 785-A | CD3 |
| $P8 | Comp-Brilliant Violet 711-A | CD4 |
| $P9 | Comp-Brilliant Violet 421-A | CD11b |
| $P10 | Comp-PerCP-Cy5-5-A | CD33 |
| $P11 | Comp-FITC-A | HLA-DR |
| $P12 | Comp-PE-Cy7-A | CD56 |
| $P13 | Comp-PE-Texas Red-A | CD45RO |
| $P14 | Comp-APC-Cy7-A | NIR |
| $P15 | Comp-Alexa Fluor 700-A | CD11c |
| $P16 | Comp-APC-A | CD16 |
| $P17 | Comp-PE-A | CD14 |
| $P18 | Comp-Brilliant Violet 605-A | CD19 |
| $P19 | Time | |

##### A.11.6 Subrahmanyam et al. CyTOF Panel

| | name | desc |
|---|---|---|
| $P1 | Time | Time |
| $P2 | Event_length | Event_length |
| $P3 | 115In_Dead | 115In_Dead |
| $P4 | 140Ce_Beads | 140Ce_Beads |
| $P5 | 141Pr_CD25 | 141Pr_CD25 |
| $P6 | 142Nd_CD19 | 142Nd_CD19 |
| $P7 | 143Nd_IL-10 | 143Nd_IL-10 |
| $P8 | 144Nd_IL-4 | 144Nd_IL-4 |
| $P9 | 145Nd_CD4 | 145Nd_CD4 |
| $P10 | 146Nd_CD8 | 146Nd_CD8 |
| $P11 | 147Sm_CD20 | 147Sm_CD20 |
| $P12 | 148Nd_CD57 | 148Nd_CD57 |
| $P13 | 149Sm_CTLA-4 | 149Sm_CTLA-4 |
| $P14 | 150Nd_MIP-1b | 150Nd_MIP-1b |
| $P15 | 151Eu_CD107a | 151Eu_CD107a |
| $P16 | 152Sm_TNFa | 152Sm_TNFa |
| $P17 | 153Eu_CD45RA | 153Eu_CD45RA |
| $P18 | 154Sm_CD3 | 154Sm_CD3 |
| $P19 | 155Gd_CD28 | 155Gd_CD28 |
| $P20 | 156Gd_CD38 | 156Gd_CD38 |
| $P21 | 157Gd_HLA-DR | 157Gd_HLA-DR |
| $P22 | 158Gd_CD33 | 158Gd_CD33 |
| $P23 | 159Tb_GM-CSF | 159Tb_GM-CSF |
| $P24 | 160Gd_CD14 | 160Gd_CD14 |
| $P25 | 161Dy_IFNg | 161Dy_IFNg |
| $P26 | 162Dy_CD69 | 162Dy_CD69 |
| $P27 | 163Dy_TCRgd | 163Dy_TCRgd |
| $P28 | 164Dy_IL-17 | 164Dy_IL-17 |
| $P29 | 165Ho_CD127 | 165Ho_CD127 |
| $P30 | 166Er_IL-2 | 166Er_IL-2 |
| $P31 | 167Er_CD27 | 167Er_CD27 |
| $P32 | 168Er_CD154 | 168Er_CD154 |
| $P33 | 169Tm_CCR7 | 169Tm_CCR7 |
| $P34 | 170Er_PD-1 | 170Er_PD-1 |
| $P35 | 171Yb_GranzymeB | 171Yb_GranzymeB |
| $P36 | 172Yb_PD-L2 | 172Yb_PD-L2 |
| $P37 | 173Yb_Perforin | 173Yb_Perforin |
| $P38 | 174Yb_CD16 | 174Yb_CD16 |
| $P39 | 175Lu_PD-L1 | 175Lu_PD-L1 |
| $P40 | 176Yb_CD56 | 176Yb_CD56 |
| $P41 | 191Ir_DNA1 | 191Ir_DNA1 |
| $P42 | 193Ir_DNA2 | 193Ir_DNA2 |

#### 1037 A.12 Manual gating strategies

1038 Manual gating strategies for analyzed trials are included here.

##### 1039 A.12.1 CITN-09 T cell Manual Gating Strategy

| | |
|---|---|
| 1 | root |
| 2 | /Singlets |
| 3 | /Singlets/45 |
| 4 | /Singlets/45/Lymphocytes |
| 5 | /Singlets/45/Lymphocytes/CD3 |
| 6 | /Singlets/45/Lymphocytes/CD3/4 |
| 7 | /Singlets/45/Lymphocytes/CD3/4/CD25+ |
| 8 | /Singlets/45/Lymphocytes/CD3/4/CD25+CD45RA+CCR7+ |
| 9 | /Singlets/45/Lymphocytes/CD3/4/CD25+CD45RA+CCR7- |
| 10 | /Singlets/45/Lymphocytes/CD3/4/CD25+CD45RA-CCR7+ |
| 11 | /Singlets/45/Lymphocytes/CD3/4/CD25+CD45RA-CCR7- |
| 12 | /Singlets/45/Lymphocytes/CD3/4/CD25-CD45RA+CCR7+ |
| 13 | /Singlets/45/Lymphocytes/CD3/4/CD25-CD45RA+CCR7- |
| 14 | /Singlets/45/Lymphocytes/CD3/4/CD25-CD45RA-CCR7+ |
| 15 | /Singlets/45/Lymphocytes/CD3/4/CD25-CD45RA-CCR7- |
| 16 | /Singlets/45/Lymphocytes/CD3/4/CD28+ |
| 17 | /Singlets/45/Lymphocytes/CD3/4/CD28+CD45RA+CCR7+ |
| 18 | /Singlets/45/Lymphocytes/CD3/4/CD28+CD45RA+CCR7- |
| 19 | /Singlets/45/Lymphocytes/CD3/4/CD28+CD45RA-CCR7+ |
| 20 | /Singlets/45/Lymphocytes/CD3/4/CD28+CD45RA-CCR7- |
| 21 | /Singlets/45/Lymphocytes/CD3/4/28-CD45RA+CCR7+ |
| 22 | /Singlets/45/Lymphocytes/CD3/4/28-CD45RA+CCR7- |
| 23 | /Singlets/45/Lymphocytes/CD3/4/28-CD45RA-CCR7+ |
| 24 | /Singlets/45/Lymphocytes/CD3/4/28-CD45RA-CCR7- |
| 25 | /Singlets/45/Lymphocytes/CD3/4/CD45RA+ |
| 26 | /Singlets/45/Lymphocytes/CD3/4/278+ |

| | |
|---|---|
| 27 | /Singlets/45/Lymphocytes/CD3/4/CCR7+ |
| 28 | /Singlets/45/Lymphocytes/CD3/4/HLADR+ |
| 29 | /Singlets/45/Lymphocytes/CD3/4/PD1+ |
| 30 | /Singlets/45/Lymphocytes/CD3/4/CD45RA+ICOS+CCR7+ |
| 31 | /Singlets/45/Lymphocytes/CD3/4/CD45RA+ICOS+CCR7- |
| 32 | /Singlets/45/Lymphocytes/CD3/4/CD45RA+ICOS-CCR7+ |
| 33 | /Singlets/45/Lymphocytes/CD3/4/CD45RA+ICOS-CCR7- |
| 34 | /Singlets/45/Lymphocytes/CD3/4/CD45RA+CCR7+ |
| 35 | /Singlets/45/Lymphocytes/CD3/4/CD45RA+CCR7+HLADR+ |
| 36 | /Singlets/45/Lymphocytes/CD3/4/CD45RA+CCR7+HLADR- |
| 37 | /Singlets/45/Lymphocytes/CD3/4/CD45RA+CCR7+PD1+ |
| 38 | /Singlets/45/Lymphocytes/CD3/4/CD45RA+CCR7+PD1- |
| 39 | /Singlets/45/Lymphocytes/CD3/4/CD45RA+CCR7- |
| 40 | /Singlets/45/Lymphocytes/CD3/4/CD45RA+CCR7-HLADR+ |
| 41 | /Singlets/45/Lymphocytes/CD3/4/CD45RA+CCR7-HLADR- |
| 42 | /Singlets/45/Lymphocytes/CD3/4/CD45RA+CCR7-PD1+ |
| 43 | /Singlets/45/Lymphocytes/CD3/4/CD45RA+CCR7-PD1- |
| 44 | /Singlets/45/Lymphocytes/CD3/4/CD45RA-ICOS+CCR7+ |
| 45 | /Singlets/45/Lymphocytes/CD3/4/CD45RA-ICOS+CCR7- |
| 46 | /Singlets/45/Lymphocytes/CD3/4/CD45RA-ICOS-CCR7+ |
| 47 | /Singlets/45/Lymphocytes/CD3/4/CD45RA-ICOS-CCR7- |
| 48 | /Singlets/45/Lymphocytes/CD3/4/CD45RA-CCR7+ |
| 49 | /Singlets/45/Lymphocytes/CD3/4/CD45RA-CCR7+HLADR+ |
| 50 | /Singlets/45/Lymphocytes/CD3/4/CD45RA-CCR7+HLADR- |
| 51 | /Singlets/45/Lymphocytes/CD3/4/CD45RA-CCR7+PD1+ |
| 52 | /Singlets/45/Lymphocytes/CD3/4/CD45RA-CCR7+PD1- |
| 53 | /Singlets/45/Lymphocytes/CD3/4/CD45RA-CCR7- |
| 54 | /Singlets/45/Lymphocytes/CD3/4/CD45RA-CCR7-HLADR+ |
| 55 | /Singlets/45/Lymphocytes/CD3/4/CD45RA-CCR7-HLADR- |
| 56 | /Singlets/45/Lymphocytes/CD3/4/CD45RA-CCR7-PD1+ |

| | |
|---|---|
| 57 | /Singlets/45/Lymphocytes/CD3/4/CD45RA-CCR7-PD1- |
| 58 | /Singlets/45/Lymphocytes/CD3/4/CD45RA- |
| 59 | /Singlets/45/Lymphocytes/CD3/4/treg |
| 60 | /Singlets/45/Lymphocytes/CD3/8 |
| 61 | /Singlets/45/Lymphocytes/CD3/8/CD25+ |
| 62 | /Singlets/45/Lymphocytes/CD3/8/CD25+CD45RA+CCR7+ |
| 63 | /Singlets/45/Lymphocytes/CD3/8/CD25+CD45RA+CCR7- |
| 64 | /Singlets/45/Lymphocytes/CD3/8/CD25+CD45RA-CCR7+ |
| 65 | /Singlets/45/Lymphocytes/CD3/8/CD25+CD45RA-CCR7- |
| 66 | /Singlets/45/Lymphocytes/CD3/8/CD25-CD45RA+CCR7+ |
| 67 | /Singlets/45/Lymphocytes/CD3/8/CD25-CD45RA+CCR7- |
| 68 | /Singlets/45/Lymphocytes/CD3/8/CD25-CD45RA-CCR7+ |
| 69 | /Singlets/45/Lymphocytes/CD3/8/CD25-CD45RA-CCR7- |
| 70 | /Singlets/45/Lymphocytes/CD3/8/CD28+ |
| 71 | /Singlets/45/Lymphocytes/CD3/8/CD28+CD45RA+CCR7+ |
| 72 | /Singlets/45/Lymphocytes/CD3/8/CD28+CD45RA+CCR7- |
| 73 | /Singlets/45/Lymphocytes/CD3/8/CD28+CD45RA-CCR7+ |
| 74 | /Singlets/45/Lymphocytes/CD3/8/CD28+CD45RA-CCR7- |
| 75 | /Singlets/45/Lymphocytes/CD3/8/28-CD45RA+CCR7+ |
| 76 | /Singlets/45/Lymphocytes/CD3/8/28-CD45RA+CCR7- |
| 77 | /Singlets/45/Lymphocytes/CD3/8/28-CD45RA-CCR7+ |
| 78 | /Singlets/45/Lymphocytes/CD3/8/28-CD45RA-CCR7- |
| 79 | /Singlets/45/Lymphocytes/CD3/8/CD45RA+ |
| 80 | /Singlets/45/Lymphocytes/CD3/8/278+ |
| 81 | /Singlets/45/Lymphocytes/CD3/8/CCR7+ |
| 82 | /Singlets/45/Lymphocytes/CD3/8/HLADR+ |
| 83 | /Singlets/45/Lymphocytes/CD3/8/PD1+ |
| 84 | /Singlets/45/Lymphocytes/CD3/8/CD45RA+ICOS+CCR7+ |
| 85 | /Singlets/45/Lymphocytes/CD3/8/CD45RA+ICOS+CCR7- |
| 86 | /Singlets/45/Lymphocytes/CD3/8/CD45RA+ICOS-CCR7+ |

| | |
|---|---|
| 87 | /Singlets/45/Lymphocytes/CD3/8/CD45RA+ICOS-CCR7- |
| 88 | /Singlets/45/Lymphocytes/CD3/8/CD45RA+CCR7+ |
| 89 | /Singlets/45/Lymphocytes/CD3/8/CD45RA+CCR7+HLADR+ |
| 90 | /Singlets/45/Lymphocytes/CD3/8/CD45RA+CCR7+HLADR- |
| 91 | /Singlets/45/Lymphocytes/CD3/8/CD45RA+CCR7+PD1+ |
| 92 | /Singlets/45/Lymphocytes/CD3/8/CD45RA+CCR7+PD1- |
| 93 | /Singlets/45/Lymphocytes/CD3/8/CD45RA+CCR7- |
| 94 | /Singlets/45/Lymphocytes/CD3/8/CD45RA+CCR7-HLADR+ |
| 95 | /Singlets/45/Lymphocytes/CD3/8/CD45RA+CCR7-HLADR- |
| 96 | /Singlets/45/Lymphocytes/CD3/8/CD45RA+CCR7-PD1+ |
| 97 | /Singlets/45/Lymphocytes/CD3/8/CD45RA+CCR7-PD1- |
| 98 | /Singlets/45/Lymphocytes/CD3/8/CD45RA-ICOS+CCR7+ |
| 99 | /Singlets/45/Lymphocytes/CD3/8/CD45RA-ICOS+CCR7- |
| 100 | /Singlets/45/Lymphocytes/CD3/8/CD45RA-ICOS-CCR7+ |
| 101 | /Singlets/45/Lymphocytes/CD3/8/CD45RA-ICOS-CCR7- |
| 102 | /Singlets/45/Lymphocytes/CD3/8/CD45RA-CCR7+ |
| 103 | /Singlets/45/Lymphocytes/CD3/8/CD45RA-CCR7+HLADR+ |
| 104 | /Singlets/45/Lymphocytes/CD3/8/CD45RA-CCR7+HLADR- |
| 105 | /Singlets/45/Lymphocytes/CD3/8/CD45RA-CCR7+PD1+ |
| 106 | /Singlets/45/Lymphocytes/CD3/8/CD45RA-CCR7+PD1- |
| 107 | /Singlets/45/Lymphocytes/CD3/8/CD45RA-CCR7- |
| 108 | /Singlets/45/Lymphocytes/CD3/8/CD45RA-CCR7-HLADR+ |
| 109 | /Singlets/45/Lymphocytes/CD3/8/CD45RA-CCR7-HLADR- |
| 110 | /Singlets/45/Lymphocytes/CD3/8/CD45RA-CCR7-PD1+ |
| 111 | /Singlets/45/Lymphocytes/CD3/8/CD45RA-CCR7-PD1- |
| 112 | /Singlets/45/Lymphocytes/CD3/8/CD45RA- |

Table S6: Manual gating strategy applied to CITN-09 T cell panel.

##### A.12.2 CITN-09 Myeloid Manual Gating Strategy

| | |
|---|---|
| 1 | root |
| 2 | /Singlets |
| 3 | /Singlets/45+ |
| 4 | /Singlets/45+/CD3-CD19- |
| 5 | /Singlets/45+/CD3-CD19-/CD20- |
| 6 | /Singlets/45+/CD3-CD19-/CD20-/CD56- |
| 7 | /Singlets/45+/CD3-CD19-/CD20-/CD56-/HLADR- |
| 8 | /Singlets/45+/CD3-CD19-/CD20-/CD56-/HLADR-/CD16- |
| 9 | /Singlets/45+/CD3-CD19-/CD20-/CD56-/HLADR-/CD16-/CD14+ |
| 10 | /Singlets/45+/CD3-CD19-/CD20-/CD56-/HLADR-/CD16-/CD14+/Q1: CD33-, CD11B+ |
| 11 | /Singlets/45+/CD3-CD19-/CD20-/CD56-/HLADR-/CD16-/CD14+/Q2: CD33+, CD11B+ (m-MDSC) |
| 12 | /Singlets/45+/CD3-CD19-/CD20-/CD56-/HLADR-/CD16-/CD14+/Q3: CD33+, CD11B- |
| 13 | /Singlets/45+/CD3-CD19-/CD20-/CD56-/HLADR-/CD16-/CD14+/Q4: CD33-, CD11B- |
| 14 | /Singlets/45+/CD3-CD19-/CD20-/CD56-/HLADR-/CD16-/CD14-CD15+ |
| 15 | /Singlets/45+/CD3-CD19-/CD20-/CD56-/HLADR-/CD16-/CD14-CD15+/Q1: CD33-, CD11B+ |
| 16 | /Singlets/45+/CD3-CD19-/CD20-/CD56-/HLADR-/CD16-/CD14-CD15+/Q2: CD33+, CD11B+ (PMN-MDSC) |
| 17 | /Singlets/45+/CD3-CD19-/CD20-/CD56-/HLADR-/CD16-/CD14-CD15+/Q3: CD33+, CD11B- |
| 18 | /Singlets/45+/CD3-CD19-/CD20-/CD56-/HLADR-/CD16-/CD14-CD15+/Q4: CD33-, CD11B- |
| 19 | /Singlets/45+/CD3-CD19-/CD20-/CD56-/HLADR-/CD16-/CD14-CD15- |
| 20 | /Singlets/45+/CD3-CD19-/CD20-/CD56-/HLADR-/CD16-/CD14-CD15-/Q1: CD33-, CD11B+ |

| | |
|---|---|
| 21 | /Singlets/45+/CD3-CD19-/CD20-/CD56-/HLADR-/CD16-/CD14-CD15-/Q2: CD33+, CD11B+ (e-MDSC) |
| 22 | /Singlets/45+/CD3-CD19-/CD20-/CD56-/HLADR-/CD16-/CD14-CD15-/Q3: CD33+, CD11B- |
| 23 | /Singlets/45+/CD3-CD19-/CD20-/CD56-/HLADR-/CD16-/CD14-CD15-/Q4: CD33-, CD11B- |
| 24 | /Singlets/45+/CD14+ |
| 25 | /Singlets/45+/CD14-CD15+ |
| 26 | /Singlets/45+/CD14-CD15- |

Table S7: Manual gating strategy applied to CITN-09 Myeloid panel.

##### A.12.3 CITN-07 Phenotyping Manual Gating Strategy

| | |
|---|---|
| 1 | root |
| 2 | /Beads |
| 3 | /Non-beads |
| 4 | /Non-beads/Singlets |
| 5 | /Non-beads/Singlets/45+ |
| 6 | /Non-beads/Singlets/45+/14+ |
| 7 | /Non-beads/Singlets/45+/14- |
| 8 | /Non-beads/Singlets/45+/14-/3-19- |
| 9 | /Non-beads/Singlets/45+/14-/3-19-/56-16- |
| 10 | /Non-beads/Singlets/45+/14-/3-19-/56-16-/Basophils |
| 11 | /Non-beads/Singlets/45+/14-/3-19-/56-16-/Basophils/HLA DR hi |
| 12 | /Non-beads/Singlets/45+/14-/3-19-/56-16-/Basophils/HLA DR med |
| 13 | /Non-beads/Singlets/45+/14-/3-19-/56-16-/Basophils/HLA DR neg |
| 14 | /Non-beads/Singlets/45+/14-/3-19-/56-16-/HLADR+ |
| 15 | /Non-beads/Singlets/45+/14-/3-19-/56-16-/HLADR+/mDC |
| 16 | /Non-beads/Singlets/45+/14-/3-19-/56-16-/HLADR+/mDC/HLADRhi |
| 17 | /Non-beads/Singlets/45+/14-/3-19-/56-16-/HLADR+/mDC/HLADRmed |
| 18 | /Non-beads/Singlets/45+/14-/3-19-/56-16-/HLADR+/mDC/HLA DR hi |
| 19 | /Non-beads/Singlets/45+/14-/3-19-/56-16-/HLADR+/mDC/HLA DR med |
| 20 | /Non-beads/Singlets/45+/14-/3-19-/56-16-/HLADR+/mDC/HLA DR neg |
| 21 | /Non-beads/Singlets/45+/14-/3-19-/56-16-/HLADR+/pDC |
| 22 | /Non-beads/Singlets/45+/14-/3-19-/56-16-/HLADR+/pDC/HLA DR hi |
| 23 | /Non-beads/Singlets/45+/14-/3-19-/56-16-/HLADR+/pDC/HLA DR med |
| 24 | /Non-beads/Singlets/45+/14-/3-19-/56-16-/HLADR+/pDC/HLA DR neg |
| 25 | /Non-beads/Singlets/45+/Lymphocytes |
| 26 | /Non-beads/Singlets/45+/Lymphocytes/3+ |
| 27 | /Non-beads/Singlets/45+/Lymphocytes/3+/4&8 |
| 28 | /Non-beads/Singlets/45+/Lymphocytes/3+/4&8/56+ |
| 29 | /Non-beads/Singlets/45+/Lymphocytes/3+/4&8/122+ |

| | |
|---|---|
| 30 | /Non-beads/Singlets/45+/Lymphocytes/3+/4&8++ |
| 31 | /Non-beads/Singlets/45+/Lymphocytes/3+/4+ |
| 32 | /Non-beads/Singlets/45+/Lymphocytes/3+/4+/HLADR+ |
| 33 | /Non-beads/Singlets/45+/Lymphocytes/3+/8+ |
| 34 | /Non-beads/Singlets/45+/Lymphocytes/3+/8+/HLADR+ |
| 35 | /Non-beads/Singlets/45+/Lymphocytes/3-19- |
| 36 | /Non-beads/Singlets/45+/Lymphocytes/3-19-/16+56- |
| 37 | /Non-beads/Singlets/45+/Lymphocytes/3-19-/56+ |
| 38 | /Non-beads/Singlets/45+/Lymphocytes/3-19-/56+/122+ |
| 39 | /Non-beads/Singlets/45+/Lymphocytes/3-19-/56+16- |
| 40 | /Non-beads/Singlets/45+/Lymphocytes/3-19-/56-16- |
| 41 | /Non-beads/Singlets/45+/Lymphocytes/3-19-/56-16-/122+ |
| 42 | /Non-beads/Singlets/45+/Lymphocytes/3-19-/56-16-/HLA DR hi |
| 43 | /Non-beads/Singlets/45+/Lymphocytes/3-19-/56-16-/HLA DR med |
| 44 | /Non-beads/Singlets/45+/Lymphocytes/3-19-/56-16-/HLA DR neg |
| 45 | /Non-beads/Singlets/45+/Lymphocytes/3-19-/56B |
| 46 | /Non-beads/Singlets/45+/Lymphocytes/3-19-/56B/HLA DR hi |
| 47 | /Non-beads/Singlets/45+/Lymphocytes/3-19-/56B/HLA DR med |
| 48 | /Non-beads/Singlets/45+/Lymphocytes/3-19-/56B/HLA DR neg |
| 49 | /Non-beads/Singlets/45+/Lymphocytes/3-19-/56B16- |
| 50 | /Non-beads/Singlets/45+/Lymphocytes/3-19-/56D |
| 51 | /Non-beads/Singlets/45+/Lymphocytes/3-19-/56D/HLA DR hi |
| 52 | /Non-beads/Singlets/45+/Lymphocytes/3-19-/56D/HLA DR med |
| 53 | /Non-beads/Singlets/45+/Lymphocytes/3-19-/56D/HLA DR neg |
| 54 | /Non-beads/Singlets/45+/Lymphocytes/19+ |
| 55 | /Non-beads/Singlets/45+/Lymphocytes/19+/B CELLS |
| 56 | /Non-beads/Singlets/45+/Lymphocytes/19+/B CELLS/HLA DR hi |
| 57 | /Non-beads/Singlets/45+/Lymphocytes/19+/B CELLS/HLA DR med |
| 58 | /Non-beads/Singlets/45+/Lymphocytes/19+/B CELLS/HLA DR neg |

Table S8: Manual gating strategy applied to CITN-07 phenotyping panel.

#### A.13 Simulation study

#### A.14 Summary of simulation results

To better understand the performance of FAUST, we conducted simulation studies that generated data from a variety of mixture models. Since FAUST assumes that each experimental unit is sampled from a finite mixture model (see Methods 4), all datasets generated in the study were designed to be compatible with the statistical assumptions underpinning FAUST. Components of the mixture for each experimental unit are assumed to arise from a common class of densities, with batch effects and other sources of experimental heterogeneity modeled as unit-specific changes in location and scale of the underlying mixture components. The mixture components represent cell sub-populations within a unit.

The study generated datasets from a variety of mixture models incorporating different combinations of assumptions, detailed in the following sub-sections. The study begins by simulating data from multivariate Gaussian distributions (producing datasets which are favorable to many existing methods) and progressively simulates data that more closely represents flow cytometry and CyTOF datasets. In the study, we compare FAUST to FlowSOM since, as noted in the main text, FlowSOM is computationally efficient, is recommended in the review [5]. Each simulated mixture component (representing a cell sub-population) is partially parameterized by a mean vector and is given a phenotypic label that describes the phenotype of the component. By treating these phenotypic labels as ground-truth, we are able to measure how well the count matrix produced by FAUST agrees with the simulated count matrix, matching discovered and simulated cell populations based on their phenotypes. FAUST is run completely unsupervised across all simulation settings.

Our results demonstrate that FAUST's discovery and annotation strategy does not severely over partition the data under a variety of generative regimes (supplementary figure S6). Results also show that the cell counts derived from FAUST's discovered clusters strongly correlate with the underlying true counts across all simulation settings. We observe a median correlation of 0.917 between FAUST and the simulated truth, when cluster counts are correlated between FAUST clusters and the ground truth using only cluster annotations to perform the comparison (Supplementary Figure S5).

The simulated datasets always include a sub-population that is differentially abundant between

50% of the subjects. Our results show that when we simulate a causal relationship of varying strength between this differential sub-population and a simulated response to therapy, FAUST discovers the differential sub-population, annotates it correctly, and often identifies that the differential population is associated with response to therapy (Supplementary Figures S7, S8, S9).

In the present simulation, FlowSOM clusters are tested for differential abundance under the same causal regimes as FAUST. Our results show that FlowSOM's ability to detect the causal association is adversely affected when the simulation departs from multivariate normality or when the simulated data contains 50 true clusters and batch effects and/or nuisance variables, even when FlowSOM is provided with the true number of clusters as a tuning parameter.

##### A.14.1 Simulation Goals

The purpose of this simulation study is to assess the performance of the FAUST algorithm, both as a clustering tool and as a discovery tool. Datasets are simulated from mixture models following the assumptions of section 4.1. The simulation measures how well FAUST recovers the underlying mixture under a variety of parametric scenarios. The simulation also measures how well FAUST is able to detect a sub-population, elevated in half the samples, that is required to have causal relationship (of varying strength) with a subject's response to therapy. We compare the performance of FAUST to the performance of the FlowSOM clustering algorithm [6].

##### A.14.2 Baseline simulation description

The basic simulation generates an experimental data collection containing 100 independent samples of 10-dimensional data from a Gaussian mixture model with 10 components. A probability vector

$$\mathbf{p} \sim \text{Dirichlet}\left(\alpha \equiv (1, 1, \ldots, 1)\right) \tag{A.7}$$

of dimension equal to the number of mixture components is generated. In a given simulation iteration, sampling from the Dirichlet continues until all elements are greater or equal to 0.001.

There are four tuning parameters that modify this baseline setting. We will first give a complete description of how the simulation study works at baseline, and then will describe how the tuning parameters modify the baseline study.

In the basic setting, the size of each of the 100 samples is $n_j = \max(5000, s)$, $1 \leq j \leq 100$, where

$s \sim T(\mu = 10000, \nu = 3)$ is a sample from a non-central $T$ distribution with 3 degrees of freedom and non-centrality parameter 10000. Each sample is meant to represent a sample taken from a subject in an immunology study and then interrogated via flow cytometry.

Before generating the samples, a fixed collection of mean vectors $\mu_c$, $1 \leq c \leq 10$ is determined for the ten Gaussian mixture components that is used across all simulated samples. Each of the ten entries of $\mu_c$ are randomly selected from the columns of table S9, and represent whether or not the measured variable exhibits a signal. When an entry of $\mu_c$ is from the "No Signal" row of table S9, the corresponding variable is labeled "-". Similarly, when an entry of $\mu_c$ is from the "Signal" row of table S9, the corresponding variable is labeled "+". An an example, the annotation "V1- V2- V3+ V4- V5+ V6- V7- V8- V9+ V10-" indicates the mean vector $\mu_c$ of the mixture component contains 0 for V1, V2, V4, V6, V7, V8, and V10, while it is 7 for V3, 6 for V5, and 4 for V9. Each mean vector is associated with an element of the probability vector (A.7). Covariance matrices $\Sigma_c$ are always constrained to have variances between 1 and 2, but otherwise are randomly generated sample-by-sample and component-by-component.

Table S9: Possible mean vector entries for the ten simulation variables.

| | V1 | V2 | V3 | V4 | V5 | V6 | V7 | V8 | V9 | V10 |
|---|---|---|---|---|---|---|---|---|---|---|
| No Signal | 0 | 0 | 0 | 0 | 0 | 0 | 0 | 0 | 0 | 0 |
| Signal | 8 | 8 | 7 | 7 | 6 | 6 | 5 | 5 | 4 | 4 |

Each simulation iteration, 50 of the 100 samples are randomly selected to have a mixture component elevated. Without loss of generality, suppose (A.7) is in sorted order, so that the first entry $p_1$ is the largest value, the tenth entry $p_{10}$ is the smallest value, and intermediate entries correspond to their order statistics. In the non-elevated samples, the mean-vector $\mu_c$ associated with the smallest element of the probability vector (A.7), $p_{10}$, is identified as the cluster component to elevate. In the samples randomly selected for elevation, the probability vector (A.7) is modified as follows. The numerical value $p_{\text{target}} \equiv p_7$ is fixed. Next, the intermediate probability vector

$$\begin{aligned}\mathbf{p_{int}} &\equiv \left(p_1 + \frac{p_{10}}{9}, p_2 + \frac{p_{10}}{9}, \ldots, p_9 + \frac{p_{10}}{9}, 0\right) \\ &\equiv (q_1, q_2, \ldots, q_9, 0) \qquad \text{(A.8)}\end{aligned}$$

is generated. Then (A.8) is modified so that

$$\begin{aligned} \mathbf{p}_{\textbf{elevated}} &\equiv \left(q_1 - q_1 \cdot p_{\text{target}}, q_2 - q_2 \cdot p_{\text{target}}, \ldots, q_9 - q_9 \cdot p_{\text{target}}, p_{\text{target}}\right) \\ &\equiv \left(r_1, r_2, \ldots, r_9, r_{10}\right) . \end{aligned} \tag{A.9}$$

The transformation from (A.7) to (A.9) causes the identified population to be, on average, the $7^{th}$ largest mixture component in half the samples, and the smallest mixture component in the other half.

A sample of size $n_j$ with $1 \leq j \leq 100$ is generated by first determining the relative size of each mixture component within the sample. When the sample is selected as having the elevated population, the size of mixture components is determined by taking a sample from a multinomial distribution with $n_j$ trials and cell probabilities determined by (A.9). Otherwise, the size of mixture components is determined by taking a sample from a multinomial distribution with $n_j$ trials and cell probabilities determined by (A.7). In both cases, the resulting multinomial vector is then used to sample multivariate Gaussian samples of the corresponding size, with mean vectors $\mu_c + e_{c,j}$ and covariance matrices $\Sigma_c$, for $1 \leq c \leq 10$. The vector $e_{c,j} = (e_{c,j,1}, \ldots, e_{c,j,10})$ is determined by taking a 10 independent samples $\epsilon_{c,j,k} \sim N(0, 1/2)$, $1 \leq k \leq 10$, and then rounding $e_{c,j,k} = round(\epsilon_{c,j,k})$ to the nearest integer. The vector $e_{c,j}$ models sample-specific perturbations (corresponding to subject-level effects) without modifying (with high probability) the semantic interpretation of the annotations corresponding to $\mu_c$.

Once the experimental data is generated, it is processed by FAUST in a completely unsupervised setting. FAUST is set to use individual samples as the experimental unit. All simulated variables are taken as admissible and the channel boundaries are set to the entire real line for all markers. The depth score selection threshold is set to 0.01, the depth score selection quantile is set to the median, and the phenotype occurence number is set to 25. The 100 samples are also concatenated and clustered by the FlowSOM algorithm in two different ways. First, following the recommendation of [49], the FlowSOM grid is set to $1 \times$ Number of mixture components to simulate one best case scenario: an oracle provides FlowSOM with the true number of clusters. Second, similar to the approach of [19], FlowSOM overpartitions the data by setting the grid to $5 \times 5$ (assuming 25 clusters when in truth there are 10).

To test how well each of the three methods discover sub-populations associated with differential abundance, a binary response is generated for each sample in the experiment. For samples where the identified population is elevated, a probability of response $p_{response}$ is varied from 0.50 to 0.80 in increments of 0.05. Each elevated sample is then associated with a response status by sampling from a Bernoulli$(p_{response})$. Similarly, samples where the identified population is not elevated are given a probability of response $q_{response} \equiv 1 - p_{response}$. Each non-elevated sample is then associated with a response status by sampling from a Bernoulli$(q_{response})$.

Once samples are associated with a binary outcome, the clusters produced by each of the three approaches are tested for differential abundance following the strategy described in section A equation (4.5). P-values are adjusted for FDR (q-values) using the method [50]. In the event FAUST discovers the elevated population by exact annotation, the associate q-value is recorded. For FlowSOM, the "best" q-value is defined as follows. Both the cluster containing the largest number of observations from the elevated population in terms of absolute counts, and the cluster containing proportionally the most observation from the elevated population are identified. The minimum q-value from the two clusters (when different) is recorded for both the oracle FlowSOM and overpartitioned FlowSOM clusterings.

We repeat this modeling procedure 50 times for each setting of $p_{response}$. The median q-values across each of the 50 iterations is recorded in a single simulation iteration. We then repeat the entire experimental simulation 50 times, and report the median of median q-values across those 50 simulation runs. In addition, we compute F-measures of the clusterings, along with several other measures of the quality of the FAUST clusterings. We will describe these measurements in the coming figures. Before doing so, we will provide details about simulation tuning parameters.

##### A.14.3 Simulation tuning parameters

The first simulation parameter we vary is the underlying number of mixture components: we set this parameter to 25 components and 50 components, in addition to the baseline of 10. While the sample sizes are random, we do not change the underlying sampling scheme, which introduces rarer and rarer populations appear across simulations as the number of mixture components increase. In both the 25 and 50 component setting, the probability vector (A.7) is expanded accordingly; $\alpha$ continues to be set to 1 for each component. In all cases, sampling from the Dirichlet continues until all elements are greater or equal to 0.001. In the 25 component setting, the elevated population has $p_{\text{target}}$ set to $p_{18}$; in the 50 component setting, $p_{\text{target}}$ is set to $p_{35}$.

The second simulation parameter we vary is used to add a batch effect to the simulation. The batch effect is modeled as a translation of the underlying mean vector. Batches are modeled as groups of 10 samples. After the initial 10 samples are generated, the mean vectors of the Gaussian mixtue components (sampled from (S9)) are translated by a constant vector $\lambda_1 = (1/3, 1/3, \ldots, 1/3)$. After the next 10 samples are generated, the translate increases to the constant vector $\lambda_2 = (2/3, 2/3, \ldots, 2/3)$. This continues in groups of 10 until the final 10 samples are translated by $\lambda_9 = (9/3, 9/3, \ldots, 9/3)$.

The third simulation parameter controls whether or not we add nuisance variables to the simulation. This parameter is meant to generate data under the scenario that several markers in the panel are uninformative because of staining issues. When this parameter is turned on the following occurs. Each time a sample of size $n_j$ is generate, an independent sample of size $n_j$ it taken from a Multivariate Gaussian distribution centered at $\mu_{\text{nuisance}} = (5, 5, 5, 5, 5)$, and $\Sigma_{\text{nuisance}}$ constrained to have variances between 1 and 2 but otherwise random. The independent Gaussian sample is then adjoined to the mixture of size $n_j$, producing a simulated dataset in 15 dimensions. Since nuisance variables are independently generated, they do not affect the mixture structure of a given simulation; consequently, Consequently, the underlying annotations of observations by their cluster component mean vector are not changed when the nuisance variables are added to the simulation.

The final simulation parameter is used to investigate departures from normality. We explored two possible settings: after generating each sample, the data are transformed coordinate-by-coordinate through the square map $f(x) = x^2$ or the gamma map $g(x) = \Gamma(1 + |(|x/4))$. The square map was used to investigate a mild departure from Normality, while we used the gamma map to transform the mixture into data that looked similar to CyTOF. Under the gamma map, we modify the space possible Gaussian mean vectors (S9) to those determined by table (S10).

Table S10: Possible mean vector entries for the ten simulation variables when data subsequently transformed by the map $g(x) = \Gamma(1 + |(|x/4))$.

| | V1 | V2 | V3 | V4 | V5 | V6 | V7 | V8 | V9 | V10 |
|---|---|---|---|---|---|---|---|---|---|---|
| No Signal | 0 | 0 | 0 | 0 | 0 | 0 | 0 | 0 | 0 | 0 |
| Signal | 8 | 8 | 8 | 7 | 7 | 7 | 7 | 6 | 6 | 6 |

##### A.14.4 Simulation results

By adjusting the tuning parameters described in supplementary section A.14.3, we explore 36 distinct scenarios *in silico*. Each simulation setting is run 25 times.

This simulation study shows that departures from multivariate-normality as well as batch-effects combined with large numbers of clusters impair FlowSOM's ability to define clusters that correlate with outcome. FAUST, on the other hand, performed robustly across simulation settings since its key methodological assumption is that some subset of the measured markers in a cytometry dataset are marginally separated into modal groups. In samples that both contain heterogeneous cell populations (such as live lymphocytes) and are stained by a large marker panel, we have empirically seen this is assumption is always met. Plots of the observed expression data show the MCC anti-PD1 dataset has non-Gaussian characteristics, and also has sample-to-sample variation which is common in many cytometry experiments. Hence, the non-Gaussian nature of the MCC anti-PD1 trial data combined with sample-to-sample variation both contribute to the discovery differences observed between FlowSOM and FAUST.

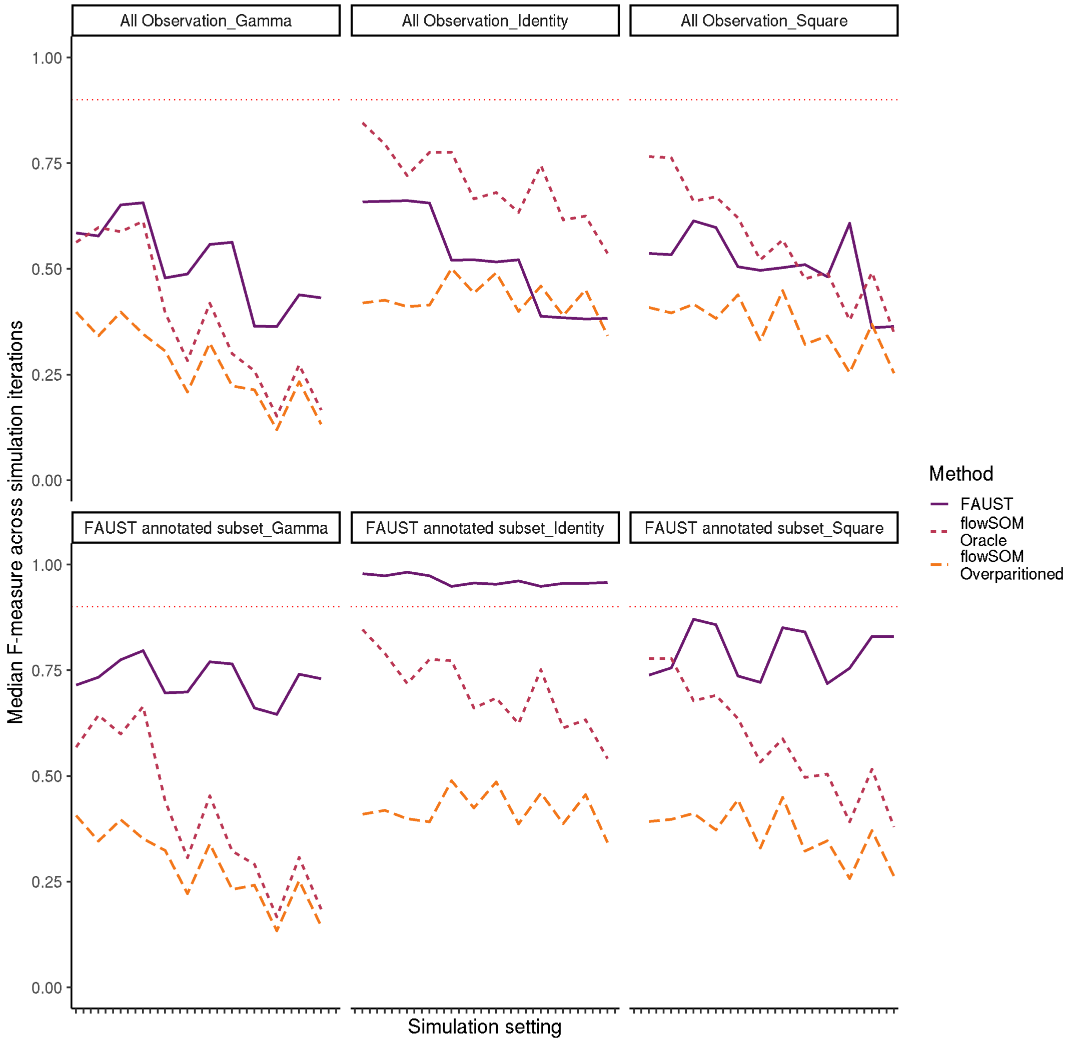


Figure S4: The median F-measure across the 50 simulation iterations (with 3 exceptions). Results are stratified by transformation type: $h(x) = x$ (identity map); $f(x) = x^2$ (square map); $g(x) = \Gamma(1 + |(|x/4))$ (gamma map). F-measures are computed between each method's clustering and the entire simulated dataset (row 1). F-measures are also computed between each method and the subset of observations that FAUST annotates (row 2). The figures show FAUST improves markedly (in terms of F-measure) on the set of labeled observations it labels, while the F-measure of FlowSOM with an oracle and FlowSOM overpartitioned perform similarly on the two sets. This figure provides a demonstration of the difficulty of comparing FAUST clusterings to computational methods in current use: classic measures of clustering performance, such as the F-measure, do not directly account for the biological information present in FAUST annotations. When the annotated subset is compared to associated subset of the ground truth, FAUST's performance improves markedly in terms of F-measure. FlowSOM's performance on the annotated subset is similar to its performance on the entire simulated dataset across simulation settings.

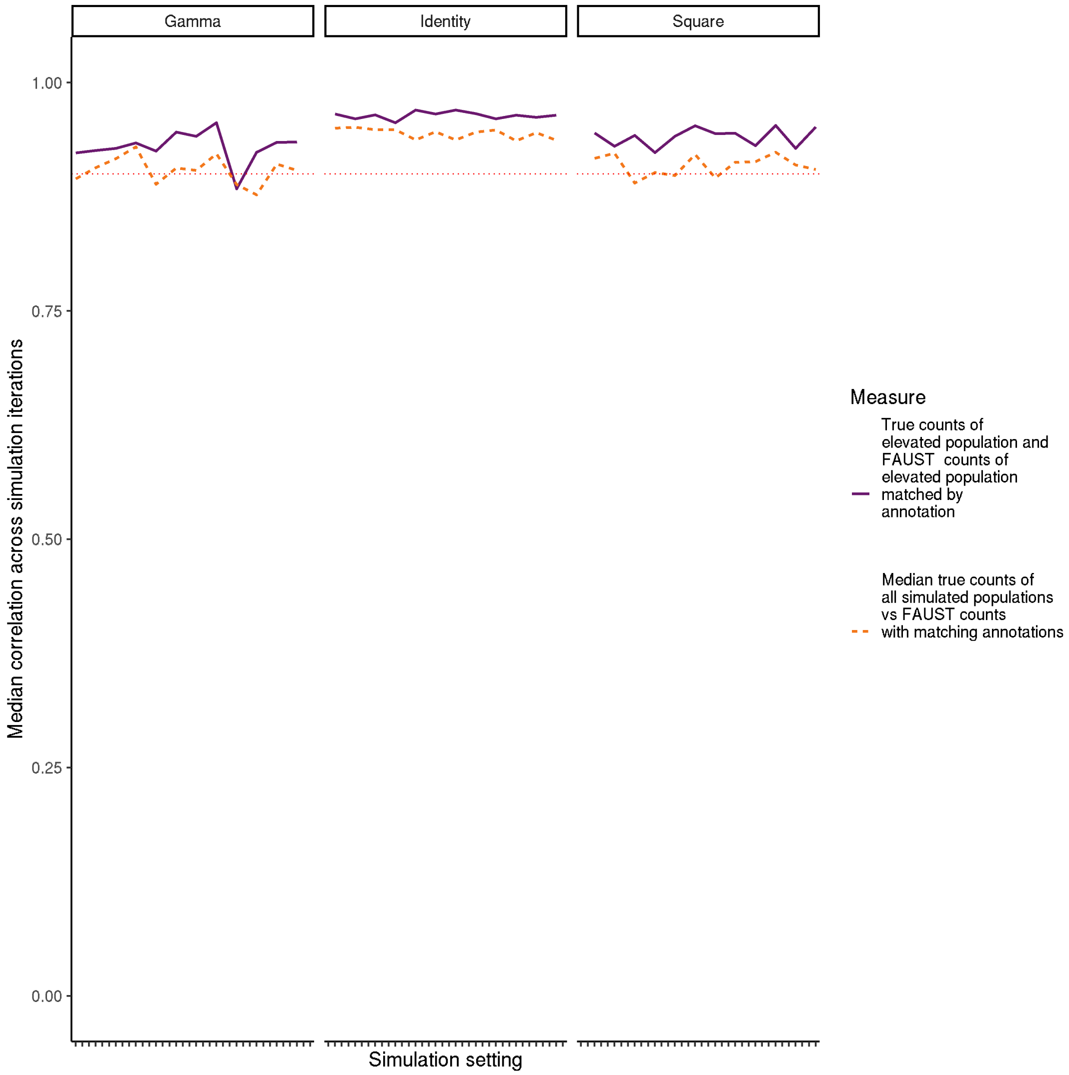


Figure S5: Dashed line: median correlation between all FAUST clusters and all simulated true populations over 25 simulation iterations. Solid line: median correlation between FAUST cluster with differential abundant population and simulated differentially abundant cluster over 25 simulation iterations. Correlations are determined only using annotations.

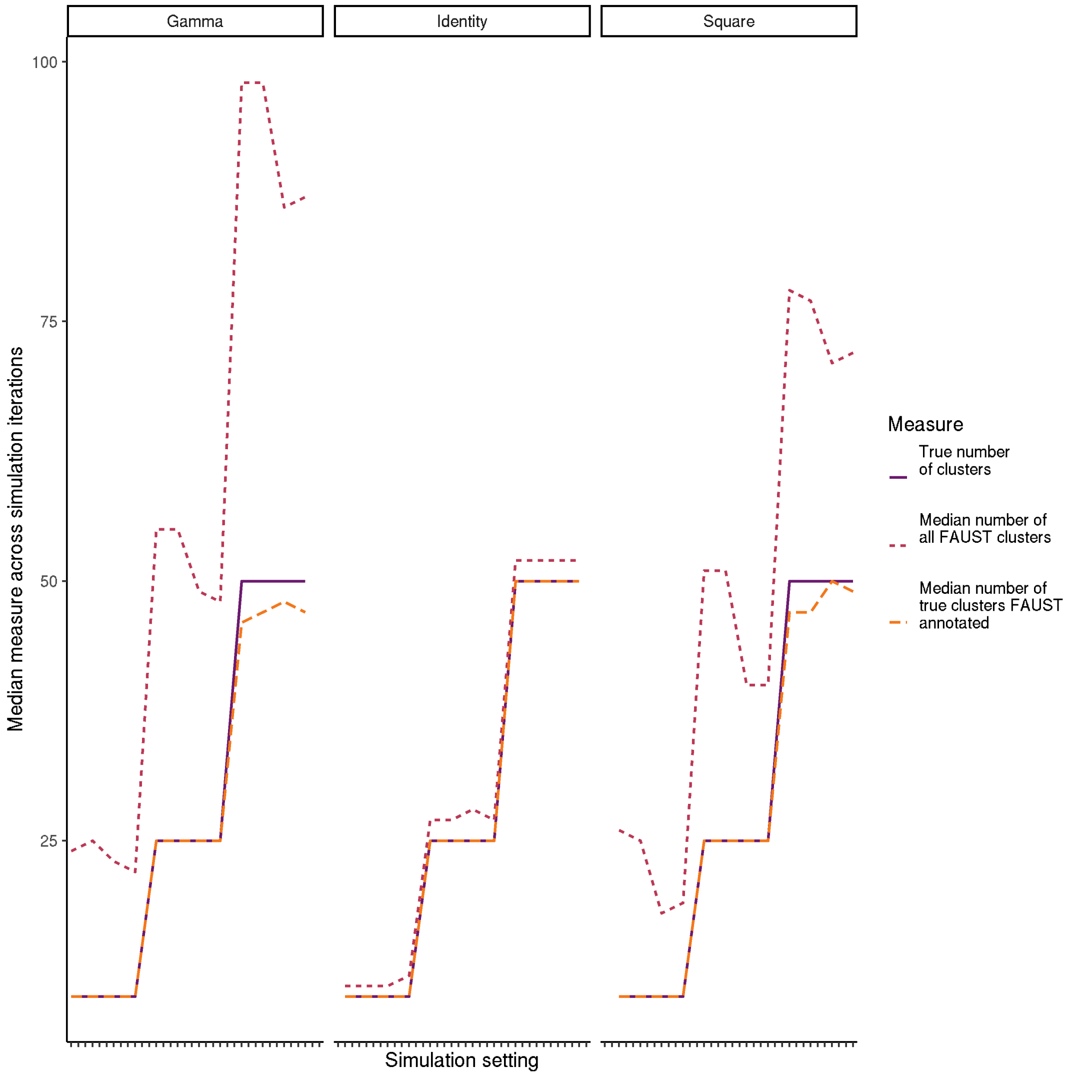


Figure S6: The true number of clusters by simulation setting is the solid purple line. The dashed orange line shows the median number of clusters matching the true annotations produced by FAUST across simulation settings over 25 simulation iterations. The dot-dashed red line show the median number of total annotated clusters produced by FAUST across simulation settings over 25 simulation iterations.

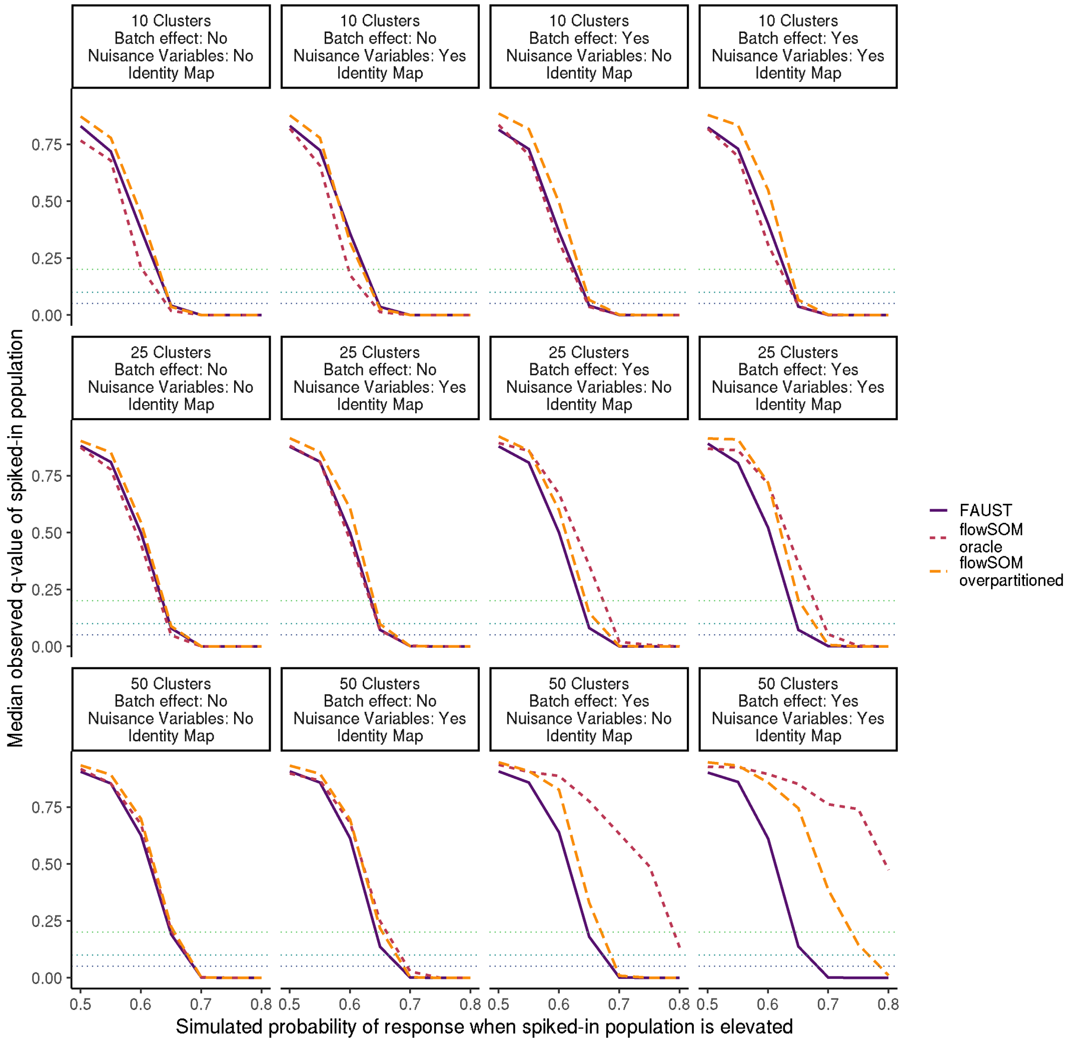


Figure S7: In each simulation, a differentially abundant sub-population is always simulated: 50 subjects have increased abundance relative to the other 50 subjects. For subjects with increased abundance, a stochastic response to therapy is then 50 times, with the response rate for subjects with increased abundance varying along the x-axis. Median FDR-adjusted p-value of FAUST cluster annotated with the differentially abundant population, and median FDR-adjusted for FlowSOM clusters identified as the true clusters are reported across 25 iterations. This plot reports performance when data are generated from a multivariate normal mixture, with the exact simulation settings reported in the facet label.

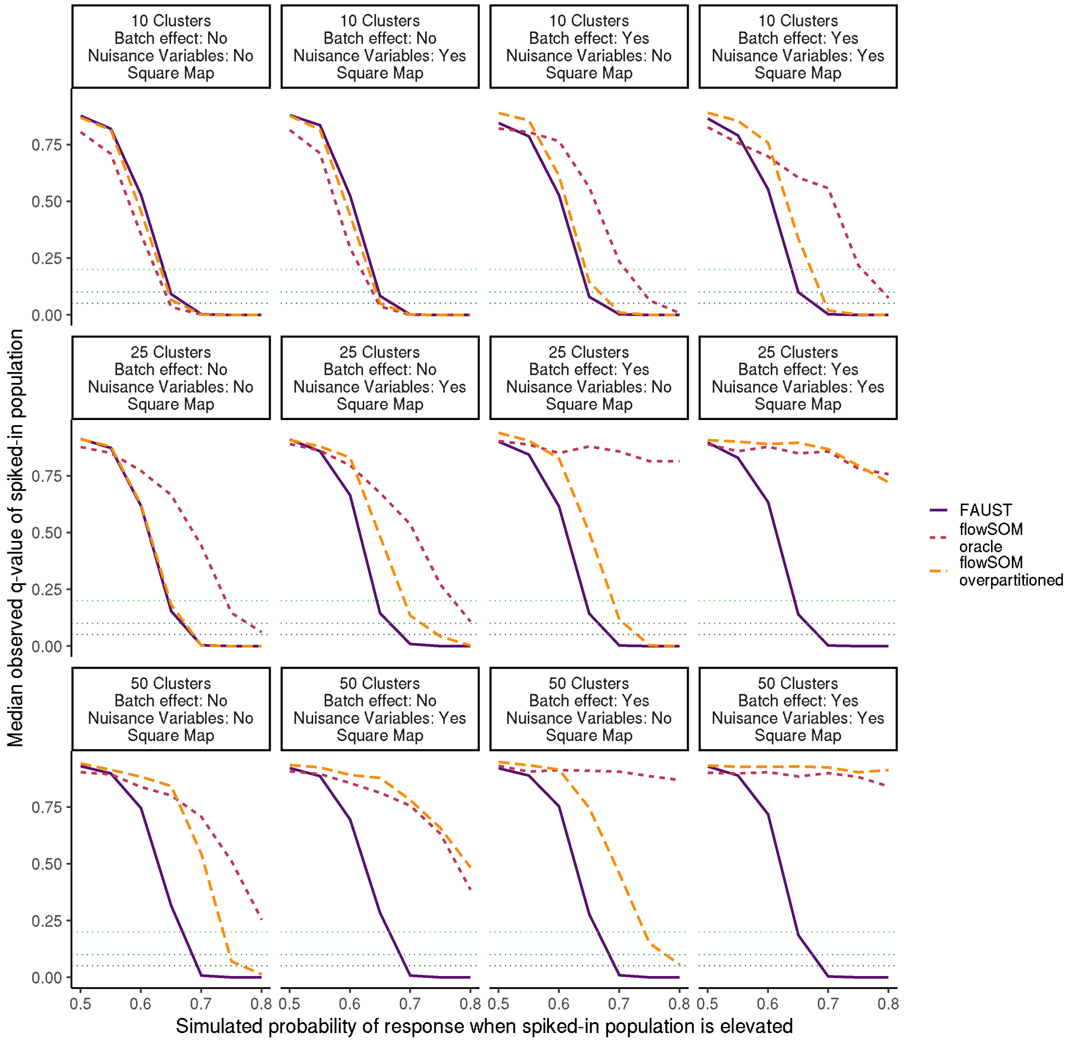


Figure S8: In each simulation, a differentially abundant sub-population is always simulated: 50 subjects have increased abundance relative to the other 50 subjects. For subjects with increased abundance, a stochastic response to therapy is then 50 times, with the response rate for subjects with increased abundance varying along the x-axis. Median FDR-adjusted p-value of FAUST cluster annotated with the differentially abundant population, and median FDR-adjusted for FlowSOM clusters identified as the true clusters are reported across 25 iterations. This plot reports performance when data are transformed by the coordinate map $f(x) = x^2$, with the exact simulation settings reported in the facet label.

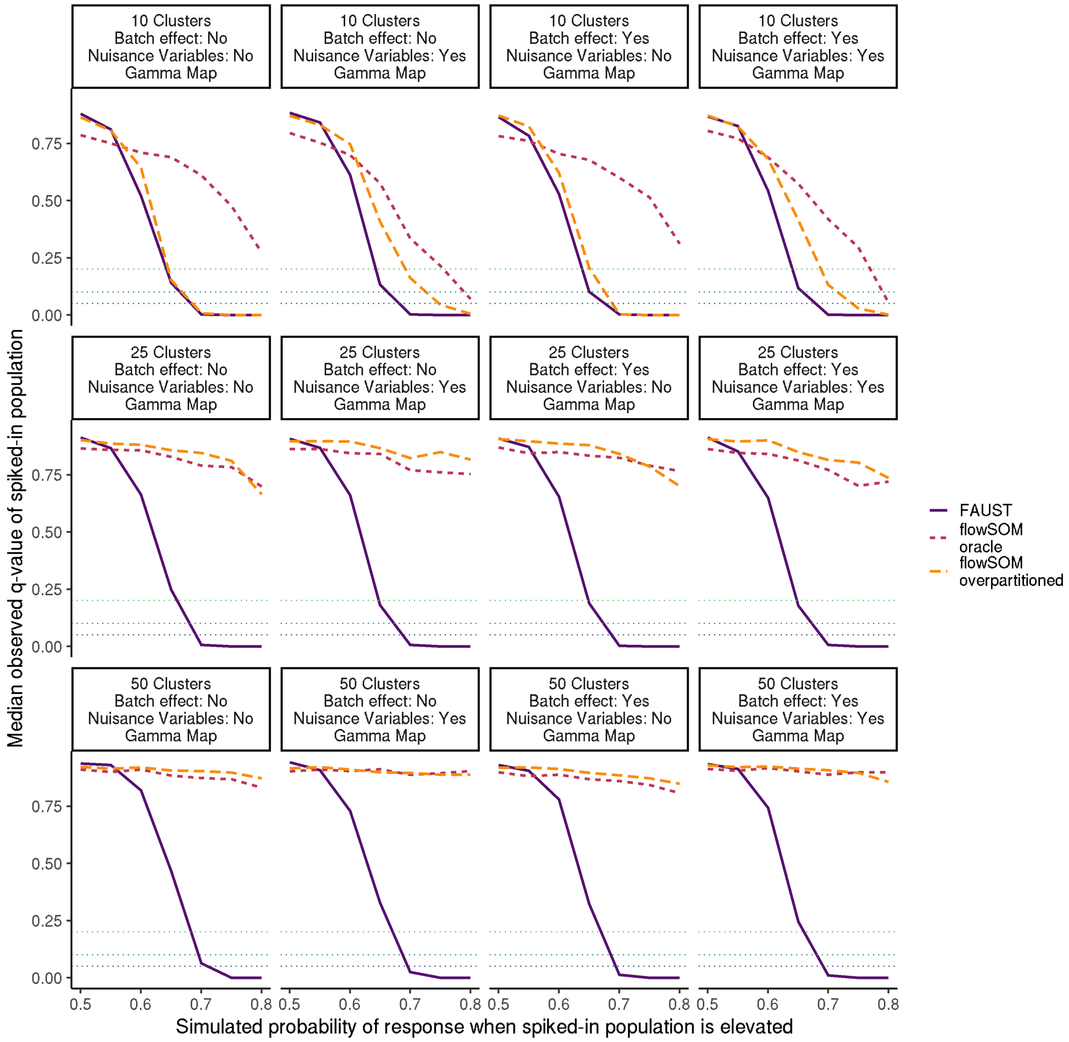


Figure S9: In each simulation, a differentially abundant sub-population is always simulated: 50 subjects have increased abundance relative to the other 50 subjects. For subjects with increased abundance, a stochastic response to therapy is then 50 times, with the response rate for subjects with increased abundance varying along the x-axis. Median FDR-adjusted p-value of FAUST cluster annotated with the differentially abundant population, and median FDR-adjusted for FlowSOM clusters identified as the true clusters are reported across 25 iterations. This plot reports performance when data are transformed by the coordinate map $g(x) = \Gamma(1 + |x/4|)$, with the exact simulation settings reported in the facet label.
